# Neutralizing antibodies induced by first-generation gp41-stabilized HIV-1 envelope trimers and nanoparticles

**DOI:** 10.1101/2020.12.02.408328

**Authors:** Sonu Kumar, Xiaohe Lin, Timothy Ngo, Benjamin Shapero, Cindy Sou, Joel D. Allen, Jeffrey Copps, Lei Zhang, Gabriel Ozorowski, Linling He, Max Crispin, Andrew B. Ward, Ian A. Wilson, Jiang Zhu

## Abstract

Antigen-specific B-cell sorting and next-generation sequencing (NGS) were combined to isolate HIV-1 neutralizing antibodies (NAbs) from mice and rabbits immunized with BG505 trimers and nanoparticles. Three mouse NAbs potently neutralize BG505.T332N and recognize a glycan epitope centered at the C3/V4 region, as revealed by electron microscopy (EM), x-ray crystallography, and epitope mapping. Three potent NAbs were sorted from rabbit B cells that target glycan holes on the BG505 envelope glycoprotein (Env) and account for a significant portion of autologous NAb response. We then determined a 3.4Å-resolution crystal structure for the clade C transmitted/founder Du172.17 Env with a redesigned heptad repeat 1 (HR1) bend. This clade C Env, as a soluble trimer and attached to a ferritin nanoparticle, along with a clade A Q482-d12 Env trimer, elicited distinct NAb responses in rabbits. Our study demonstrates that nanoparticles presenting gp41-stabilized trimers can induce potent NAb responses in mice and rabbits with Env-dependent breadth.

**TEASER:** Mouse and rabbit NAbs elicited by gp41-stabilized trimers and nanoparticles neutralize autologous HIV-1 by targeting different epitopes

## INTRODUCTION

The envelope glycoprotein (Env) on HIV-1 virions mediates cell entry and is the target of broadly neutralizing antibodies (bNAbs) (*1*). Diverse bNAb families have been identified from HIV-1 infected individuals. Structural characterization of these human bNAbs in complex with Env proteins has defined multiple sites of HIV-1 vulnerability such as the CD4 binding site (CD4bs), quaternary V1/V2 glycan site, N332-oligomannose patch, silent face, gp120-gp41 interface, fusion peptide (FP), and membrane-proximal external region (MPER) (*2–4*). These bNAbs often possess unusual sequence characteristics acquired during extensive virus-host coevolution. As a result, the targets of bNAbs differ substantially from the strain-specific epitopes recognized by autologous NAbs early in human infection (*5–8*). Information on both of these types of antibodies, and tracing them back to their unmutated common ancestors (UCAs) and early intermediates, are valuable tools in guiding rational design of vaccine immunogens (*9–11*).

Soluble native-like Env trimers have emerged as a promising platform for HIV-1 vaccine design (*12, 13*). As the leading design platform, SOSIP trimers have been created and characterized for diverse HIV-1 subtypes and strains (*14–17*), followed by native flexibly linked (NFL) (*18*) and uncleaved prefusion optimized (UFO) trimers (*19*). Env structures from x-ray crystallography and cryo-electron microscopy (cryo-EM) provided a rational basis for improving trimer design (*20–23*). While mutations aiming to increase Env stability and immunogenicity were extensively tested in the context of SOSIP and NFL trimers (*24–36*), the causes of Env metastability were probed on the basis of the UFO trimer (*19, 37*). To further enhance immune recognition, these three trimer designs have been displayed on nanoparticles (NPs) of diverse chemical nature such as protein, lipid, and iron oxide (*30, 37–44*). Various animal models such as mouse, rabbit, and nonhuman primate (NHP) have been used to assess immunogenicity of HIV-1 Env in soluble or particulate forms. In wildtype mice, BG505 SOSIP.664 Env trimer failed to elicit a detectable tier 2 NAb response, but showed robust non-neutralizing binding titers towards a “neoepitope” at its base (*45*). However, a tier 2 NAb response was observed for a 60-mer presenting 20 gp41-stabilized BG505 trimers (*37*). Using various vaccine design strategies, mouse antibodies were elicited to the N332-glycan supersite (non-neutralizing) (*46*) and fusion peptide (weak but broad) (*47*). A germline-targeting strategy proved to be successful in engineered mice with knocked-in genes corresponding to bNAbs and precursors (*48–50*). In contrast to the challenges of NAb elicitation in mice, potent and sometimes broad tier 2 NAb responses have been reported in *in vivo* studies where rabbits were immunized with diverse trimers and trimer-presenting NPs (*17, 27-30, 35, 51-53*). However, epitope mapping identified that specific glycan holes on the HIV-1 Env dominated the autologous NAb response in rabbits (*54–61*). HIV-1 immunogens in trimeric and particulate forms have also been assessed in NHPs, where they were found to elicit consistent autologous but sparse cross-subtype NAb responses (*17, 42, 51, 52, 59, 62-65*). The C3/465 epitope was identified as a major target of NAb responses in macaques induced by the BG505 SOSIP.664 trimer (*59, 66*). Overall, recognition of Env by mouse and NHP NAbs is substantially less understood compared to rabbit NAbs. In addition, the effect of both HR1 and gp41 stabilization, which is the core of the UFO trimer design (*19, 37*), on NAb elicitation and epitope targeting in wildtype animal models has not been as well characterized compared to SOSIP and NFL trimers.

Previously, we designed gp41-stabilized trimers and NPs and assessed their NAb responses in mice and rabbits (*37*). In this particular study, we set out to characterize mouse and rabbit NAbs induced by these immunogens in greater detail. First, we identified tier 2 mouse NAbs elicited by an HR1-redesigned BG505 trimer presented on a 60-meric I3-01 NP. A potent NAb, M4H2K1, was identified by pairing representative heavy and light chains obtained from next-generation sequencing (NGS) analysis of Env-specific splenic B cells, with two somatically related NAbs isolated by single B-cell sorting and antibody cloning. Negative-stain EM (nsEM) analysis showed that M4H2K1 recognized the C3/V4 region of the native-like BG505 Env. The crystal structure of M4H2K1 bound to a BG505 gp120 core at 4.3Å resolution revealed key antibody interactions with the C2/C3/V4/V5 epitope, which were confirmed in TZM-bl neutralization assays against a panel of BG505.T332N mutant viruses. A less potent NAb from a different mouse (M1), M1H2K1, was also identified, which likely targets the same epitope. We then performed single B-cell sorting and NGS for one rabbit immunized with an HR1-redesigned BG505 trimer and another with a ferritin NP presenting this trimer (*37*). Three representative rabbit NAbs were tested against a panel of glycan hole variants of BG505.T332N and found to target the glycan holes at 241/289 and 465. Further analysis of plasma neutralization confirmed that these glycan holes accounted for a large portion of the polyclonal antibody response, suggesting that ferritin display cannot broaden the rabbit NAb response induced by soluble BG505 Env. Lastly, we determined a 3.4 Å-resolution crystal structure for an HR1-redesigned trimer derived from the Env of a clade C transmitted founder (T/F) virus, Du172.17. In rabbits, the Du172.17 trimer and ferritin NP induced modest cross-clade NAb responses, whereas the UFO-BG trimer derived from a clade A T/F Q842-d12 Env exhibited a narrow NAb response. Our study thus confirmed that self-assembling protein NPs presenting gp41-stabilized trimers are capable of inducing potent tier-2 NAbs in mice and rabbits, in addition to structural and functional evaluation of the newly designed T/F Env trimers.

## RESULTS

### Mouse NAbs isolated by Env-specific B cell sorting and antibody NGS

Previously, we reported tier 2 NAb response in mouse immunization with NPs presenting an HR1-redesigned BG505 trimer (*37*). In separate studies, we used a BG505 trimer probe bearing this redesigned HR1 to identify early intermediates of the PGT121 lineage from a phage antibody library (*67*) and two N332-directed bNAbs from peripheral blood mononuclear cells (PBMCs) of an HIV-1-infected Chinese donor (*68*). Here, we used this BG505 trimer probe in two strategies to assist in NAb identification from mouse splenic B cells (**Fig. 1**). One strategy focused on NGS analysis of bulk-sorted B cells and the other involved on single-cell sorting and antibody cloning. The I3-01 NP group (*37*), in which two mice (M1 and M4) developed a robust tier 2 NAb response, was selected to characterize the mouse NAbs at the monoclonal level.

**Fig. 1.**
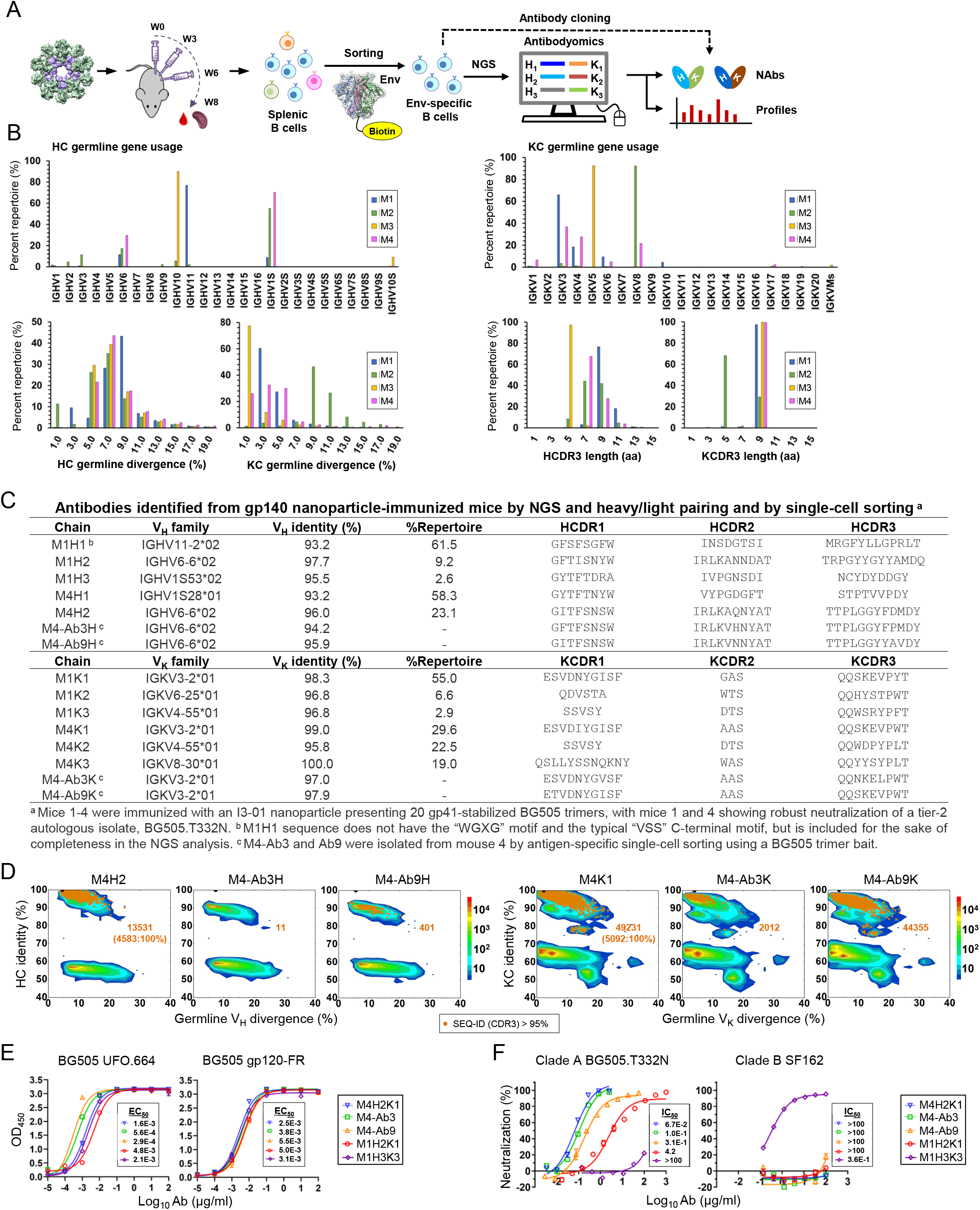
Tier 2 neutralizing antibodies isolated from gp140 nanoparticle-immunized mice. (**A**) Schematic representation depicting mouse immunization with BG505 gp140.664.R1-PADRE-I3-01 nanoparticle and antibody isolation from mouse splenic B cells using two approaches: Env-specific bulk B cell sorting followed by next-generation sequencing (NGS) and Env-specific single B cell sorting combined with antibody cloning. **(B)** Quantitative B cell repertoire profiles derived from the NGS analysis of Env-specific splenic B cells from four mice in the I3-01 group, including germline gene usage, degree of somatic hypermutation (SHM), and CDR3 loop length. **(C)** Characteristics of antibody heavy and κ-light chains (HC and KC) identified from clustering analysis of mouse NGS data and from single B cell sorting and antibody cloning. **(D)** Divergence-identity analysis of murine NAbs in the context of Env-specific splenic B cells from mouse #4 (M4). HCs and KCs are plotted as a function of sequence identity to the template and sequence divergence from putative germline genes. Color-coding denotes sequence density. The template and sequences identified based on the CDR3 identity of 95% or greater to the template are shown as black and orange dots on the plots, with the number of related sequences labeled accordingly. **(E)** ELISA binding of mouse NAbs to the BG505 UFO.664 trimer and BG505 gp120-ferritin (FR) nanoparticle probe with EC50 values labeled next to the binding curves. **(F)** Percent neutralization of mouse NAbs against autologous tier 2 clade A BG505.TN332N and heterologous tier 1 clade B SF162 pseudoviruses with IC50 values labeled next to the neutralization curves. Five mouse NAbs including M4H2K1 (blue), M4-Ab3 (green), M4-Ab9 (orange), M1H2K1 (red), and M1H3K3 (purple) are shown in **(E)** and **(F)**.

We first isolated mouse NAbs through NGS analysis of bulk-sorted Env-specific B cells and random pairing of consensus heavy and light chains (**Fig. 1**). This approach was devised based on the hypothesis that the small number of vaccine-induced B cell lineages will enable frequency-based identification of functional antibodies. In bulk sorting, 87∼1064 BG505 Env-specific B cells were obtained from the four mice studied (**fig. S1A**). Unbiased mouse antibody heavy and κ-light chain (HC and KC) libraries were constructed and sequenced on an Ion S5 platform, which yielded up to 1.22 million raw reads (**fig. S1B**). The antibody NGS data were processed using a mouse antibodyomics pipeline (*69*) to remove low-quality and incomplete reads (**fig. S1B**). Quantitative profiles of Env-specific B cell populations were determined for each mouse in the I3-01 NP group, revealing distinct patterns (**Fig. 1B**). Diverse antibody variable (VH and VK) genes were activated in response to Env immunization with some overlap observed for the two mice (M1 and M4) that developed a tier 2 autologous NAb response (*37*). While IGHV6 and IGHV1S were used by Env-specific antibodies from both M1 and M4, 77% of M1 HCs were derived from IGHV11 and 70% of M4 HCs from IGHV1S. A similar pattern was observed for the VK distribution, with overlap on the IGKV3, IGKV4, and IGKV6 genes. In terms of somatic hypermutation (SHM), a consistent VH distribution was observed for four mice that peaked at the 7-9% nucleotide (nt) difference from assigned germline genes. In contrast, four mice exhibited significant differences in their VK SHM distributions, with M2 and M3 showing the largest difference in average SHM of 10.0% and 1.6%, respectively. In terms of complementarity-determining region 3 (CDR3) length, M2 and M3 also appeared to show a notable difference from M1 and M4 by using predominantly 5-aa KCDR3 and HCDR3 loops, respectively. Nonetheless, a CDR3-based clustering algorithm (*67*) was used to calculate consensus HCs and KCs from the M1 and M4 NGS data (**Fig. 1C** and **fig. S1C**), because IgG purified from these two mice neutralized BG505.T332N (*37*). Interestingly, M1H2 and M4H2, both from the second largest sequence family, were of the IGHV6-6*02 origin, whereas M1K1 and M4K1 shared the IGKV3-2*01 germline gene (**Fig. 1C**). These consensus HCs and KCs were synthesized to reconstitute mouse antibodies. To further enrich the antibody pool, we performed single B-cell sorting on M4 splenic B cells using the same BG505 probe. The natively paired HCs and KCs of two monoclonal antibodies (mAbs), M4-Ab3 and M4-Ab9, were derived from IGHV6-6*02 and IGKV3-2*01, suggesting that they might be somatically related to M4H2 and M4K1, respectively (**Fig. 1C** and **fig. S1D**). Two-dimensional (2D) divergence/identity analysis (*68, 70*) was performed to compare the prevalence of these mouse mAbs in the NGS-derived antibody repertoire (**Fig. 1D**). Using an HCDR3 identity cutoff of 95%, 13531, 11, and 401 sequences were related to M4H2, M4-Ab3 HC, and M4-Ab9 HC, respectively. Based on the same KCDR3 identity cutoff, 49231, 2012, and 44355 sequences were somatically related to M4K1, M4-Ab3 KC, and M4-Ab9 KC, respectively. Of note, a significant portion of somatically related HCs and KCs were identical to M4H2 (33.9%) and M4K1 (10.3%), respectively, suggesting that these two consensus sequences represent native antibody chains used by Env-specific B cells from M4. Taken together, a panel of mAbs were identified from two NAb-producing mice in our previous study (*37*).

We characterized the binding of these mouse mAbs to a panel of Env antigens by enzyme-linked immunosorbent assay (ELISA) (**Fig. 1E** and **fig. S2A**). When the BG505 UFO.664 trimer was used as a coating antigen, three mAbs from M4, including NGS-derived M4H2K1 and single-cell sorting-derived M4-Ab3 and M4-Ab9, and two NGS-derived mAbs from M1, M1H2K1 and M1H3K3, bound to this native-like Env trimer with up to 16.6-fold difference in the half maximal effective concentration (EC50) value (**Fig. 1E**, left). Among the three M4 mAbs, the two single-cell sorting-derived mAbs bound to BG505 UFO.664 Env with 2.9 and 5.5-fold lower EC50 values than M4H2K1. Other NGS-derived HC-KC pairs showed low or no trimer binding (**fig. S2A**, top). We examined the epitope specificity of trimer-binding mAbs by testing four NP probes derived from ferritin (FR), including BG505 gp120-FR (*41*), an N332-FR termed 1GUT_A_ES-5GS-FR (*69*), a BG505 V1V2-FR (*41*), and a newly developed FP-5GS-FR. In ELISA, all five mAbs bound to BG505 gp120-FR with comparable EC50 values (**Fig. 1E**, right), but failed to show any detectable binding to the N332 supersite, V1V2 apex, and FP epitope in the context of the probes (**fig. S2A**, bottom), suggesting that they may recognize a different epitope in gp120.

We characterized the neutralizing activity of these mouse mAbs in TZM-bl assays (**Fig. 1F**; **fig. S2B** and **S2C**). All trimer-binding mAbs, except for M1H3K3, neutralized the autologous tier 2 BG505.T332N with up to 63-fold difference in the half maximal inhibitory concentration (IC50) value (**Fig. 1F**, left). The NGS-derived mAb from M4 (M4H2K1) appeared to be the most potent neutralizer with an IC50 value of 0.067 μg/ml, which is 2- to 5-fold higher IC50 than bNAbs PGT121 (0.029 μg/ml) and PGT128 (0.013 μg/ml), respectively (*68*). In terms of potency, this mouse NAb was comparable to those C3/V5-specific autologous NAbs isolated from NHPs, which showed a median IC50 value of 0.06 μg/ml (*66*). Despite stronger Env binding, the sorting-derived M4-Ab3 and M4-Ab9 neutralized BG505.T332N less effectively than M4H2K1 with up to 4.6-fold higher IC50 values. In comparison, other NGS-derived HC-KC pairs only exhibited low levels of autologous neutralization at high immunoglobulin G (IgG) concentrations (**fig. S2B**, top panel). When tested against a tier 1 clade B virus SF162, M1H3K3 yielded an IC50 value of 0.36 μg/ml, whereas the other four autologous tier 2 NAbs did not exhibit any reactivity with SF162 (**Fig. 1F**, right; **fig. S2B**, middle panel). All mouse mAbs did not neutralize the murine leukemia virus (MLV) Env-pseudotyped virus except for M4-Ab9, which showed non-specific signals at high IgG concentrations (**fig. S2B**, bottom panel). Lastly, we assessed the neutralizing activity of these mouse mAbs against a global panel of 12 isolates (*71*). Using MLV as a negative control in TZM-bl assays, M1H3K3 from M1, but not any of the M4 mAbs, modestly neutralized two heterologous HIV-1 isolates, clade A/E pCNE8 and clade G pX1632 (**fig. S2C**).

In brief, a panel of mAbs was identified from mice immunized with an I3-01 60-mer using a BG505 Env probe in two B cell sorting strategies followed by NGS and bioinformatics analysis. Functional evaluation confirmed that M4H2K1 is an autologous tier 2 NAb with high potency, whereas M1H3K3 is less potent but cross-reactive with other tier 2 isolates. These two murine NAbs neutralized HIV-1 by targeting as yet unidentified epitopes in the gp120 subunit.

### Autologous tier 2 mouse NAb M4H2K1 binds laterally to the BG505 Env trimer

Previously, a modified BG505 SOSIP.664 trimer (RC1) displayed on virus-like particles (VLPs) expanded mouse germinal center (GC) B cells specific to the V3 glycan patch (*46*). In a recent study, Ringe et al. reported that mice immunized with the soluble trimer generated autologous serum neutralizing response to the glycan hole at position 289 (*44*). Here, we combined negative-stain EM (nsEM) and x-ray crystallography to elucidate the mechanism of how M4H2K1, one of the most potent murine NAbs identified thus far, interacts with HIV-1 Env.

We first performed EM analysis to visualize where the mouse NAb M4H2K1 binds on the BG505 Env trimer. We produced the antigen-binding fragment (Fab) of M4H2K1 and incubated with the BG505 UFO.664 trimer to form a complex, which was subjected to single-particle nsEM analysis (**fig. S3A**). The three-dimensional (3D) reconstruction showed that the major species of this complex was Env trimers each bound to three M4H2K1 Fabs, with each Fab approaching the Env laterally (**Fig. 2A**, leftmost). After fitting a crystal structure of BG505 SOSIP.664 [PDB ID: 4TVP (*72*)] into the EM electron density, M4H2K1 was found to interact with an epitope that lies approximately in the gp120 C2/C3/V4/V5 region. To determine how much the M4H2K1 epitope overlaps with the neighboring bNAb epitopes, EM maps containing Fabs of four representative bNAbs, VRC01 (*73*), 2G12 (*74*), PGT135 (*75*), and 8ANC195 (*76*), were aligned to the M4H2K1 EM complex (**Fig. 2A**, right four). Our analysis revealed that M4H2K1 and VRC01 (but not other NAbs) would “clash” in their Env-bound mode as indicated by slightly overlapping EM densities, suggesting that the M4H2K1 epitope is in proximity to the CD4bs targeted by VRC01.

**Fig. 2.**
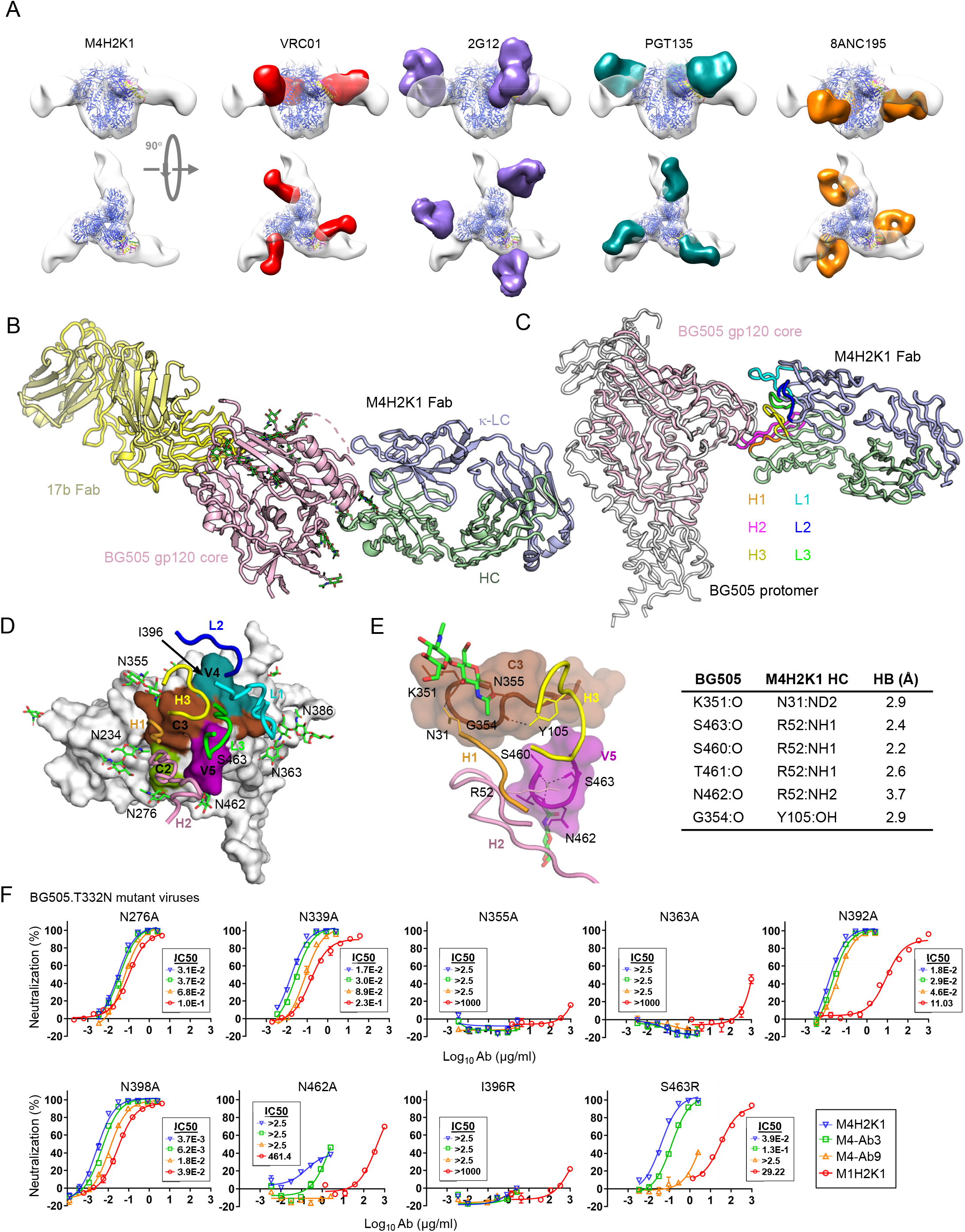
Structural epitope mapping of M4H2K1 on HIV-1 Env. **(A)** 3D EM reconstruction of M4H2K1 Fab/BG505 UFO.664 complex. The crystal structure of BG505 SOSIP.664 trimer (PDB ID: 4TVP) is docked into the trimer EM density and displayed in blue ribbons. Comparison of the mode of M4H2K1 Fab binding to BG505 UFO.664 trimer with four bNAbs, VRC01 (red; EMD-6252), 2G12 (purple; EMD-5982), PGT135 (cyan; EMD-2331), and 8ANC195 (orange; EMD-2625). **(B)** Crystal structure of BG505 gp120 core (pink) in complex with Fabs 17b (yellow) and M4H2K1 at 4.3Å resolution. **(C)** Side view of the crystal structure of the M4H2K1 Fab-BG505 gp120 core (light pink) complex superimposed onto one protomer of BG505 gp140 (grey) (PDB ID: 5CEZ). The M4H2K1 Fab is shown with the HCDR loops labeled and colored [H1 (orange), H2 (pink), and H3 (yellow)] and LCDR loops [L1 (cyan), L2 (blue), and L3 (green)]. **(D)** Epitope of M4H2K1 Fab mapped onto the BG505 gp120 core shown in surface representation and defined as two residues containing an atom within 4.0Å of each other (C2: green; C3: brown; V4: cyan; and V5: pink). The M4H2K1 Fab is shown with HCDR and LCDR loops labeled and colored accordingly. **(E)** Left: hydrogen bonds are shown between HCDR loops (H3:yellow; H2:pink; and H1:orange) and gp120 core (C3:brown; and V5:pink) and right, table showing residues involved in hydrogen-bond (HB) interaction and distance were measured in Å. **(F)** Percent neutralization of mouse NAbs against nine BG505 mutant pseudoviruses with IC50 values labeled next to the neutralization curves. NAbs M4H2K1 (blue), M4-Ab3 (green), M4-Ab9 (orange), and M1H2K1 (red) are shown.

We then applied x-ray crystallography to further understand the molecular interactions of M4H2K1 with BG505 Env. To this end, we first obtained the crystal structure of M4H2K1 Fab at 1.50 Å resolution (**fig. S3B**). In this structure, HCDR3 (10 aa) is sandwiched between HCDR1 and KCDR2, while KCDR3 (9 aa) fits between KCDR1 and HCDR2 (**fig. S3B**). To gain more atomic details of the M4H2K1-Env interaction, we generated a gp120 core from the clade A BG505 Env to complex with Fab M4H2K1 as well as Fab 17b (*77*) to aid in crystallization. A crystal structure of this complex was determined at 4.30 Å resolution in an orthorhombic (P21212) crystal lattice. The structure showed that M4H2K1 Fab bound to the BG505 gp120 core by targeting the C2/C3/V4/V5 region (**Fig. 2B** and **2D**). We then superimposed the M4H2K1 Fab-gp120 core complex onto a protomer of the BG505 SOSIP.664 trimer (PDB ID: 5CEZ) (*78*), which defined the orientation of M4H2K1 Fab HC and KC relative to BG505 Env in a lateral binding mode (**Fig. 2C**). The extended HCDR2 (19 aa) and KCDR1 (15 aa) engage the trimer in a pincer-like grasp (magenta and cyan loops in **Fig. 2C**). A total of 865 Å^2^ of the Fab is buried on the BG505 gp120 core surface, where HC and KC contribute to 70% and 30% of the Fab-buried surface area (BSA), respectively (**Fig. 2D** and **fig. S3C**). The tip of the HCDR2 loop is deeply buried (348 Å^2^) inside the pocket formed by multiple parts of C2/C3/V5 (**Fig. 2D** and **fig. S3C**) and makes most contact with Env, followed by KCDR1 with the next largest BSA of 193 Å^2^. In addition, the other CDRs (BSA; H3: 117 Å^2^, H1: 130 Å^2^, L3: 67 Å^2^) and HC framework region 1 (BSA; HFR1: 9 Å^2^) are buried into the gp120 core surface except for KCDR2, which has no BSA (**fig. S3C**). This analysis highlights the importance of a long HCDR2 (19 aa) in anchoring M4H2K1 Fab to Env in a lateral orientation. Despite the moderate resolution, interactions at the interface of BG505 gp120 core and M4H2K1 Fab could be observed with little ambiguity. A hydrogen bond network in the interface appears to be formed by HCDR1 (N31), HCDR2 (R52), and HCDR3 (Y105) in M4H2K1 Fab and T278, R350, K351, N356, S460, T461, N462, and S463 in the BG505 gp120 core (**Fig. 2E**). To determine the regions involved in steric clashes between M4H2K1 and VRC01 Fabs upon EM fitting (**Fig. 2A**, panel 2 to the left), we superimposed the crystal structure of M4H2K1 Fab-BG505 gp120 core complex onto VRC01-bound BG505 F14 SOSIP trimer (PDB ID: 6V8X (*79*)). The overlap between the 19aa-long HCDR2 loop of M4H2K1 Fab and LCDR1 of VRC01 Fab would suggest competition between M4H2K1 and VRC01-class bNAbs for Env binding (**fig. S3D**). Due to similar steric hindrance, the IOMA-class bNAbs (*80*) could also compete with M4H2K1 for Env binding.

In a recent study, a group of NAbs was isolated from guinea pigs immunized with a BG505 SOSIP.664 trimer (*81*). Of these NAbs, CP506 Fab targets the C3/V4 region of HIV-1 Env and neutralizes BG505.T332N with an IC50 value of 0.1 μg/ml. To compare the angle of approach between M4H2K1 and CP506 Fabs, we first constructed a model of M4H2K1 Fab-bound BG505 Env trimer by superimposing our crystal structure of M4H2K1 Fab-BG505 gp120 core complex onto the BG505 SOSIP.664 trimer (PDB ID: 4TVP (*72*)). We then docked this model into the 3D reconstruction of CP506 Fab-BG505 SOSIP.664 trimer complex derived from the nsEM analysis (EMD-9003). A slight variation in angle of approach was observed (**fig. S3E**). In comparison with CP506, which has a 16-aa HCDR2 and an 8-aa HCDR3, M4H2K1 utilizes longer HCDR2 (19-aa) and HCDR3 (10-aa) to recognize HIV-1 Env. The crystal structure (**Fig. 2B**) suggests that glycans at N276, N339, N355, N363, and N462, as well as amino acids I396 and S463 (**Figs. 2D** and **2E**), may be involved in BG505 Env recognition by M4H2K1. Glycans N234 and N386 point sideways with no direct contact with M4H2K1. To verify these interactions, we created nine BG505.T332N variants, N276A, N339A, N355A, N363A, N462A, I396R and S463R, along with N392A and N398A, which are in proximity to the binding site, and tested their neutralization by four of newly identified mouse NAbs (**Fig. 2F**). While glycan knockouts (KO) at positions 276, 339, 392, and 398 and the S463R mutation exhibited little to modest effect, glycan KOs N355A, N363A, and N462A and the I396R mutation significantly reduced or completely abrogated neutralization by the mouse NAbs (**Fig. 2F**), suggesting that glycans at N355 and N363, as well as a contribution by glycan N462, along with I396 are critical for M4H2K1-Env interaction. In contrast, glycans at N339, N363 and N392 were critical for Env recognition by the guinea pig NAb, CP506 (*81*), which targets an overlapping C3/V4 epitope. Lastly, we examined the conservation of three critical residues (N355, N363, and I396) in 6,966 HIV-1 Env sequences (www.hig.lanl.gov/). A large fraction of isolates contain an NXT/S sequon at N355 (∼80%) and N462 (∼42%) as does BG505, whereas positions 363 and 396 are less conserved in the group M isolates, with 9% being Asn and 4.5% being Ile, respectively, leading to the autologous nature of NAb M4H2K1.

In summary, our structural analysis identified a critical epitope on BG505 Env that can be recognized by potent murine NAbs exemplified by M4H2K1. Compared to CP506, a guinea pig NAb targeting C3/V4 (*81*), M4H2K1 achieves its potency by interacting with an expanded Env surface area spanning C2/C3/V4/V5. A less potent NAb from another mouse, M1H2K1, exhibited similar sensitivity to a panel of BG505.T332N variants in TZM-bl neutralization assays (**Fig. 2F**), suggesting that this NAb may recognize a similar epitope to M4H2K1, albeit with differential effects of mutations at N363, N392, and S463.

### Stabilized BG505 trimer and ferritin nanoparticle elicit glycan hole NAbs in rabbits

Extensive studies of native-like trimers, particularly of the BG505 backbone, have revealed that the autologous NAb response in rabbits was mainly directed to “glycan holes” (*54–61*). Strategies intended to broaden the autologous NAb response in rabbit immunization with mixed SOSIP.664 trimers of two different clades and with the immune complex of a BG505 SOSIP.664 trimer and a glycan hole mAb proved not to be effective (*82, 83*). Previously, we immunized rabbits with a BG505 trimer containing a redesigned HR1 bend, termed gp140.664.R1, and an FR NP displaying this trimer (*37*) (**Fig. 3A**). The gp140.664.R1-FR NP elicited a more rapid autologous tier 2 NAb response than the soluble trimer (*37*). Here, we sought to isolate NAbs from previously immunized rabbits for functional evaluation, repertoire profiling, and epitope mapping.

**Fig. 3.**
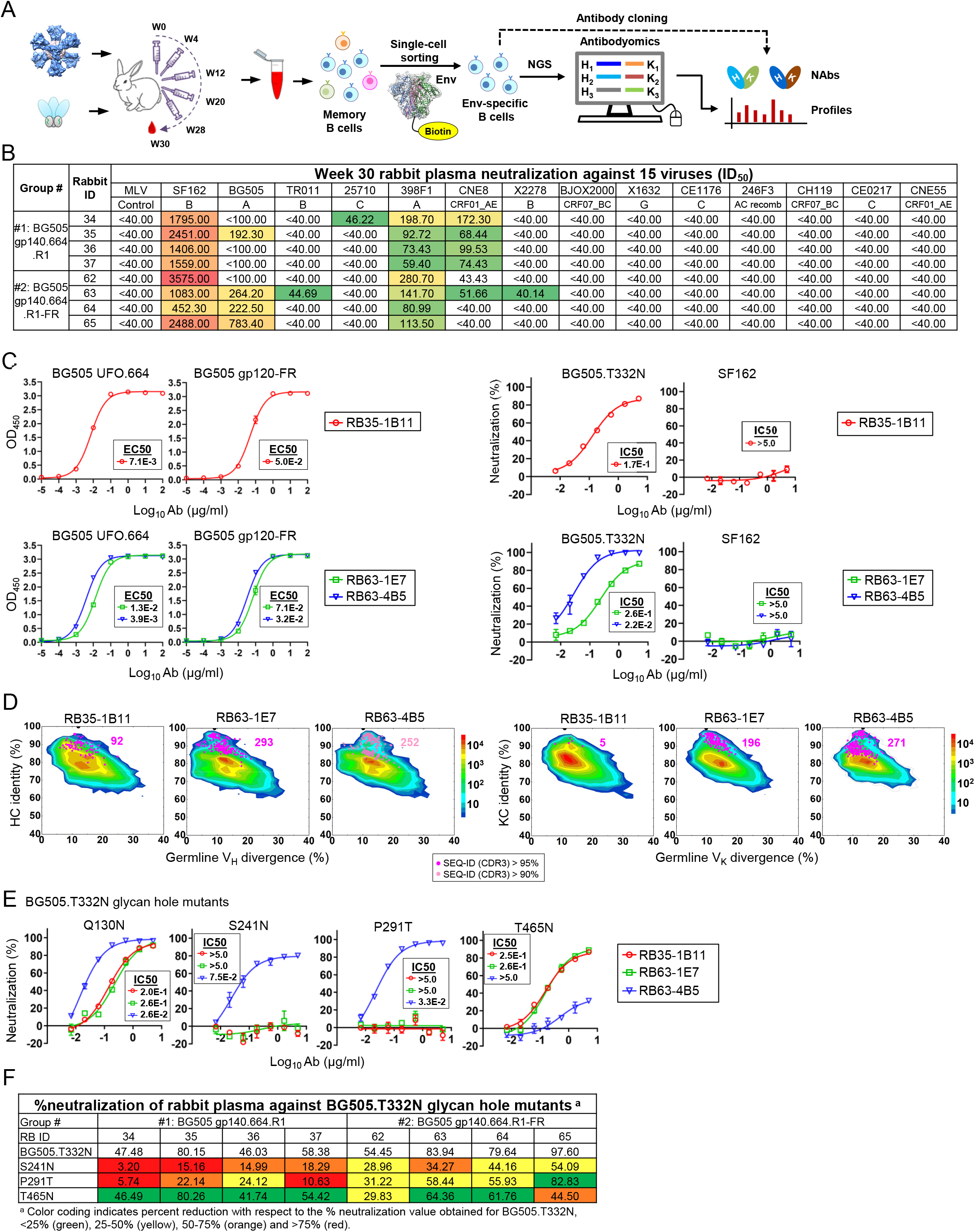
Tier-2 neutralizing antibodies isolated from rabbits immunized with HR1-redesgined BG505 gp140 trimer and its ferritin nanoparticle. (**A**) Schematic representation depicting rabbit immunization with the BG505-gp140.664.R1 trimer and ferritin (FR) nanoparticle and antibody isolation from rabbit PBMCs by Env-specific single B cell sorting coupled with antibody cloning. (**B**) Neutralization (measured by ID50 values) of week-30 plasma from two rabbit groups against autologous tier 2 clade A BG505.T332N, tier 1 clade B SF162, and a 12-virus global panel with MLV included as a negative control. Color coding indicates neutralization potency (red: potent; green neutralizing but not potent; no color: non-neutralizing). **(C)** Left: ELISA binding of rabbit NAbs to the BG505 UFO.664 trimer and BG505 gp120-FR nanoparticle probe with EC50 values labeled next to the ELISA curves; Right: Percent neutralization of rabbit NAbs against autologous tie 2 clade A BG505.TN332N and heterologous tier 1 clade B SF162 pseudoviruses with IC50 values labeled next to the neutralization curves. Antibodies were diluted to 10μg/ml and subjected to a 3-fold dilution series in the TZM-bl assay. **(D)** Divergence-identity analysis of rabbit mAbs in the context of Env-specific rabbit B cell repertoires from rabbits RB35 and RB63. HC and KC sequences are plotted as a function of sequence identity to the template and sequence divergence from putative germline genes. Color-coding denotes sequence density. Templates and sequences identified based on the CDR3 identity cutoffs of 95% and 90% are shown as pink and light pink dots on the plots with the number of sequences labeled accordingly. **(E)**. Percent neutralization of rabbit NAbs against four BG505 pseudovirus containing glycan hole mutations with IC50 values labeled next to the neutralization curves. NAbs RB35-1B11 (red), RB63-1E7 (green), and RB63-4B5 (blue) are shown in (**C**) and (**E**). **(F)** % neutralization of rabbit plasma against BG505.T332N and its three glycan hole mutants. Color coding in the table indicates percent reduction with respect to the % neutralization value obtained for BG505.T332N, <25% (green), 25-50% (yellow), 50-75% (orange) and >75% (red).

We first assessed rabbit plasma at the last time point (week 30) against autologous tier 2 clade A BG505.T332N, tier 1 clade B SF162, and a global panel of 12 diverse isolates, with MLV included as a negative control (**Fig. 3B**). Half maximal inhibitory dilutions (ID50) were calculated from percent neutralization upon fitting (**fig. S4**). Consistent with our previous finding (*37*), the FR NP, BG5050 gp140.664.R1-FR, elicited a more potent autologous tier 2 NAb response than the soluble trimer, BG505 gp140.664.R1, with an ID50 values of 222 to 783 for three of four rabbits, whereas only one of four rabbits in the trimer group yielded a detectable ID50 value (192) using a 100-fold starting dilution. While the week-30 plasma from both groups neutralized clade A p398F1, only the soluble trimer group showed consistent measurable neutralization against a clade A/E recombinant strain (pCNE8) using a 40-fold starting dilution. The week-30 rabbit plasma potently neutralized tier 1 SF162 without non-specific MLV reactivity. As another control, pre-immunization samples (-d10) were tested against the global panel in TZM-bl assays and exhibited a clean background (**fig. S4**).

We selected RB35 in the trimer group and RB63 in the FR group for antibody isolation. Using the biotinylated Avi-tagged HR1-redesigned BG505 trimer probe (*67, 68*), we isolated Env-specific single B cells from PBMCs. A panel of rabbit mAbs was reconstituted from cloned HCs and KCs using a previously reported protocol (*55*), producing 34 and 55 HC-KC pairs for RB35 and RB63, respectively (**fig. S5A**). A rapid functional screening based on antibody yield and BG505.T332N neutralization resulted in three hits, one from RB35, and two from RB63 (**fig. S5A**). Sequence analysis revealed diverse germline gene usage (**fig. S5B**): RB35-1B11 is derived from IGHV1S45*01 and IGKV1S36*01, while RB63-1E7 and RB63-4B5 use the same HC germline gene (IGHV1S40*01); their KCs are of IGKV1S10*01 and IGKV1S15*01 origin, respectively. In ELISA, the three rabbit NAbs were tested against BG505 UFO.664 trimer (*19*) and four epitope probes, including a BG505 gp120-FR (*41*), an N332-I3-01 NP termed 1GUT_A_ES-I3-01 (*37*), a trimeric scaffold (PDB: 1TD0) presenting ZM109 V1V2 (termed ZM109 V1V2-5GS-1TD0), and a FP-5GS-1TD0. As indicated by the EC50 values as well as ELISA curves, all three rabbit Nabs showed high affinity for the trimer and gp120 probes (**Fig. 3C**, left), but no detectable binding to the N332, V1V2, and FP probes (**fig. S5C**), suggesting that they recognize other epitopes in gp120 of BG505 Env. All three NAbs neutralized the autologous tier 2 clade A BG505.T332N, but not the tier 1 clade B SF162 and negative control, MLV (**Fig. 3C**, right; **fig. S5D**). Of note, the most potent rabbit NAb, RB63-4B5, yielded an IC50 value of 0.022 μg/ml, which is ∼3-fold and 5-fold lower than the IC50 values of mouse NAb M4H2K1 and previously identified glycan hole NAbs (*55*), respectively. Lastly, all three rabbit NAbs showed negligible neutralization against the 12-virus global panel (**fig. S5E**). Six non-NAbs, three from each rabbit, were confirmed to be non-reactive with BG505.T332N in TZM-bl assays (**fig. S5F**).

To examine B cell lineages associated with these three NAbs, we applied NGS to analyze the Env-specific B cells from RB35 and RB63. Using the HR1-redesigned trimer probe, we sorted 363 and 370 Env-specific B cells from RB35 and RB63, respectively (**fig. S6A**). Unbiased rabbit antibody HC and KC libraries were constructed for sequencing on the Ion S5 platform using a 5′-RACE PCR protocol (*84*). NGS produced ∼1.1 and 1.9 million raw reads for RB35 and RB63, respectively, providing sufficient coverage for both HC and KC repertoires after processing using a rabbit antibodyomics pipeline (**fig. S6B**). B cell repertoire profiles revealed focused HC germline gene usage of IGHV1S40 (>22%), IGHV1S45 (>46%), and IGHV1S47 (>6%) accompanied by a broader and more diverse distribution of KC germline genes, but with some light chains more highly preferred (**fig. S6C**). Notably, RB63, which was immunized with a BG505 gp140.664.R1-FR NP, showed a higher degree of SHM for KCs than RB35, which was immunized with a soluble BG505 trimer (**fig. S6C**). In addition, RB63 appeared to have generated a large percentage (∼50%) of the B cell lineage with a much longer (22-aa) HCDR3 loop (**fig. S6C**). We then investigated the lineage prevalence of three potent rabbit NAbs and four non-NAbs (two per rabbit) within the NGS-derived repertoires (**Fig. 3D** and **fig. S6D**). Using a CDR3 identity cutoff of 95% (90% for RB63-4B5), putative somatic variants were identified for HC and KC of each antibody. All three NAbs exhibited reasonable lineage size, as indicated by the distribution of NGS-derived variants on the 2D plots, whereas non-NAbs showed either no somatic variants or highly expanded population, suggesting that they were either non-specific Env binders or induced by Env vaccination but failed to achieve any neutralizing activity during maturation.

Lastly, we examined whether these potent autologous NAbs target the previously identified glycan holes. We created a set of BG505.T332N Envs bearing mutations Q130N, D230N/K232T, S241N, P291T, and T465N (*55, 58*). Neutralization by three rabbit NAbs was tested in the TZM-bl assay against these BG505.T332N mutants except for D230N/K232T, which was not included due to the low yield of pseudoparticles (**Fig. 3E**). Among the four glycan hole mutations, Q130N did not affect HIV-1 neutralization by any of the three rabbit NAbs. In contrast, S241N and P291T completely abrogated neutralization by RB35-1B11 and RB63-1E7, but not for the more potent RB63-4B5, whereas T465N significantly reduced the potency of RB63-4B5 (IC50 > 5.0 μg/ml), confirming that these three NAbs were targeting the glycan holes at positions 241/289 and 465 that were reported in previous rabbit studies (*55, 58*). We then performed TZM-bl assays to investigate the prevalence of these glycan hole NAbs in total polyclonal NAb response. Indeed, plasma neutralization against the four BG505.T332N mutants demonstrated that filling glycan holes partially depleted the neutralizing activity (**Fig. 3F** and **fig. S6E**), consistent with recent findings that autologous rabbit NAbs could recognize a variety of epitopes other than glycan holes (*57, 59*). Notably, the trimer group showed a more visible reduction when glycan holes at 241/289 were filled, suggesting a preference of trimer-induced autologous NAb response for these two specific sites. Ferritin display was able to diversify but not broaden the NAb response, which nevertheless remained autologous.

By combining single-cell antibody isolation, functional elevation, and repertoire NGS, we demonstrated that a gp41-stabilized BG505 trimer and its FR NP can elicit potent autologous tier 2 NAbs targeting previously identified glycan holes. It remains unclear whether E2p and I3-01 60-mers (*37, 41*) can redirect or broaden the NAb response more effectively than FR 24-mer. EM-based epitope mapping may be used in future studies to reveal other vulnerable sites recognized by trimer and NP-induced polyclonal antibody responses in rabbit immunization (*61*).

### Structural, functional, and in vivo characterization of a tier 2 clade C T/F Env

BG505 trimers, regardless of the design and display platforms, mostly induced glycan hole NAbs in rabbits (*55*). It is therefore imperative to identify HIV-1 Envs capable of eliciting a broader NAb response during immunization. Clade C viruses are important as they are responsible for about half of the global infections (*85*). NFL trimers have been designed for a tier 2 clade C T/F strain, 16055, to facilitate structural analysis of clade C Envs (*31*). This NFL trimer induced a bNAb response in rabbits when displayed on liposome NPs and immunized using a heterologous regimen (*30*). In our previous studies, we demonstrated various HR1 redesigns, as well as UFO and UFO-BG trimer designs, for Env of a tier 2 clade C T/F strain, Du172.17 (*19, 37*). Here, we assessed the potential of this clade C Env as a template for HIV-1 vaccine design.

We first determined the crystal structure for a Du172.17 Env trimer, which is cleaved and contains a strain-specific HR1 redesign [(HR1-#4 (*19*), or simply R4]. This construct, termed Du172.17 gp140.664.R4, was expressed in HEK239S cells and purified on a 2G12 affinity column (*14*) before adding Fabs PGT124 and 35O22 to aid in crystallization. The Fab-bound Du172.17 trimer complex crystallized at 20°C, and its structure was determined at 3.40 Å resolution in an hexagonal (P63) crystal lattice (**Fig. 4A**). The HR1 bend in this construct was designed specifically to stabilize the prefusion Du172.17 Env (*19*) and, therefore, is different from the HR1 bend designed for BG505 Env (**Fig. 4B**). Little difference was observed in the overall Env structure (Cα RMSD = 0.6 Å) between BG505 gp140.664.R1 and Du172.17 gp140.664.R4 except in the N terminus of HR1, termed HR1N (**Fig. 4C**). To further evaluate the difference in overall Env conformation and the redesigned HR1N, we superimposed the Du172.172 protomer onto crystal structures previously determined for clade A, B, and C Envs (**Fig. 4C**). As expected, Du172.17 gp140.664.R4 adopts a protomer structure similar to BG505 SOSIP.664 (*78*), B41 SOSIP.664 (*86*), and 16055 NFL.664 (*31*) with a Cα RMSD of 0.6 Å. Nevertheless, a large conformational change in HR1N was observed between Du172.17 and BG505; comparison to the other Envs with their native-like full-length HR1 was not possible due to disorder in the HR1N helical region in their crystal structures. In addition, we superimposed the Du172.17 gp140.664.R4 protomer onto crystal structures and cryo-EM models of several clade C Envs [PDB ID: 5UM8 (*31*), PDB ID: 6P65 (*30*), PDB ID: 6MYY (*87*), PDB ID: 6UM6 (*88*)]. Although the sequence identity among these clade C Envs is 76-78%, they share a high structural similarity with Cα RMSDs of 0.7-1.3 Å (**fig. S7**). Taken together, the low Cα RMSD values observed for SOSIP, NFL, and HR1-redesigned trimers suggest that the HR1N modification has no adverse impact on the overall Env architecture and compactness.

**Fig. 4.**
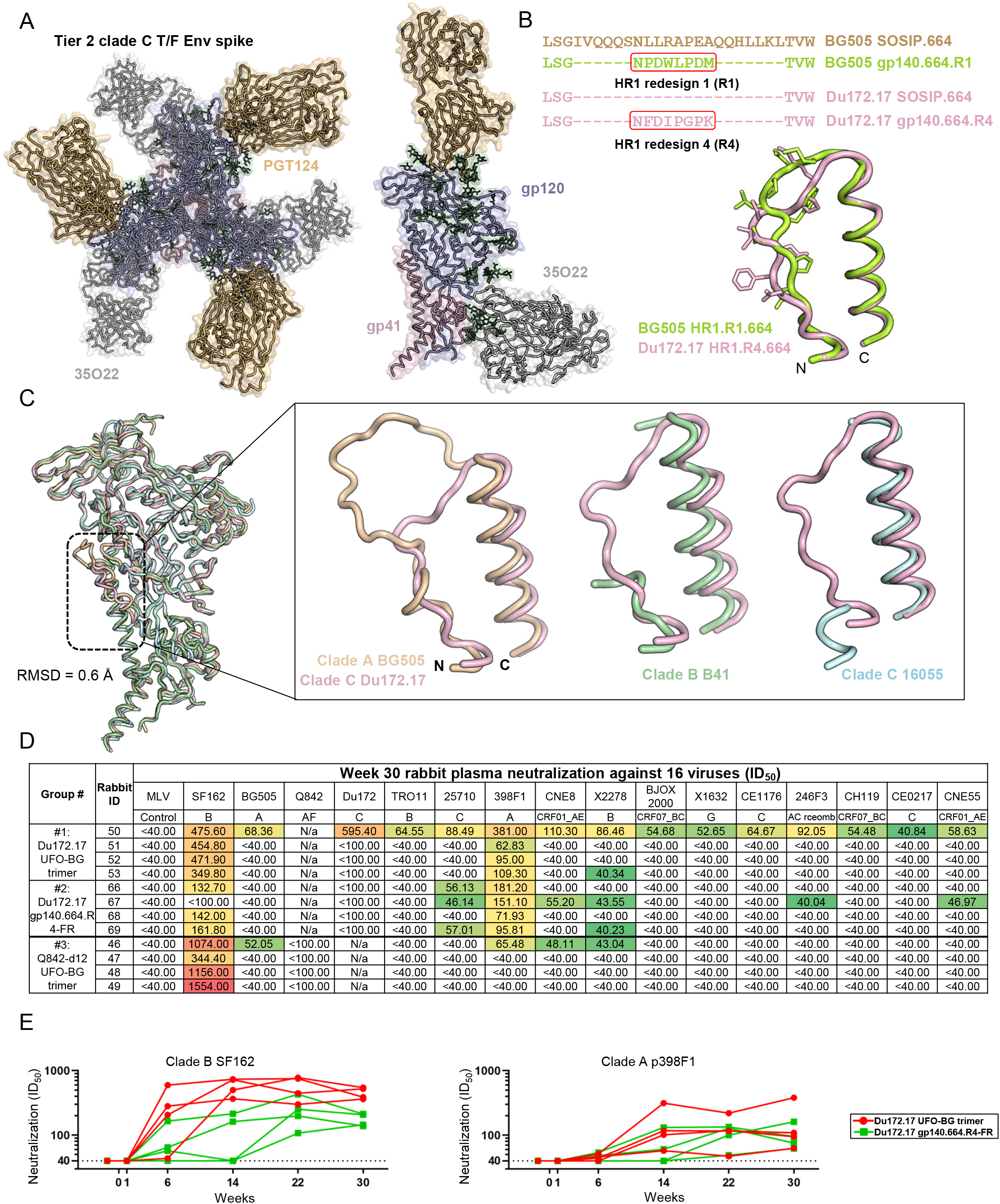
Structural and in vivo characterization of tier-2 clade-C Du172.17 T/F Env. **(A)** Crystal structure of closed prefusion structure of Du172.17 gp140.664.R4 Env trimer, which is uncleaved and contains a computationally redesigned HR1 (HR1-#4, see ref. 19). Top view of the Du172.17 Env-Fab complex along the trimer axis with gp120 in blue and gp41 in pink. Side view of the Du172.17 Env protomer bound to the PGT124 (orange) and 35O22 (dark gray) Fabs from the 3.4 Å resolution crystal structure. The cartoon representation is overlaid with the transparent molecular surface. **(B)** Sequence and structural alignment of the N-terminus of HR1 region (HR1N) in two trimer designs, BG505 gp140.664.R1 and Du172.17 gp140.664.R4. The redesigned 8-residue HR1 region is highlighted to facilitate comparison. **(C)** Superimposition of cleaved Du172.17 gp140.664.R4 (pink), cleaved BG505 SOSIP.664 (orange; PDB ID: 5CEZ), cleaved B41 SOSIP.664 (green; PDB ID: 6MDT) and uncleaved 16055 NFL.664 (cyan; PDB ID: 6P65) protomers. The inset on the right shows a close-up view of Du172.17 HR1N superimposed onto HR1N from each of the three Envs. **(D)** Neutralization (measured by ID50 values) of week-30 plasma from three rabbit groups against two respective autologous viruses (Du172.17 and Q842-d12), tier 2 clade A BG505.T332N, tier 1 clade B SF162, and a 12-virus global panel with MLV included as a negative control. Color coding indicates neutralization potency (red: potent; green neutralizing but not potent; no color: non-neutralizing). **(E)** Longitudinal analysis of plasma from two rabbit groups immunized with Du172.17 trimer and FR nanoparticle at six time points against tier 1 clade B SF162 (left) and tier 2 clade A p398F1 (right). Rabbits from the trimer and FR groups are shown as red and green lines, respectively.

Previously, we compared glycosylation profiles of BG505 SOSIP.664 and HR1-redesigned trimers produced in ExpiCHO cells (*37*). Here, we compared glycosylation patterns of HEK293F-expressed SOSIP, HR1-redesigned, and UFO trimers to determine whether trimer design affects the glycans in the major epitopes on Du172.17 Env. Secreted proteins were harvested from media and purified on a PGT145 affinity column (*15*) followed by size exclusion chromatography (SEC) on a Superdex 200 column. Liquid-chromatography mass spectrometry (LC-MS) was employed to determine site-specific glycosylation (**fig. S8**), which was enabled by digesting the Env proteins into peptides and glycopeptides using three separate proteases: trypsin, chymotrypsin, and elastase. The relative proportions of different glycans were determined and grouped to facilitate comparison between the samples (**fig. S8**). The key features that define native-like glycosylated trimers include the high mannose content and the occupancy at each site. The majority of *N*-linked glycosylation sites on all three trimers contain high amounts of oligomannose-type glycans. Conserved glycan sites across HIV-1 strains that presented oligomannose-type glycans include the N332 supersite and the apex glycan N160. All three design formats have ∼100% occupancy at these sites with oligomannose-type glycans, consistent with a well-folded native-like trimer. The complex-type glycans observed across the samples are fucosylated bi- and tri-antennary glycans that are common in HEK293F or CHO cells. The occupancy at every site in all three trimers is greater than 95% except for N611. However, glycan holes may still be present at sites that could not be resolved in this analysis (**fig. S8**). As expected, the only regions with significant deviation in glycosylation were all within the gp41 ectodomain (gp41ECTO). While no oligomannose-type glycans were observed at N611 on SOSIP, high mannose content was observed on HR1 redesign and UFO, 67% and 24%, respectively. Likewise, SOSIP and HR1 redesign contained 39% and 51% oligomannose-type glycans at N625, whereas UFO had no oligomannose-type glycans at this site. Our data suggest that the steric restrictions imposed upon the glycan sites by surrounding protein regions differ slightly and most of epitopes in gp120 and at the gp120-gp41 interface will not be affected by the design platform.

After structural characterization and glycan analysis of three design platforms for this clade C Env, we assessed two Du172.17 Env immunogens along with a UFO-BG trimer designed for a tier 2 clade A T/F strain, Q842.d12, in three groups of rabbits. To maximize the outcome of *in vivo* assessment for Du172.17 Env, we tested a Du172.17 UFO-BG trimer and a FR NP presenting the structurally defined Du172.17 gp140.664.R4 trimer (**Fig. 4A**). Of note, the same regimen was used to facilitate comparison with the previous BG505 immunization (**Fig. 3A**) (*37*). We first assessed rabbit plasma at the last time point (week 30) against an expanded panel of viruses including the respective autologous virus (either Du172.17 or Q842-d12), BG505.T332N, tier 1 SF162, and the 12-virus global panel, with MLV included as a control (**Fig. 4D** and **fig. S9**). Overall, distinct neutralization patterns were observed compared to the previous BG505 immunization (**Fig. 3B**). Using a 100-fold starting dilution, autologous NAb responses were not observed for any group except for the plasma from RB50 in the Du172.17 trimer group, which also neutralized all tested HIV-1 isolates and exhibited a detectable non-specific response to MLV (**fig. S9A**). The lack of autologous neutralization here therefore differed significantly from our previous study (*37*), where a robust autologous NAb response was observed for three of four rabbits in the BG505 gp140.664.R1-FR group (**Fig. 3B**). Furthermore, the clade C Du172.17 trimer and FR NP elicited consistently, albeit slightly, stronger NAb responses to clade A p398F1 than did their clade A BG505 counterparts, using a 40-fold starting dilution. Weak but consistent plasma neutralization was also developed against other isolates such as clade C p25710 and clade B pX2278 in rabbits immunized with the clade C Du172.17 gp140.664.R4-FR NP, but less so in the other two trimer groups irrespective of the Env origin. In terms of tier 1 NAb response, the clade A Q842-d12 trimer elicited the most potent response with ID50 values of 344 to 1554, whereas the clade C Du172.17 gp140.664.R4-FR NP yielded ID50 values of 162 or lower. As multiple studies have shown that BG505 Env mainly induced autologous NAb responses to glycan holes, our current study indicated that other Envs may elicit a broader response with the potential to neutralize more HIV-1 isolates. Pre-immunization (-d10) samples exhibited negligible reactivity on the 12-virus panel, indicating a clean background (**fig. S9**). Lastly, we characterized the longitudinal NAb development for the clade C Du172.17 trimer and FR groups (**Fig. 4E** and **fig. S10**). The soluble trimer elicited a more rapid tier 1 NAb response to clade B SF162 than the gp140-FR NP and, for most time points, this response also appeared to be more potent than the NP-induced tier 1 NAb response (**Fig. 4E**, left). However, the tier 2 NAb response to clade A p398F1 exhibited a distinct pattern compared to the tier 1 NAb response, with plasma neutralization (measured by ID50) only observed for week 14 and onwards (**Fig. 4E**, right). The difference in such a tier 2 response was not significant between trimer and FR groups, likely due to the small group size and the outlier (RB50) in the trimer group.

The crystal structure of a novel tier 2 clade C T/F Env confirmed the effectiveness of HR1 redesign in trimer stabilization. The rabbit study on three Env immunogens derived from two T/F Envs of different subtypes suggested that the BG505-elicited glycan hole NAb responses may be specific to BG505 Env and that other HIV-1 Envs may achieve a broader NAb response. However, elicitation of a potent bNAb response remains a challenge for HIV-1 vaccine design.

## DISCUSSION

HIV-1 vaccine development has entered a new era since the demonstration of tier 2 NAb responses in rabbits and NHPs elicited by vaccination of prototypic native-like SOSIP.664 trimers (*51, 89*). The success of BG505 SOSIP.664 trimer as a structural template to study bNAb-Env interactions and as an Env backbone to experiment with various rational designs has led to a plethora of studies that position native-like Env trimers at the center of HIV-1 vaccine research (*12, 20*). However, some fundamental questions related to the inherent features of HIV-1 Env need to be addressed to guide future vaccine development (*90*). Critical issues related to trimer design, particulate display platforms, and animal models for vaccine evaluation, will remain open questions while multiple vaccine strategies continue to be explored in parallel.

In this follow-up study, we examined some of these issues by utilizing animal samples generated in our previous study (*37*) and by conducting new immunization experiments. First, we provide evidence at the monoclonal level that potent tier 2 NAbs can be elicited in the wildtype mouse model using a multivalent Env immunogen. The nsEM model and crystal structure of NAb M4H2K1 in complex with BG505 Env identified a target for potent mouse NAbs – an epitope that is recognized by the autologous NAb response in early human infection (*5*). Together with another cross-clade NAb, M1H3K3, our study indicates that wildtype mice provide a useful small animal model for testing HIV-1 vaccines. However, the non-specific antiviral component in mouse serum poses a challenge for pseudovirus neutralization assays and has been the source of inconsistencies in recent studies (*37, 44, 45, 91*). Thus, purified IgG must be used to unambiguously demonstrate the elicitation of tier 2 NAbs in mouse immunization (*37*). Since the tier 2 NAb response can be readily observed in rabbits and such NAbs often target glycan holes, wildtype mice may offer a more advantageous animal model for evaluating HIV-1 vaccine designs and understanding their immunologic mechanisms, as shown in recent studies of multivalently displayed Envs in mice (*91–93*). Second, we provide evidence at the monoclonal level that BG505 Env bearing a redesigned HR1N segment (the core of the UFO trimer design (*19*)), both as a soluble trimer and on a 24-meric FR NP, can induce potent tier 2 NAbs, which, however, are mostly directed to the known glycan holes (*55, 59*). Display of BG505 trimers on this small protein NP did not broaden the autologous NAb response in rabbits (*37*) (**Fig. 3B**). These results, together with the recent finding from a rabbit study of mixed BG505 and B41 SOSIP trimers, highlight a limitation of the rabbit model for evaluation of HIV-1 Env vaccines. Nonetheless, the multivalent display on E2p and I3-01 60-mers may still warrant investigation, as glycan-modified NFL trimers on liposome NPs elicited an impressive cross-clade NAb response in rabbits (*30*). Third, we demonstrated that a strain-specific HR1 redesign could render a stable, native-like trimer for a tier 2 clade C T/F Env, which appeared to modestly broaden the NAb response in rabbits. In contrast, a tier 2 clade A Env-derived UFO-BG trimer exhibited a narrow NAb response. Nonetheless, the crystal structure of Du172.17 Env provides a valuable template for clade C-specific HIV-1 vaccine development. Our new study also indicates that the breadth of vaccine-elicited NAb response may be related to particular features in Env backbone, highlighting the necessity for screening diverse Envs in HIV-1 vaccine design.

Based on our previous studies of Env design and NP display (*19, 37, 41*), several directions may be explored in our future HIV-1 vaccine effort. First, more advanced NP platforms may be employed to display the stabilized Env trimers. Recently, we reengineered E2p and I3-01 60-mers to develop single-component, multilayered, self-assembling protein NPs as vaccine carriers, which were successfully used to present stabilized Ebola virus (EBOV) and SARS-CoV-2 glycoproteins (*94, 95*). These newly engineered NPs offer potential advantages in stability, immunogenicity, and manufacturability in comparison with the two-component NP platforms (*39, 40*). Second, an in-depth immunological understanding will be crucial for the future development of UFO trimers and UFO-NP vaccines, as demonstrated for other HIV-1 vaccine candidates (*91–93, 96*). In our proof-of-concept study of hepatitis C virus (HCV) vaccines, NPs presenting optimized E2 cores elicited NAb responses more effectively than E2 core alone, with quantitative B cell patterns revealed by NGS (*97*). Analysis of NP retention, trafficking and germinal center activation (*98*) may provide the much-needed insight into the mode of action of NP vaccines. Critical questions related to HIV-1 Env (*90*) can therefore be pursued in the context of UFO trimers and UFO-NPs.

## MATERIALS and METHODS

### Expression and purification of HIV-1 Env probes, trimers, and gp140 nanoparticles

The Avi-tagged BG505 gp140.664.R1 trimer probe was transiently expressed in HEK293F cells (Thermo Fisher) (*67*). Env protein was purified from the supernatant by a *Galanthus nivalis* lectin (GNL) column (Vector Labs) and eluted with PBS containing 500 mM NaCl and 1 M methyl-α-D-mannopyranoside. Biotinylation was performed using the BirA biotin-protein ligase standard reaction kit (BirA-500) as per the manufacturer’s instructions (Avidity). This BG505 trimer probe was further purified by SEC on a HiLoad 16/600 Superdex 200 PG column (GE Healthcare). A ferritin (FR) NP presenting BG505 V1V2 and a trimeric scaffold (1TD0) presenting ZM109 V1V2 were transiently expressed in *N*-acetylglucosaminyltransferase I-negative (GnTI^-/-^) HEK293S cells (Thermo Fisher) (*41*). Both FR and I3-01 NPs presenting an N332-scaffold, 1GUT_A_ES, were transiently expressed in HEK293F cells treated with Kifunensine (TOCRIS Bioscience) (*69*). Both V1V2 and N332 epitope probes were extracted from the supernatant using a GNL column. Fusion peptide (FP) probes were created by fusing the FP motif, AVGIGAVFL, to a FR or 1TD0 subunit with a 5GS (G4S) linker. The FP-5GS-FR and BG505 gp120-FR probes were transiently expressed in ExpiCHO cells (Thermo Fisher) using a similar protocol to BG505 gp140 NPs (*37*). The trimeric FP-5GS-1TD0 probe was also transiently expressed in ExpiCHO cells. Immunoaffinity columns based on bNAbs VRC34 (*99*) and PGT145 (*100*) were used to extract two FP probes (FP-5GS-FR NP and FP-5GS-1TD0 trimer) and the BG505 gp120-FR probe from the supernatant, respectively. After purification using a GNL or antibody column, the NP and 1TD0-derived epitope probes were further purified by SEC on a Superose 6 10/300 GL column and a Superdex 75 10/300 GL column (GE Healthcare), respectively. For rabbit immunization, the Du172.17 UFO-BG trimer, the FR NP presenting a structurally defined Du172.17 gp140.664.R4 trimer, which is cleaved and contains a redesigned HR1 (HR1-#4 (*19*)), and the Q842-d12 UFO-BG trimer were transiently expressed in ExpiCHO cells (*37*). For the two UFO-BG trimers, Env protein was extracted from the supernatant by a GNL column and trimer was purified by SEC on a HiLoad 16/600 Superdex 200 PG column. The Du172.17 gp140.664.R4-FR NP was purified using a 2G12 affinity column (*14*) followed by SEC on a Superose 6 10/300 GL column. Protein concentrations were determined using ultraviolet absorbance at 280 nm (UV280) with theoretical extinction coefficients.

### Env-specific sorting of mouse and rabbit B cells

Mouse spleen cells harvested 15 days after the last injection were prepared for sorting. Cells were first stained for the exclusion of dead cells with Fixable Aqua Dead Cell Stain (Thermo Fisher). Receptors FcγIII (CD16) and FcγII (CD32) were blocked by 20 μl of 2.4G2 mAb (BD Pharmigen). Cells were then incubated with 10 μg of biotinylated Avi-tagged BG505 gp140.664.R1 trimer probe for 5 min at 4 °C, followed by the addition of 2.5 μl of anti-mouse IgG fluorescently labeled with FITC (Jackson ImmunoResearch) and incubated for 15 min at 4 °C. Finally, 5 μl of premium-grade allophycocyanin (APC)-labeled streptavidin (Thermo Fisher) was added to the cells and incubated for 15 min at 4 °C. In each step, cells were washed with 500 μl FACS buffer (DPBS with 2% FBS). FITC^+^APC^+^ Env-specific B cells were sorted using MoFloAstrios EQ (Beckman Coulter). Rabbit PBMCs obtained 30 days after the last injection were prepared for sorting. After staining for exclusion of dead cells with Fixable Aqua Dead Cell Stain (Thermo Fisher), cells were incubated with 10 μg of biotinylated Avi-tagged BG505 gp140.664.R1 trimer probe for 5 min at 4 °C, followed by addition of 2 μl of anti-rabbit IgG conjugated with Dylight 405 (Jackson ImmunoResearch), 2 μl of anti-rabbit T lymphocytes fluorescently labeled with FITC (BioRad), and 2 μl of anti-rabbit IgM fluorescently labeled with FITC (BioRad), and then incubated for 15 min at 4 °C. Finally, 5 μl of APC-labeled streptavidin (Thermo Fisher) was added to the cells and incubated for 15 min at 4 °C. In each step, cells were washed with 500 μl FACS buffer (DPBS with 2% FBS). FITC^−^Dylight 405^+^APC^+^ Env-specific B cells were sorted using MoFloAstrios EQ (Beckman Coulter). For bulk sorting, positive cells were sorted into an Eppendorf microtube with 20 μl of lysis buffer. For single B-cell sorting, individual positive cells were sorted into the inner wells of a 96-well plate with 20 μl of a pre-reverse transcription (RT) lysis mix containing 0.1 μl NP40 (Sigma-Aldrich), 0.5 μl RNAse Inhibitor (Thermo Fisher), 5 μl 5× First Strand Buffer and 1.25 μl DTT from SuperScript IV kit (Invitrogen), and 13.15 μl H2O per well.

### Antibody cloning from Env-specific single B cells and antibody production

Antibody cloning of Env-sorted single B cells was conducted as follows. A mix containing 3 μl Random Hexamers (GeneLink), 2 μl dNTPs, and 1 μl of the SuperScript IV enzyme (Thermo Fisher) was added to each well of a single-cell sorted 96-well plate that underwent thermocycling according to the program outlined in the SuperScript IV protocol resulting in 25 μl of cDNA for each single cell. 5 μl of cDNA was then added to a PCR mix containing 12.5 μl 2× Multiplex PCR mix (Qiagen), 9 μl H2O, 0.5 μl of forward primer mix, and 0.5 μl of reverse primer mix (mouse (*101*) and rabbit (*55*)) for heavy and κ-light chains within each well. A second PCR reaction was then performed using 5 μl of the first PCR as template and respective primers (mouse (*101*) and rabbit (*55*)) utilizing the same recipe as the first PCR. The PCR products were run on 1% Agarose gel and those with correct heavy and light chain bands were then used for Gibson ligation (New England Biolabs), cloning into IgG expression vectors, and transformation into competent cells. Mouse and rabbit mAbs were expressed by transient transfection of ExpiCHO cells (Thermo Fisher) with equal amount of paired heavy and κ-light chain plasmids and purified from the culture supernatant after 12-14 days using Protein A beads columns (Thermo Fisher).

### NGS and bioinformatics analysis of mouse and rabbit B cells

We combined the 5′-rapid amplification of cDNA ends (RACE) protocol with previously reported heavy and κ-light chain primers for mouse (*101*) and rabbit (*84*) to facilitate NGS analysis of Env-specific mouse splenic B cells and rabbit B cells, respectively. Briefly, 5′-RACE cDNA was obtained from bulk-sorted B cells of each animal with SMART-Seq v4 Ultra Low Input RNA Kit for Sequencing (TaKaRa). The Ig PCRs were set up with Platinum *Taq* High-Fidelity DNA Polymerase (Thermo Fisher) in a total volume of 50 µl, with 5 μl of cDNA as template, 1 μl of 5′-RACE primer, and 1 μl of 10 µM reverse primer. The 5′-RACE primer contained a PGM/S5 P1 adaptor, while the reverse primer contained a PGM/S5 A adaptor. For mouse samples, we adapted the mouse 3′-Cγ1-3/3′-Cμ inner primers and 3′-mCκ outer primer (*101*) as reverse primers for 5′-RACE PCR processing of heavy and κ-light chains, respectively. For rabbit samples, we adapted rabbit RIGHC1/RIGHC2 primers and RIGkC primers (*84*) as reverse primers for 5′-RACE PCR processing of heavy and κ-light chains, respectively. A total of 25 cycles of PCR was performed and the expected PCR products (500-600 bp) were gel purified (Qiagen). NGS was performed on the Ion S5 GeneStudio platform. Briefly, heavy and κ-light chain libraries from the same animal were quantitated using Qubit® 2.0 Fluorometer with Qubit® dsDNA HS Assay Kit, and then mixed at a ratio of 2:1 or 3:1 before being pooled with antibody libraries from the other animals at an equal ratio. Template preparation and Ion 530 chip loading were performed on Ion Chef using the Ion 520/530 Ext Kit, followed by sequencing on the Ion S5 system with default settings. The mouse antibodyomics pipeline (*69*) was used to process the mouse NGS data. The rabbit antibodyomics pipeline was created by incorporating rabbit germline genes from IMGT (http://www.imgt.org/) into the reference libraries. Quantitative repertoire profiles were generated for germline gene usage, degree of SHM, and H/KCDR3 loop length. The two-dimensional (2D) divergence/identity plots were generated to visualize selected mouse and rabbit NAb/mAb chains in the context of Env-specific B cell repertoires. A previously described sequence clustering algorithm (*67*) was used to derive consensus heavy and κ-light chains for prevalent antibody lineages from NGS data of bulk-sorted mouse splenic B cells. NGS-derived mAbs were transiently expressed in ExpiCHO cells (Thermo Fisher) with equal amount of heavy and κ-light chain plasmids and purified from culture supernatants after 12-14 days using Protein A beads columns (Thermo Fisher).

### Enzyme-linked immunosorbent assay

Each well of a Costar^TM^ 96-well assay plate (Corning) was first coated with 50 µl PBS containing 0.2 μg of the appropriate antigens. The plates were incubated overnight at 4 °C, and then washed five times with wash buffer containing PBS and 0.05% (v/v) Tween 20. Each well was then coated with 150 µl of a blocking buffer consisting of PBS, 40 mg ml^-1^ blotting-grade blocker (Bio-Rad), and 5% (v/v) FBS. The plates were incubated with the blocking buffer for 1 hour at room temperature, and then washed five times with wash buffer. For antigen binding, antibodies were diluted in the blocking buffer to a maximum concentration of 10 μg ml^-1^ followed by a 10-fold dilution series. For each antibody dilution, 50 μl was added to the appropriate wells. Next, a 1:5000 dilution of goat anti-human IgG antibody (Jackson ImmunoResearch Laboratories, Inc) was made in the wash buffer (PBS containing 0.05% Tween 20), with 50 μl of the diluted secondary antibody added to each well. The plates were incubated with the secondary antibody for 1 hour at room temperature, and then washed five times with PBS containing 0.05% Tween 20. Finally, the wells were developed with 50 μl of TMB (Life Sciences) for 3-5 min before stopping the reaction with 50 μl of 2 N sulfuric acid. The resulting plate readouts were measured at a wavelength of 450 nm. The EC50 values were calculated in GraphPad Prism 8.4.3.

### Pseudovirus Production and Neutralization Assays

Pseudoviruses were generated by transfection of HEK293T cells with an HIV-1 Env expressing plasmid and an Env-deficient genomic backbone plasmid (pSG3ΔEnv), as previously described (*102*). HIV-1 Env expressing vectors for BG505 (Cat# 11518), SF162 (Cat# 10463), and the global panel (*71*) (Cat# 12670) were obtained through the NIH AIDS Reagent Program, Division of AIDS, NIAID, NIH (https://www.aidsreagent.org/). A T332N mutation was introduced into BG505 Env to produce the BG505.T332N clone. Other BG505.T332N mutants were created by introducing mutations as previously described (*55, 59, 81*). Pseudoviruses were harvested 72 hours post-transfection for use in neutralization assays. Neutralizing activity of heat-inactivated rabbit plasma was assessed using a single round of replication pseudovirus assay and TZM-bl target cells, as described previously (*102*). Briefly, pseudovirus was incubated with serial dilutions of antibodies or rabbit plasma in a 96-well flat bottom plate for 1 hour at 37 °C before TZM-bl cells were seeded in the plate. For antibody neutralization, a starting concentration of 5 μg/μl was used and subjected to a 3-fold dilution series in the TZM-bl assays. Rabbit plasma was diluted by 100-fold and 40-fold against autologous and heterologous pseudoviruses, respectively, and then subjected to a 3-fold dilution series in the TZM-bl assays. As a negative control, pseudoparticles displaying the envelope glycoproteins of murine leukemia virus (MLV) were tested in the TZM-bl assays following the same protocol. Luciferase reporter gene expression was quantified 48-72 hours after infection upon lysis and addition of Bright-GloTM Luciferase substrate (Promega). Data were retrieved from a BioTek microplate reader with Gen 5 software, the background luminescence from a series of uninfected wells was subtracted from each experimental well, and neutralization curves were generated using GraphPad Prism 8.4.3, in which values from experimental wells were compared against a well containing virus only. To determine IC50 and ID50 values, dose-response curves were fit by nonlinear regression in GraphPad Prism 8.4.3.

### Expression, purification of BG505 gp120 core, Du172.17 gp140, Fabs and complex formation

The antigen-binding fragments (Fabs) of M4H2K1, 17b, PGT124, and 35O22 were expressed in FreeStyle HEK293F cells (Invitrogen) and purified by CaptureSelect CH1-XL affinity (Thermo Fisher) chromatography followed by SEC on a Superdex 75 16/600 column (GE Healthcare). The BG505 gp120 core protein was transiently expressed in FreeStyle HEK293S cells, extracted from the supernatant using a GNL affinity column, followed by SEC on a Superdex 200 16/600 column (GE Healthcare). A complex was formed by combining gp120:M4H2K1:17b in a 1:2:2 molar ratio, followed by deglycosylation using endoH digestion (New England Biolabs) at 37 °C for 1 hour before SEC purification. The gp120:M4H2K1:17b complex was then analyzed by sodium dodecyl sulphate–polyacrylamide gel electrophoresis (SDS-PAGE). The clade C Du172.17 gp140.664.R4 Env trimer was expressed in FreeStyle HEK293S cells. Env protein was harvested from media and purified with a 2G12 column (*14*) followed by SEC on a Superdex 200 column (GE Healthcare). The Du172.17 trimer complex was formed by mixing PGT124 and 35O22 Fabs in a molar ratio of 1:3.5:3.5 (Du172:PGT124:35O22) at room temperature for 30 min. The trimer complex was then partially deglycosylated using endoH digestion (New England Biolabs) (*23*) at 37 °C for 1 hour and then purified on a Superdex 200 column. The complex was SEC-purified in 50 mM Tris-HCl, 150mM NaCl (pH = 7.4) and concentrated to ∼10 mg/ml prior to crystallization trials.

### Crystallization and data collection

The SEC-purified Fab M4H2K1 and complex were each concentrated to 12 mg/ml before being screened at both 4 °C and 20 °C using our high-throughput CrystalMation^TM^ robotic system (Rigaku) at TSRI (*103*). High-quality crystals of unbound Fab M4H2K1 were grown in 0.1 M CHES (pH = 9.5) and 36% PEG600 at 4 °C, and M4H2K1 bound gp120 core complex in 0.1 M Tris (pH = 7), 1.825 M ammonium sulfate, 0.29 M lithium sulfate, and 15% ethylene glycol at 20°C. Crystals were harvested, and followed by immediate flash cooling in liquid nitrogen. The Du172.17 trimer complex was set up at both 4 °C and 20 °C using our Rigaku CrystalMation^TM^ robotic system. High-quality crystals of Fabs PGT124 and 35O22 bound to the HR1-redesigned Du172.17 trimer were obtained in 0.1 M Tris (pH = 8.4), 25% (v/v) PEG400 at 20 °C. Data were collected at Advanced Photon Source (APS) on beamlines 23-IDD and 23-IDB.

### Structure determination and refinement

The unbound Fab M4H2K1 and the Fab M4H2K1-BG505 gp120 core -Fab 17b crystals diffracted to 1.50 Å and 4.30 Å resolution, respectively. The data were indexed, integrated and scaled using HKL2000 (*104*) in P3121 for unbound M4H2K1-Fab and in P21212 for the complex. The unbound Fab structure was solved by molecular replacement (MR) using Phaser (*105*) with Fab structures [PDB 5GS1 (*106*) for the variable region and PDB 5BZW (*107*) for the constant region] as MR search models. The BG505 gp120 core in complex with Fabs M4H2K1 and 17b was determined by MR using PDB 6ONF (*108*) for the gp120 core, the unbound Fab M4H2K1 structure for the bound Fab M4H2K1, and PDB 1GC1 (*77*) for Fab 17b as the search models. The unbound M4H2K1 Fab crystal structure was refined to *R_cryst_ / R_free_* of 14.5%/18.4% with 99.8% completeness and unit cell parameters *a* = *b* = 68.3Å, *c* = 184.7Å (**Table S1**). The Fab M4H2K1 bound gp120 core complex structure was refined to *R_cryst_ / R_free_* of 29.7%/33.3% with 86.8% completeness and unit cell parameters *a*= 204.0Å, *b*= 60.6Å, *c*= 166.7Å (**Table S1**). The Du172.17 trimer in complex with Fabs PGT124 and 35O22 crystal diffracted to 3.40 Å resolution and the diffraction data were processed (indexed, integrated and scaled) with HKL2000 in P63 space group. The Du172.17 trimer in complex with Fabs PGT124 and 35O22 was determined by using PDB 5CEZ (*78*) for Env Du172.17 gp140, PDB 4TOY (*109*) for 35O22, and PDB 4R26 (*110*) for the Fab PGT124 structure as the MR search models. The crystal structure of the Du172.17 trimer complex was refined to *R_cryst_ / R_free_* of 29.7%/31.8% and overall completeness of 97.6% and unit cell parameters *a*=*b*= 127.0Å, *c*= 316.5Å (**Table S1**). Model building was carried out with Coot and refinement with Phenix (*111–113*). Structure quality was determined by MolProbity (*114*). The Kabat numbering scheme (*115*) was used for Fabs M4H2K1 and 17b. The BG505 gp120 core and Du172.17 trimer were numbered according to the HXB2 system (*116*). Structure validation was performed using the PDB Validation Server (validate.wwpdb.org), PDB-care (*117*) and Privateer (*118*). Data collection and refinement statistics are shown in **Table S1**.

### Negative-stain electron microscopy

Complexes of M4H2K1 Fab and BG505 UFO.664 trimer were purified by SEC to remove unbound Fab and diluted to 0.01 mg/mL in Tris-buffered saline prior to adsorption onto carbon-coated and plasma cleaned copper mesh grids (Cu400, Electron Microscopy Sciences). Grids were stained with 2% (w/v) uranyl formate for about 60 s and imaged on an FEI Tecnai Spirit microscope operating at 120 keV, equipped with a TVIPS TemCam F416 4k × 4k CMOS camera. Automated data collection was performed using Leginon (*119*). Particles were picked using DogPicker in the Appion software suite (*120*), extracted using Relion 3.0 (*121*), and imported into cryoSPARC v2 (*122*). After one round each of 2D and 3D classification, 14,027 particles were included in a 3D refinement with C3 symmetry imposed and a low pass-filtered volume of ligand-free HIV-1 Env used as the initial model. The final resolution for the negative-stain reconstruction is estimated to be ∼25 Å (Fourier shell correlation cutoff of 0.5).

### Glycopeptide analysis by mass spectrometry

Three 50 μg aliquots of each sample were denatured for 1h in 50 mM Tris/HCl, pH 8.0 containing 6 M of urea and 5 mM dithiothreitol (DTT). Next, Env proteins were reduced and alkylated by adding 20 mM iodoacetamide (IAA) and incubated for 1h in the dark, followed by a 1h incubation with 20 mM DTT to eliminate residual IAA. The alkylated Env proteins were buffer-exchanged into 50 mM Tris/HCl, pH 8.0 using Vivaspin columns (3 kDa) and digested separately overnight using trypsin, chymotrypsin or elastase (Mass Spectrometry Grade, Promega) at a ratio of 1:30 (w/w). The next day, the peptides were dried and extracted using C18 Zip-tip (MerckMilipore). The peptides were dried again, re-suspended in 0.1% formic acid and analyzed by nanoLC-ESI MS with an Easy-nLC 1200 (Thermo Fisher Scientific) system coupled to a Fusion mass spectrometer (Thermo Fisher Scientific) using higher energy collision-induced dissociation (HCD) fragmentation. Peptides were separated using an EasySpray PepMap RSLC C18 column (75 µm × 75 cm). A trapping column (PepMap 100 C18 3μM 75μM × 2cm) was used in line with the LC prior to separation with the analytical column. The LC conditions were as follows: 275 min linear gradient consisting of 0-32% acetonitrile in 0.1% formic acid over 240 min followed by 35 min of 80% acetonitrile in 0.1% formic acid. The flow rate was set to 200 nl/min. The spray voltage was set to 2.7 kV and the temperature of the heated capillary was set to 40 °C. The ion transfer tube temperature was set to 275 °C. The scan range was 400-1600 *m/z*. The HCD collision energy was set to 50%, appropriate for fragmentation of glycopeptide ions. Precursor and fragment detection were performed using an Orbitrap at a resolution MS1= 100,000. MS2= 30,000. The AGC target for MS1 = 4e5 and MS2 = 5e4 and injection time: MS1 = 50ms MS2 = 54ms.

Glycopeptide fragmentation data were extracted from the raw file using Byonic^TM^ (Version 3.5) and Byologic^TM^ software (Version 3.5; Protein Metrics Inc.). The glycopeptide fragmentation data were evaluated manually for each glycopeptide; the peptide was scored as true-positive when the correct b and y fragment ions were observed along with oxonium ions corresponding to the glycan identified. The MS data were searched using the Protein Metrics 305 N-glycan library. The relative amounts of each glycan at each site, as well as the unoccupied proportion, were determined by comparing the extracted chromatographic areas for different glycotypes with an identical peptide sequence. All charge states for a single glycopeptide were summed. The precursor mass tolerance was set at 4 part per million (ppm) and 10 ppm for fragments. A 1% false discovery rate (FDR) was applied. Glycans were categorized according to the composition detected. HexNAc(2)Hex(9−5) was classified as M9 to M5, HexNAc(3)Hex(5−6)X as Hybrid with HexNAc(3)Fuc(1)X classified as Fhybrid. Complex-type glycans were classified according to the number of HexNAc residues, which are attributed to number of processed antenna/bisecting GlcNAc (B), and fucosylation (F). For example, HexNAc(3)Hex(3-4)X is assigned to A1, HexNAc(4)X to A2/A1B, HexNAc(5)X to A3/A2B, and HexNAc(6)X to A4/A3B. If all of these compositions had a fucose, they are assigned to the corresponding FA category. Note that this analytical approach does not distinguish between isomers, which could influence the formal assignment of number of antennae in some cases.

### Rabbit immunization and sample collection

The Institutional Animal Care and Use Committee (IACUC) guidelines were followed with animal subjects tested in the immunization study. Rabbit immunization and blood sampling were carried out under a subcontract at Covance (Denver, PA) following a previously described protocol (*37*). Three groups of female New Zealand White rabbits, four rabbits per group, were immunized intramuscularly with 30 μg of trimer or NP formulated in 250 μl of adjuvant AddaVax (InvivoGen) with a total volume of 500 μl, at weeks 0, 4, 12, 20, and 28. Blood samples, 15 ml each time, were collected at day -10, weeks 1, 6, 14, 22, 28, and 30. Plasma was separated from blood and heat inactivated for ELISA binding and TZM-bl neutralization assays.

## SUPPLEMENTARY MATERIALS

Supplementary material for this article is available at XXX.

**fig S1.**
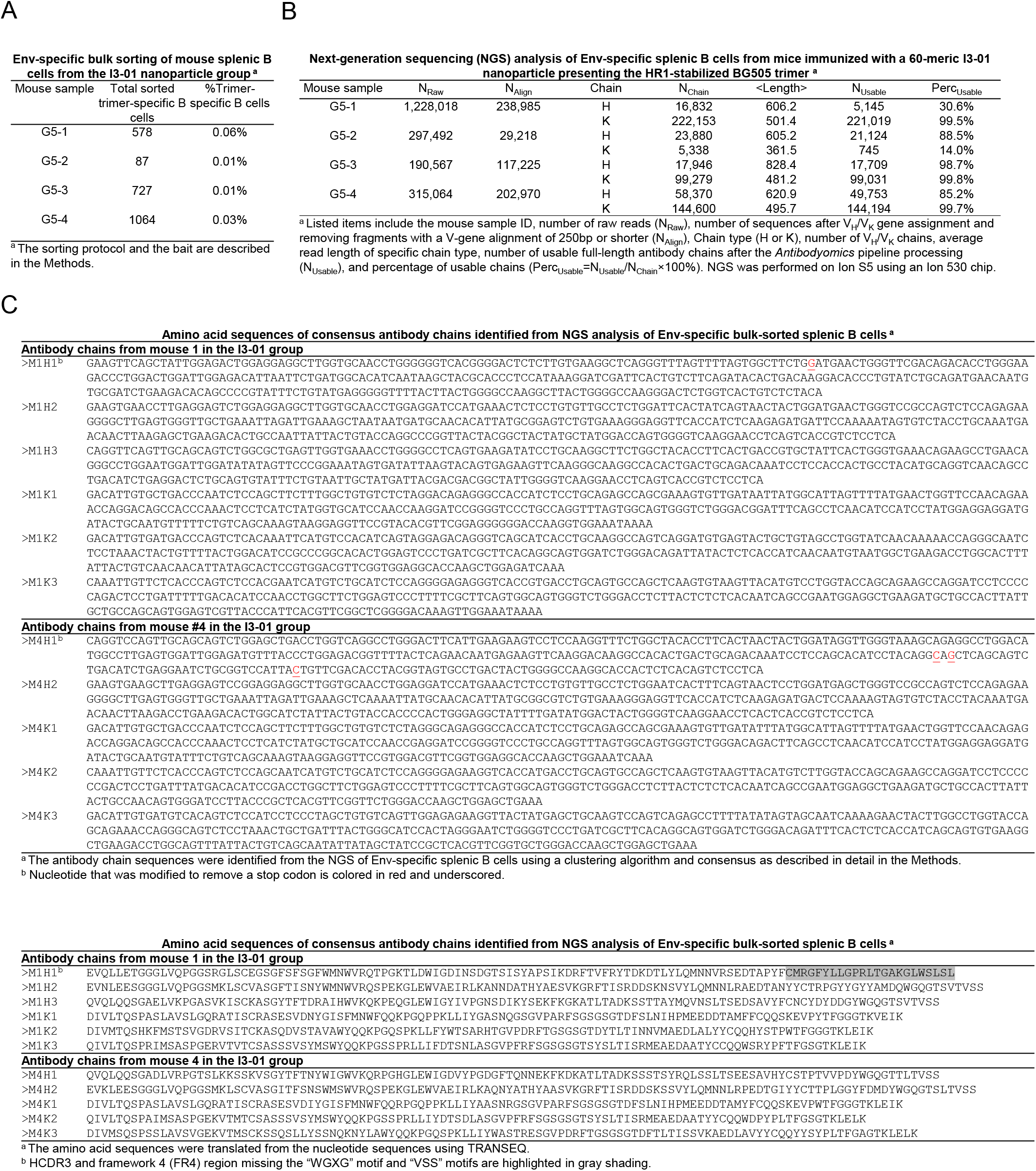
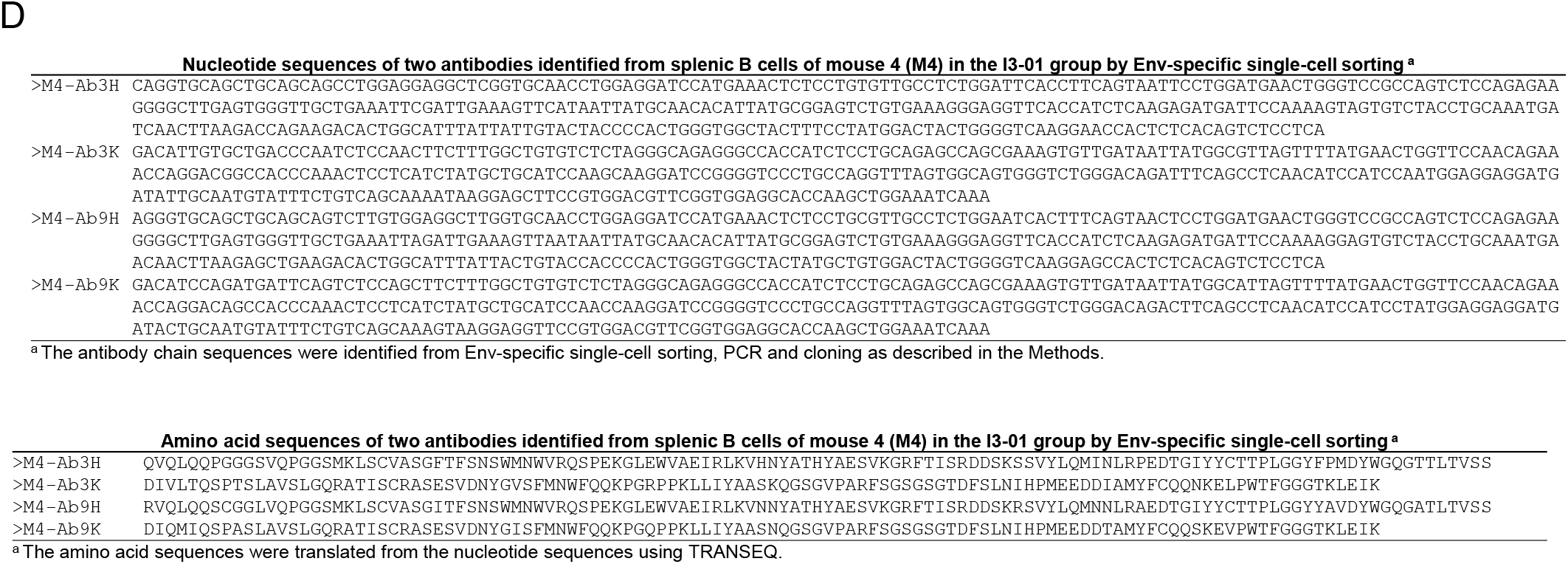
HIV-1 Env-specific sorting and NGS of mouse splenic B cells for antibody isolation. Mice immunized with BG505 gp140.664.R1-PADRE-I3-01 nanoparticle (see ref. 37) were analyzed in this study. (**A**) Env-specific mouse splenic B cells obtained from bulk sorting using a biotinylated Avi-tagged BG505 gp140.664.R1 trimer probe. (**B**) Antibodyomics pipeline processing of NGS data obtained from sequencing of Env-specific mouse splenic B cells on the Ion S5 platform. (**C**) Nucleotide and amino acid sequences of consensus antibody heavy and κ-light chains (HC and KC) identified from NGS analysis of Env-specific splenic B cells from mice 1 and 4 (M1 and M4) in the I3-01 nanoparticle group. (**D**) Nucleotide and amino acid sequences of two antibodies, Ab3 and Ab9, identified by single-cell sorting and antibody cloning from M4 splenic B cells. Ab3 and Ab9 use the same germline genes as the NGS-derived NAb, M4H2K1.

**fig S2.**
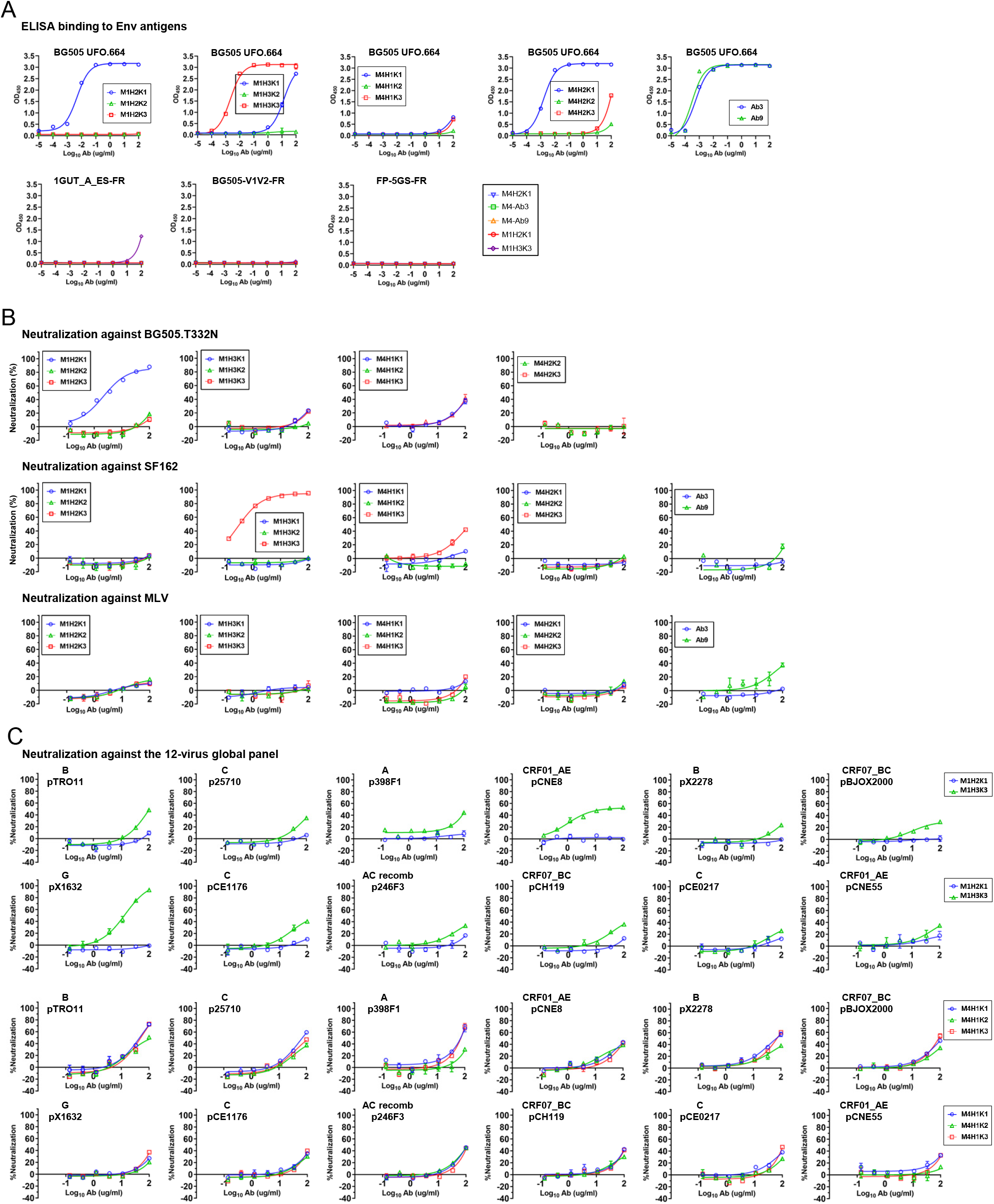
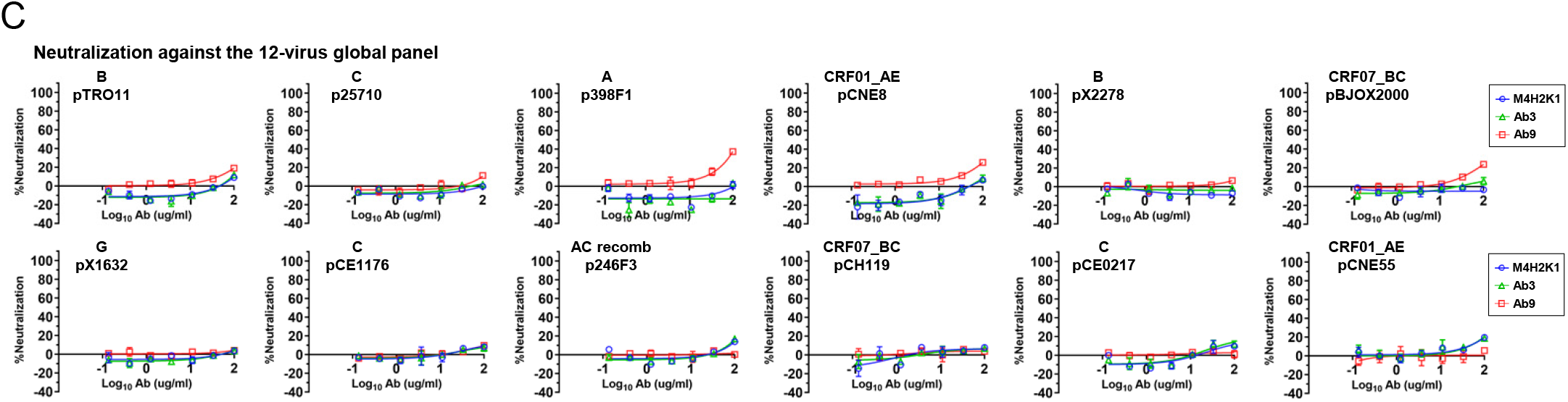
Functional evaluation of NGS and single-cell-derived mouse mAbs. (**A**) ELISA binding of mouse mAbs to Env antigens including BG505 UFO.664 trimer (top panel) and three individual epitope probes including 1GUT_A_ES-FR (N332 supersite), BG505 V1V2-FR (V1V2 apex), and FP-5GS-FR (fusion peptide), which are all ferritin nanoparticles. **(B)** Neutralization of autologous tier 2 clade A BG505.T332N by mouse mAbs. **(C)** Neutralization of all 12 isolates from a global panel by mouse mAbs.

**fig. S3.**
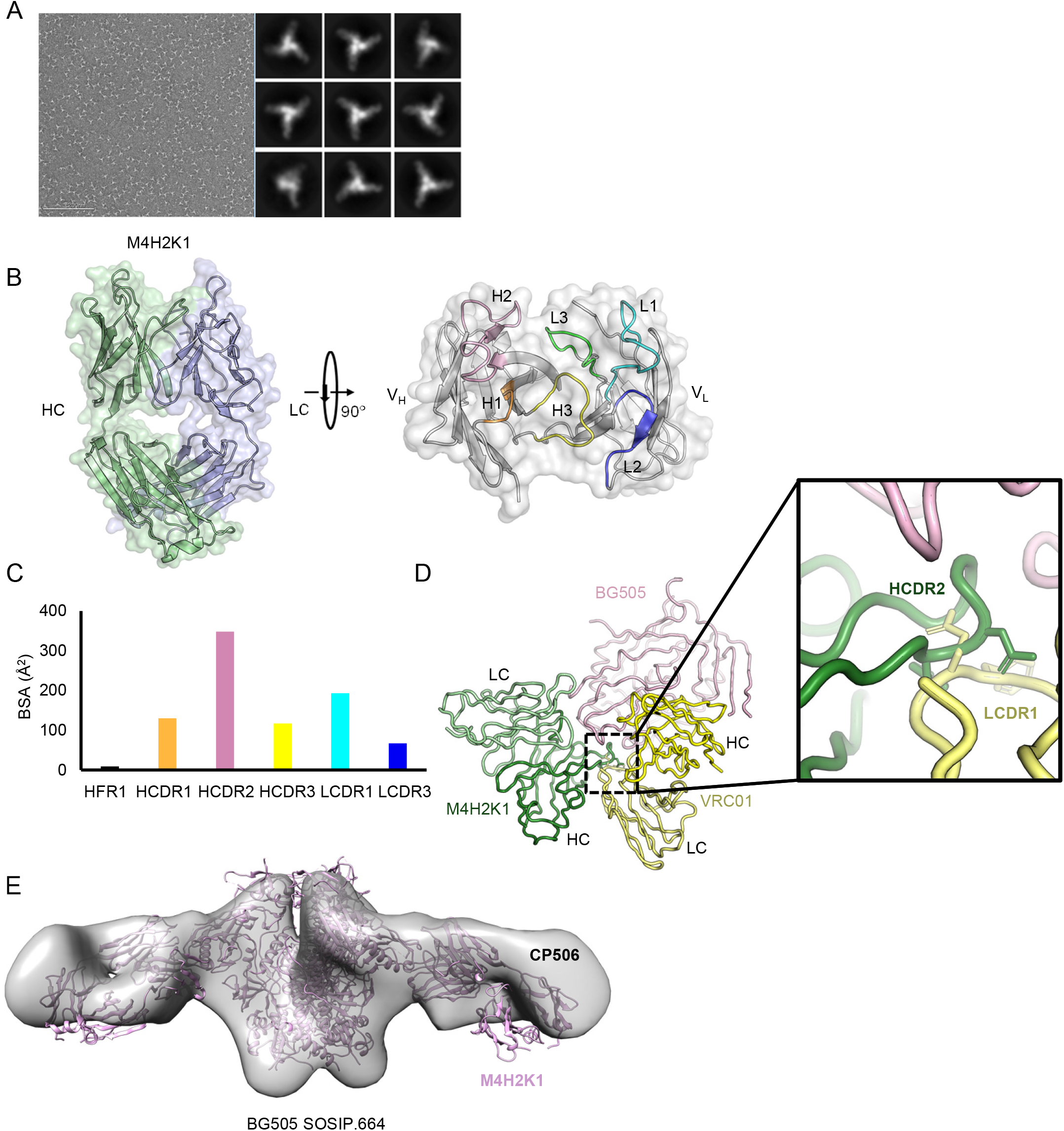
Structural characterization of the NGS-derived mouse NAb, M4H2K1. (**A**) Negative-stain EM (nsEM) analysis of mouse NAb M4H2K1 in complex with BG505 UFO.664 Env trimer. Left: EM micrograph; Right: 2D class averages. (**B**) The unbound structure of M4H2K1 in a ribbons model within the molecular surface. Left: side view; Right: top view. The H/LCDR loops are labeled on the structure. (**C**) Buried surface area (Å^2^) of the CDR loops and FRs of M4H2K1 Fab when bound to BG505 gp120 core. (**D**). Superimposition of VRC01 (yellow) Fab-bound BG505 SOSIP with M4H2K1 (green) Fab-bound BG505 core (pink). The right inset shows a clash of M4H2K1 HCDR2 with VRC01 LCDR1. (**E**) Comparison of the mode of recognition for M4H2K1 and CP506 when bound to the BG505 SOSIP.664 Env trimer.

**fig. S4.**
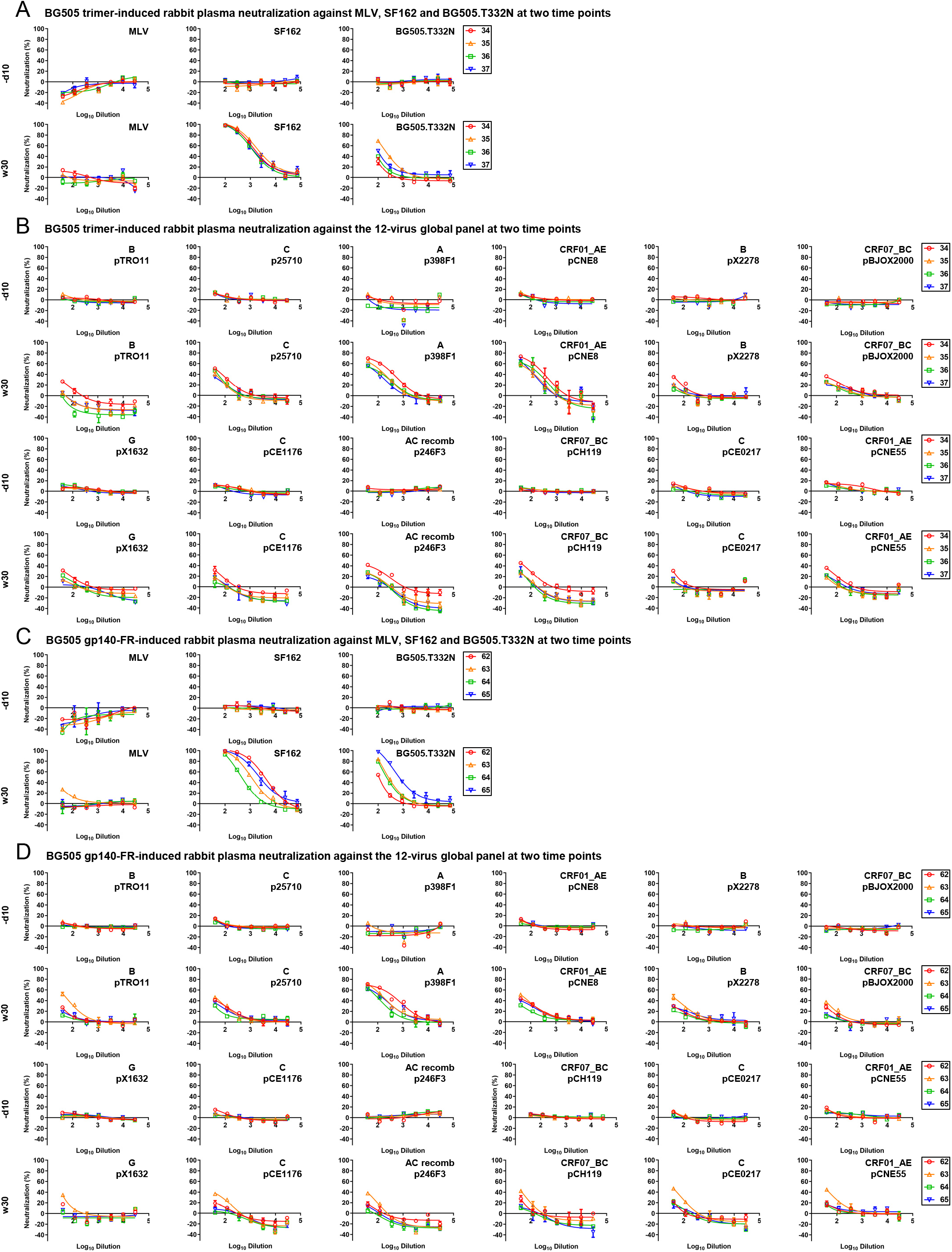
Rabbit plasma neutralization from two BG505 Env-immunized rabbit groups. In the previous study (see ref. 37), two groups of rabbits were immunized with BG505 gp140.664.R1 trimer and its ferritin nanoparticle. (**A**) Neutralization of MLV, tier 1 clade B SF162, and tier 2 clade A BG505.T332N by day -10 (-d10) and week 30 (w30) rabbit plasma from the BG505 trimer group. (**B**) Neutralization of 12 isolates in the global panel by day -10 (-d10) and week 30 (w30) rabbit plasma from the BG505 trimer group. (**C**) Neutralization of MLV, SF162, and BG505.T332N by day -10 (-d10) and week 30 (w30) rabbit plasma from the BG505 ferritin nanoparticle group. (**D**) Neutralization of 12 isolates in the global panel by day -10 (-d10) and week 30 (w30) rabbit plasma from the BG505 ferritin nanoparticle group. The heat-inactivated rabbit plasma was diluted 100-fold for autologous tier 2 BG505.T332N and tier 1 SF162 and subjected to a 3-fold dilution series in the TZM-bl assay. To increase the sensitivity of detection, heat-inactivated plasma was diluted 40-fold for MLV and all 12 isolates from a global panel and followed by a 3-fold dilution series in the TZM-bl assays. ID_50_ titers for plots (A) – (D) are shown in Fig. 3B.

**fig. S5.**
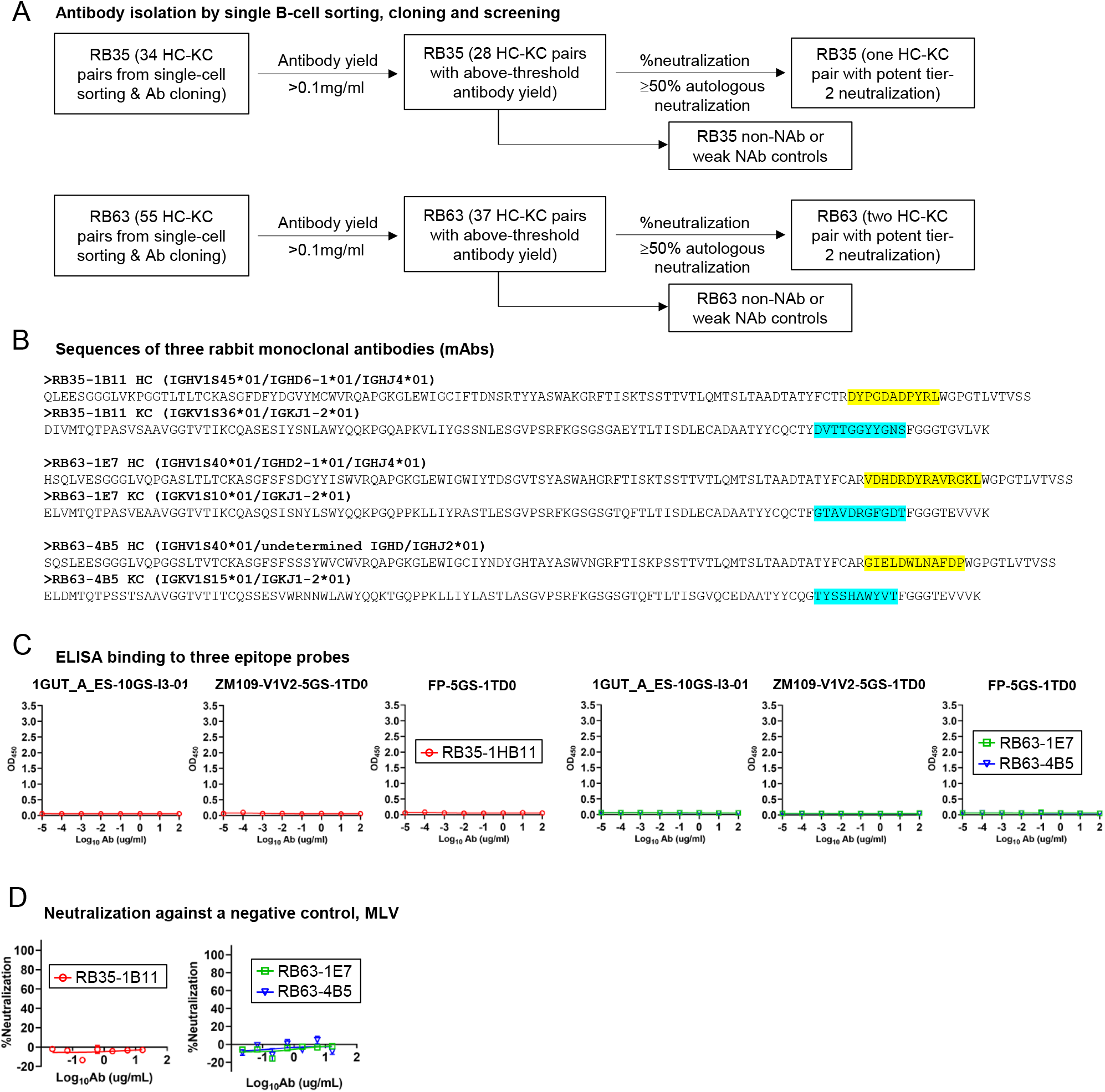
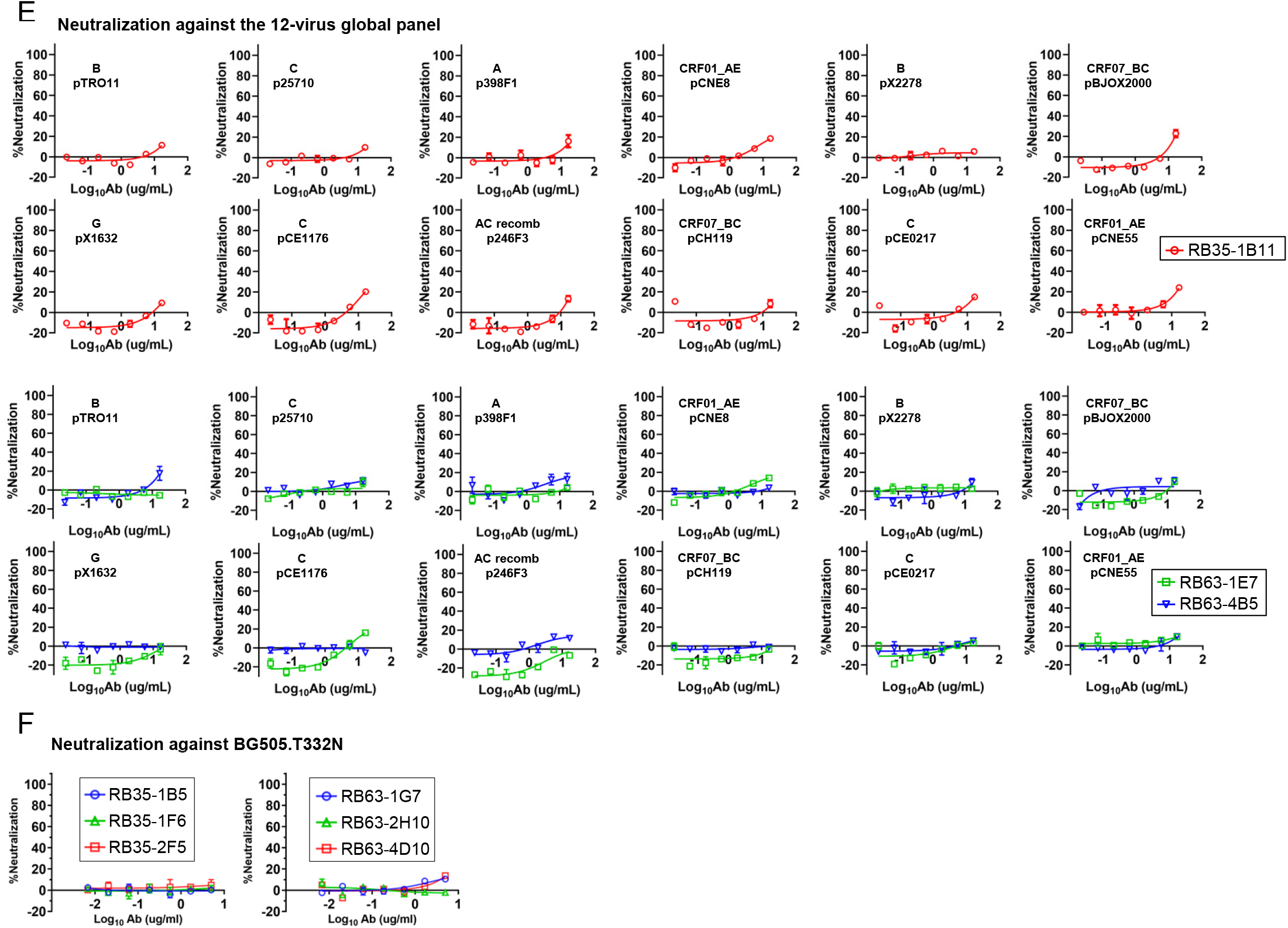
Functional evaluation of single-cell sorted rabbit mAbs. (**A**) Schematic representation of the procedure used to select functional mAbs from a rabbit immunized with BG505 gp140.664.R1 trimer (RB35) and a rabbit immunized with BG505 gp140.664.R1-FR nanoparticle (RB63). The two major selection criteria are: (1) yield ≥ 0.1mg/ml after purification and concentration, and (2) %neutralization ≥ 50% at 10ug/ml for BG505.T332N. Weak/non-NAbs matching only the first criterion may be selected for comparison. (**B**) Amino acid sequences of three rabbit NAbs identified from this screening procedure. (**C**) ELISA binding by the RB35/RB63 NAbs to an I3-01 nanoparticle presenting 24 copies of an N332 scaffold (1GUT_A_ES), a trimeric scaffold (1TD0) presenting ZM109 V1V2, and the same trimeric scaffold (1TD0) presenting fusion peptide (FP-5GS-1TD0). Antibodies were diluted to 100ug/ml and subjected to a 10-fold dilution series in the assay. (**D**) Neutralization of MLV by the RB35/RB63 NAbs. **(E)** Neutralization of all 12 isolates from a global panel by the RB35/RB63 NAbs. Antibodies were diluted to 33.3ug/ml and followed by a 3-fold dilution series in the TZM-bl assay. **(F)** ELISA binding of six non-NAbs, two from each rabbit, to BG505 UFO.664 trimer.

**fig. S6.**
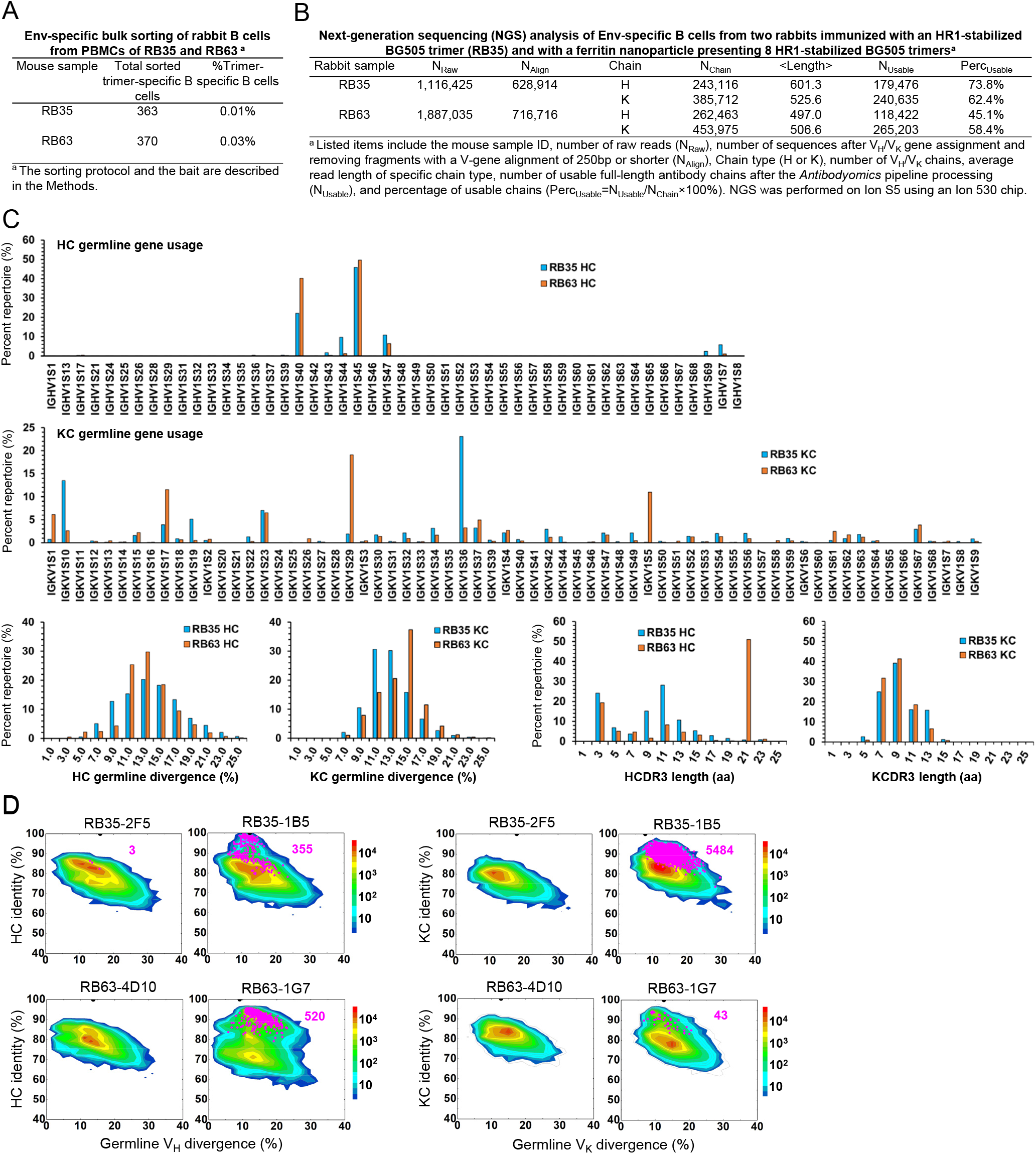
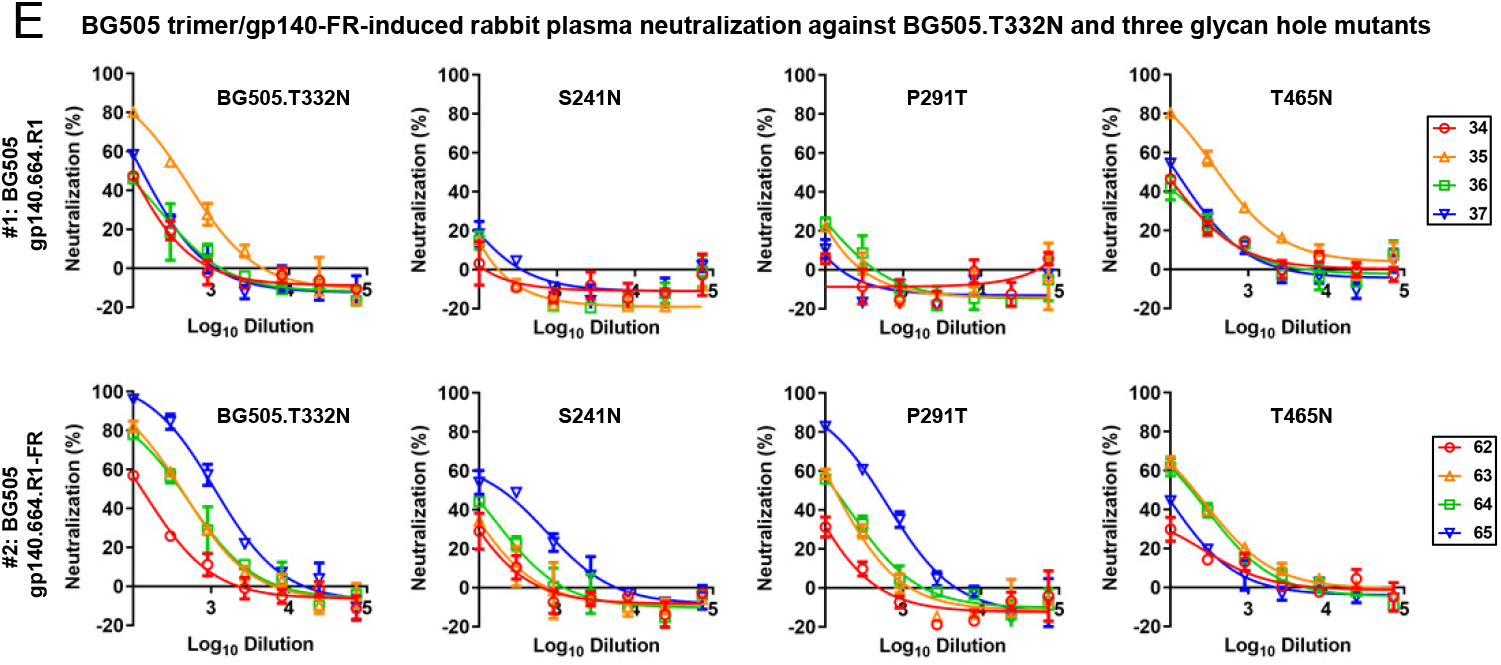
HIV-1 Env-specific sorting and NGS of rabbit B cells for antibody isolation. PBMCs from a rabbit immunized with BG505 gp140.664.R1 trimer (RB35) and a rabbit immunized with BG505 gp140.664.R1-FR10 nanoparticle (RB63) were analyzed. (**A**) Env-specific rabbit B cells obtained from bulk sorting using a biotinylated Avi-tagged BG505 gp140.664.R1 trimer probe. (**B**) Antibodyomics pipeline processing of NGS data obtained from sequencing of Env-specific rabbit B cells on the Ion S5 platform. **(C)** Quantitative B cell repertoire profiles derived from the NGS analysis of Env-specific RB35 and RB63 B cells, including HC and KC germline gene usage, somatic hypermutation (SHM), and CDR3 length. **(D)** Divergence-identity analysis of four representative non-NAbs in the context of Env-specific antibody repertories for RB35 and RB63. HC and KC sequences are plotted as a function of sequence identity to the template and sequence divergence from putative germline genes. Color coding indicates sequence density. Templates and sequences identified based on the CDR3 identity of 95% or greater are shown as black and magenta dots on the plots, respectively, with the number of sequences labeled accordingly. **(E)** Rabbit plasma neutralization from two BG505 Env-immunized rabbit groups against three glycan hole mutants with respect to BG505.T332N. The heat-inactivated rabbit plasma was diluted 100-fold as the starting point and subjected to a 3-fold dilution series in the TZM-bl assay. The %neutralization values obtained from the first dilution are reported in Fig. 3F.

**fig. S7.**
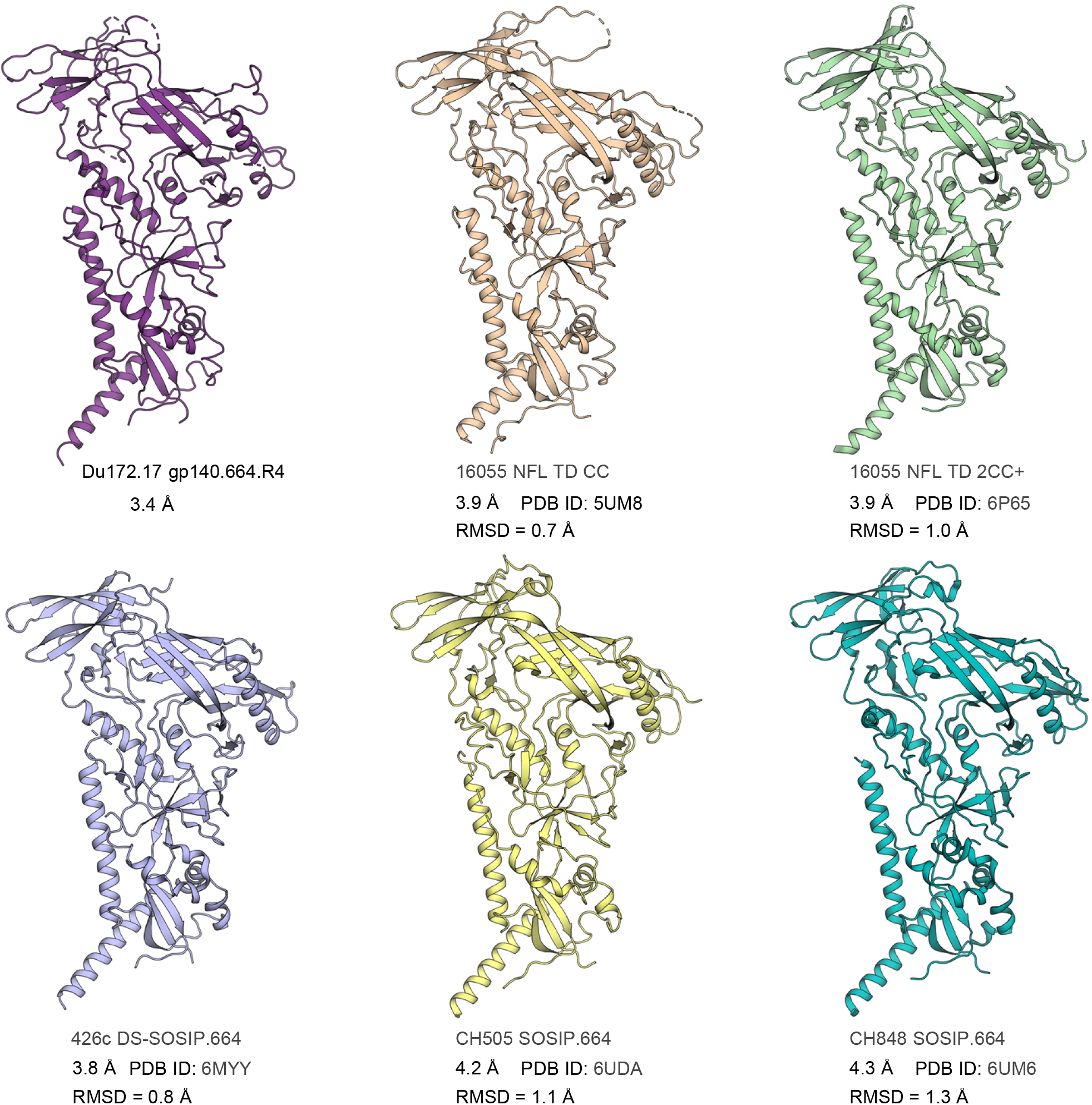
Structural comparison of HIV-1 Envs across clade C isolates. Ribbons view of two crystal structures (Du172.17 here and PDB ID: 5UM8) and four cryo-EM models (PDB IDs: 6P65, 6MYY, 6UDA, and 6UM6) obtained for clade C isolates. The Cα RMSD after superposition of each structure on Du172.17 gp140.664.R4 is shown.

**fig. S8.**
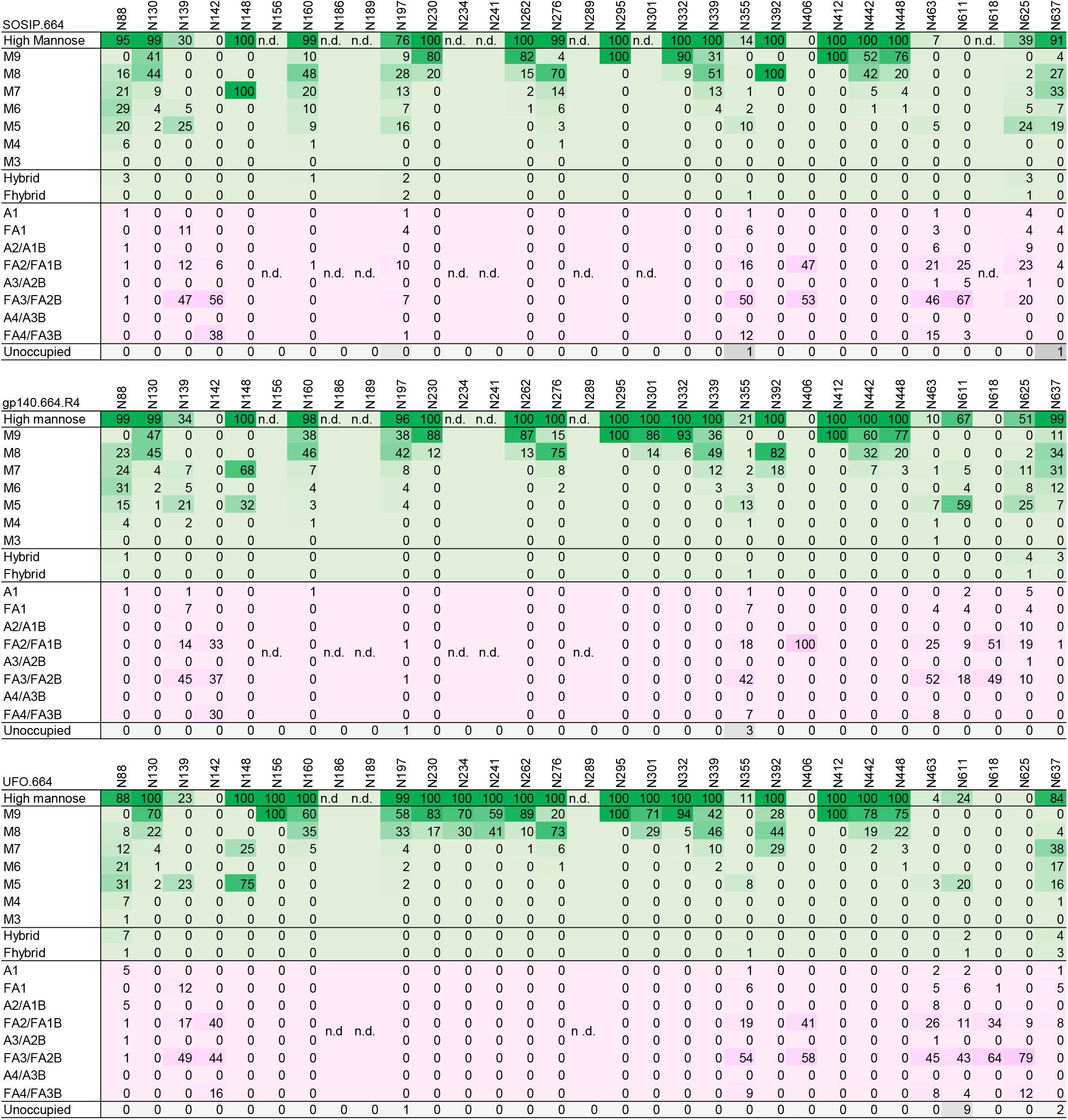
Site-specific *N*-linked glycan analysis of Du172.17 SOSIP.664, HR1-redesigned (gp140.664.R4), and UFO.664 trimers produced in HEK293F cells. Quantification of site-specific glycan occupancy and composition. The table shows the compositions found at each site. Compositions corresponding to oligomannose/hybrid-type glycans are colored in green and fully processed complex type glycans are colored in magenta. The proportion of peptides at each lacking an attached glycan are colored in grey. Oligomannose-type glycans are categorized according to the number of mannose residues, hybrid-type glycans according to the presence/absence of fucose, and complex-type glycans according to the number of processed antenna and the presence/absence of fucose.

**fig. S9.**
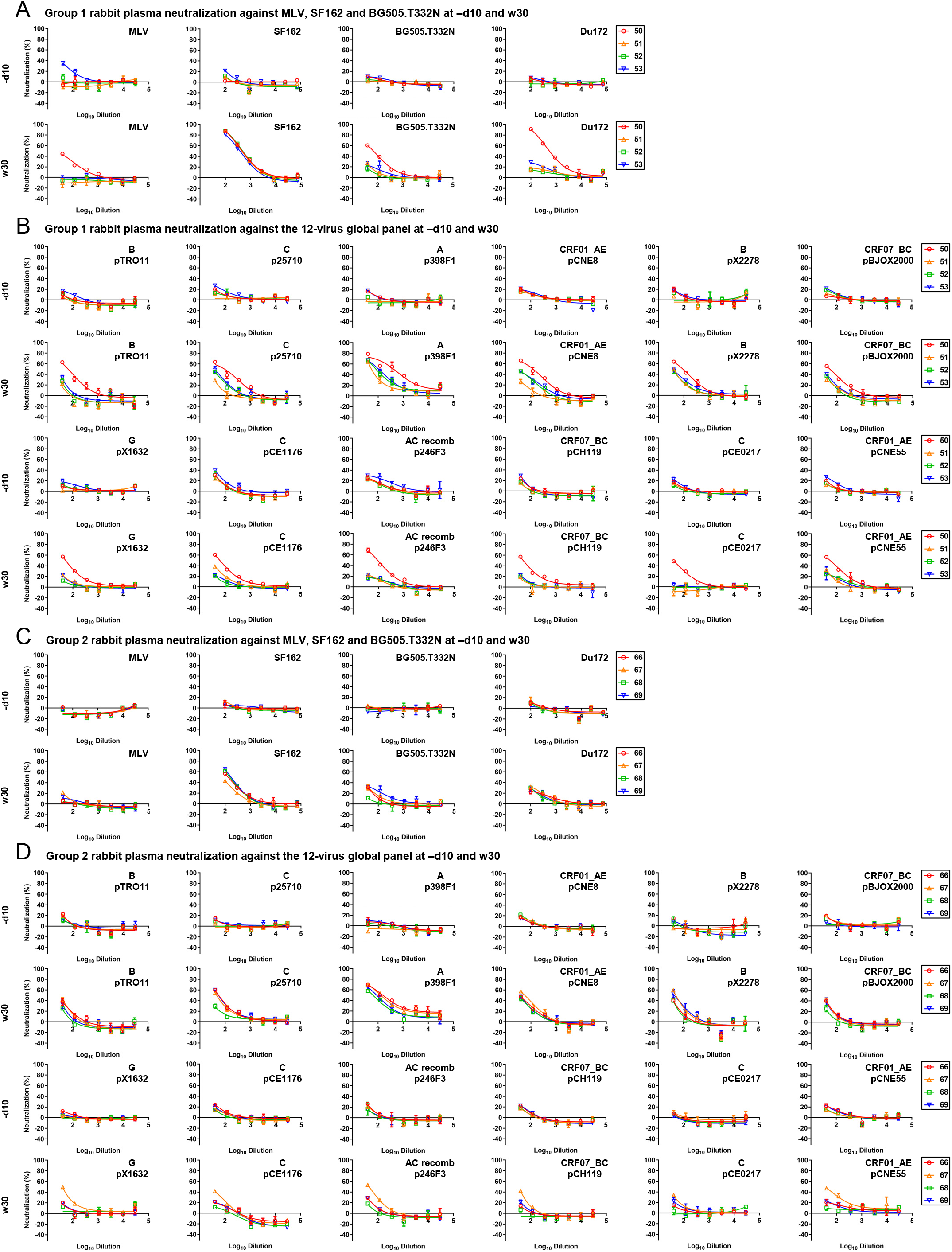
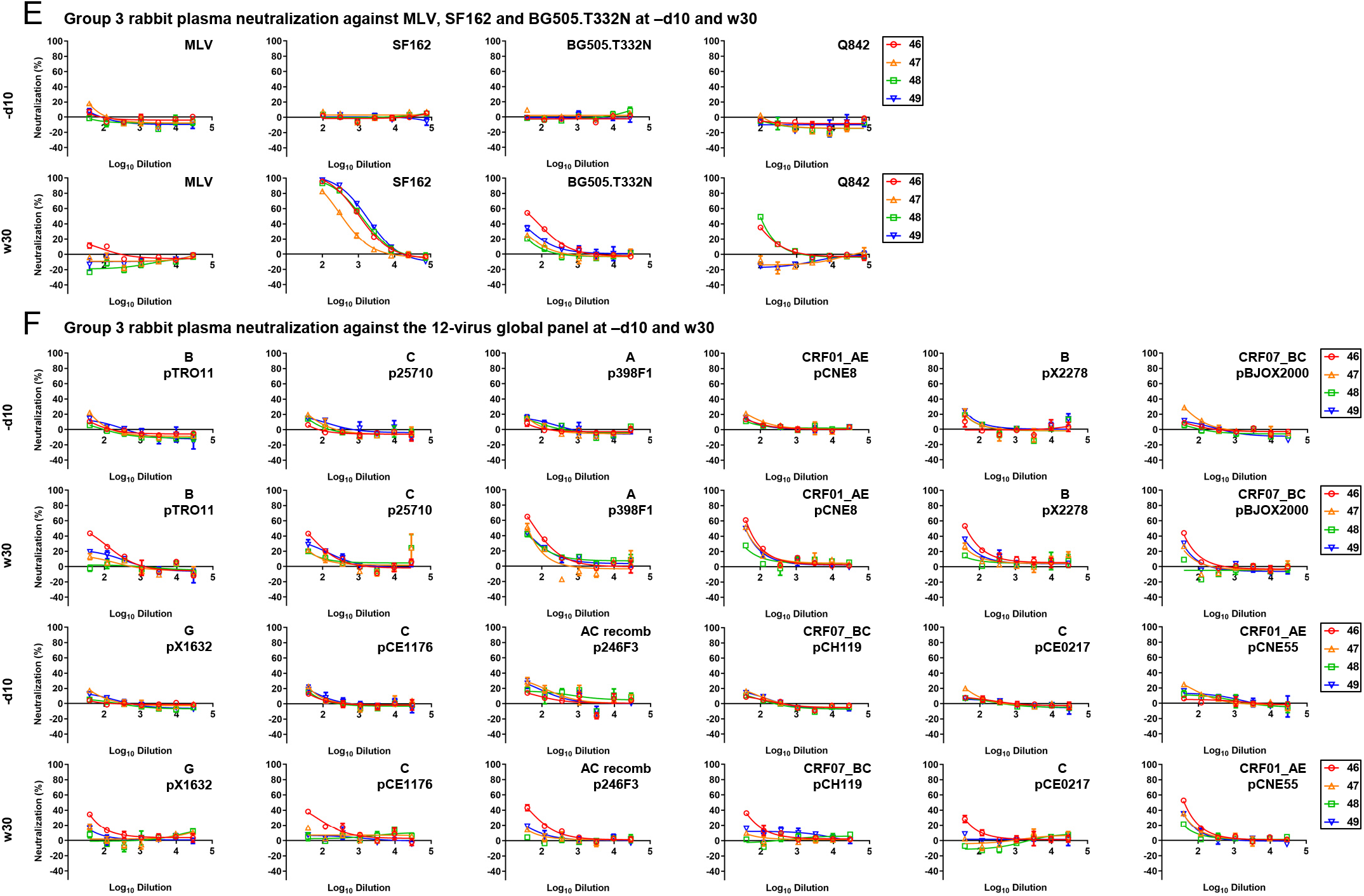
Rabbit plasma neutralization from three non-BG505 Env-immunized rabbit groups. Three groups of rabbits were immunized with Du172.17 UFO-BG trimer, gp140.664.R4-FR nanoparticle, and Q842-d12 UFO-BG trimer. (**A**) Neutralization of MLV, tier 1 clade B SF162, tier 2 clade A BG505.T332N, and tier 2 clade C Du172.17 by day -10 (-d10) and week 30 (w30) rabbit plasma from the Du172.17 trimer group. (**B**) Neutralization of all 12 isolates from a global panel by day -10 (-d10) and week 30 (w30) rabbit plasma from the Du172.17 trimer group. (**C**) Neutralization of MLV, SF162, BG505.T332N, and Du172.17 by day -10 (-d10) and week 30 (w30) rabbit plasma from the Du172.17 ferritin nanoparticle group. (**D**) Neutralization of all 12 isolates from a global panel by day -10 (-d10) and week 30 (w30) rabbit plasma from the Du172.17 ferritin nanoparticle group. (**E**) Neutralization of MLV, SF162, BG505.T332N, and tier 2 clade A Q842-d12 by day -10 (-d10) and week 30 (w30) rabbit plasma from the Q842-d12 trimer group. (**F**) Neutralization of all 12 isolates from a global panel by day -10 (-d10) and week 30 (w30) rabbit plasma from the Q842-d12 trimer group. The heat-inactivated plasma was diluted 100-fold for autologous virus and tier 1 SF162 and subjected to a 3-fold dilution series in the TZM-bl assay. To increase the sensitivity of detection, heat-inactivated plasma was diluted 40-fold for MLV and all other heterologous tier 2 isolates and followed by a 3-fold dilution series in the TZM-bl assay. ID_50_ titers for plots (A) – (F) are summarized in Fig. 4D.

**fig. S10.**
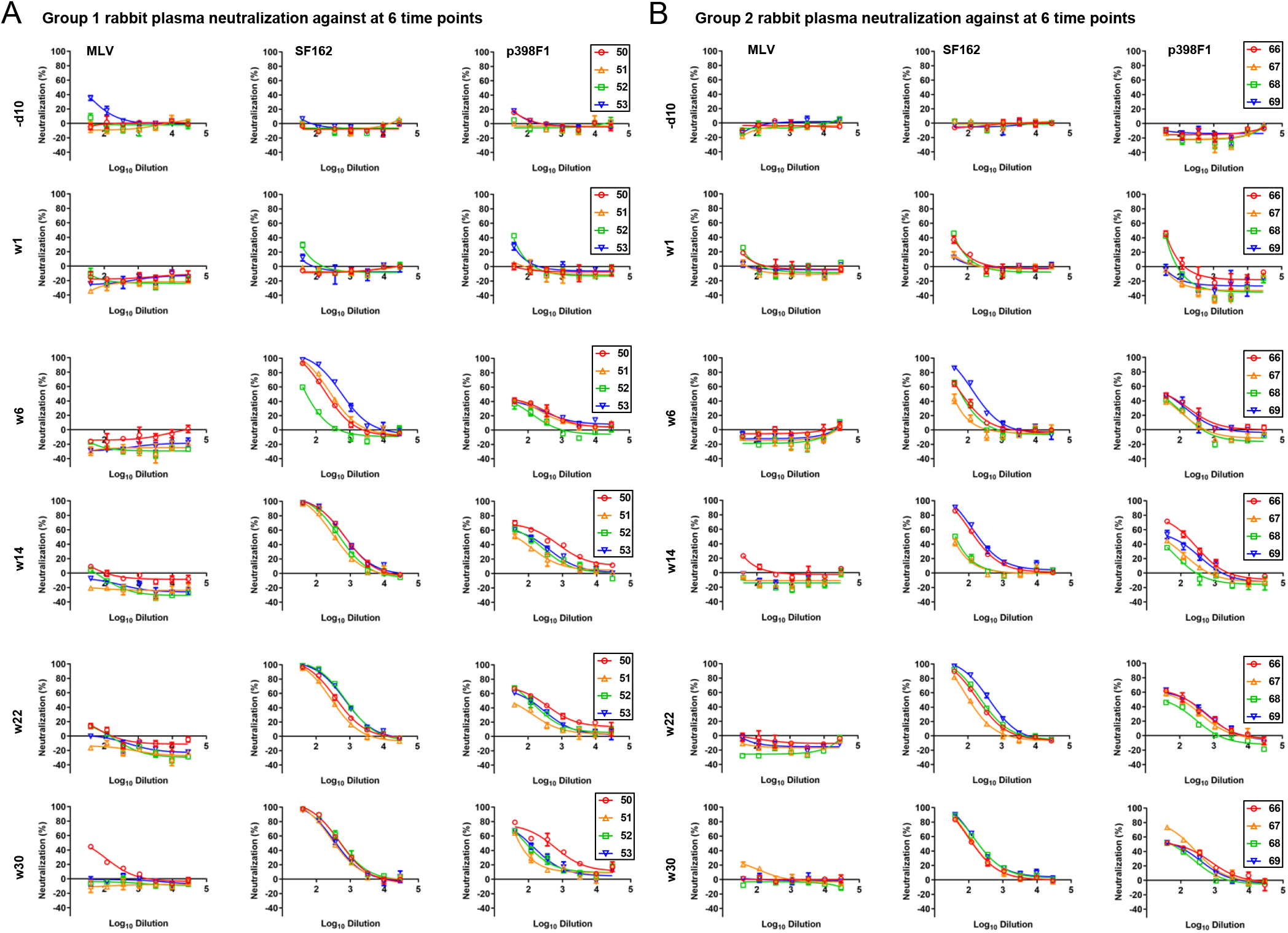
Longitudinal rabbit plasma neutralization from two clade C Du172.17 Env-immunized rabbit groups. Two rabbit groups immunized with Du172.17 UFO-BG trimer and gp140.664.R4-FR nanoparticle were analyzed. (**A**) Neutralization of MLV, tier 1 clade B SF162, and tier 2 clade A p398F1 by day -10 (-d10) and weeks 1, 6, 14, 22, and 30 rabbit plasma from the Du172.17 trimer group. **(B)** Neutralization of MLV, tier 1 clade B SF162, and tier 2 clade A p398F1 by day -10 (-d10) and weeks 1, 6, 14, 22, and 30 rabbit plasma from the Du172.17 gp140.664.R4-FR nanoparticle group. In this analysis, the heat-inactivated plasma was diluted 40-fold for both SF162 and p398F1 and then subjected to a 3-fold dilution series in the TZM-bl assay. ID_50_ titers for plots (A) – (B) are shown in Fig. 4E.

**Table S1.**
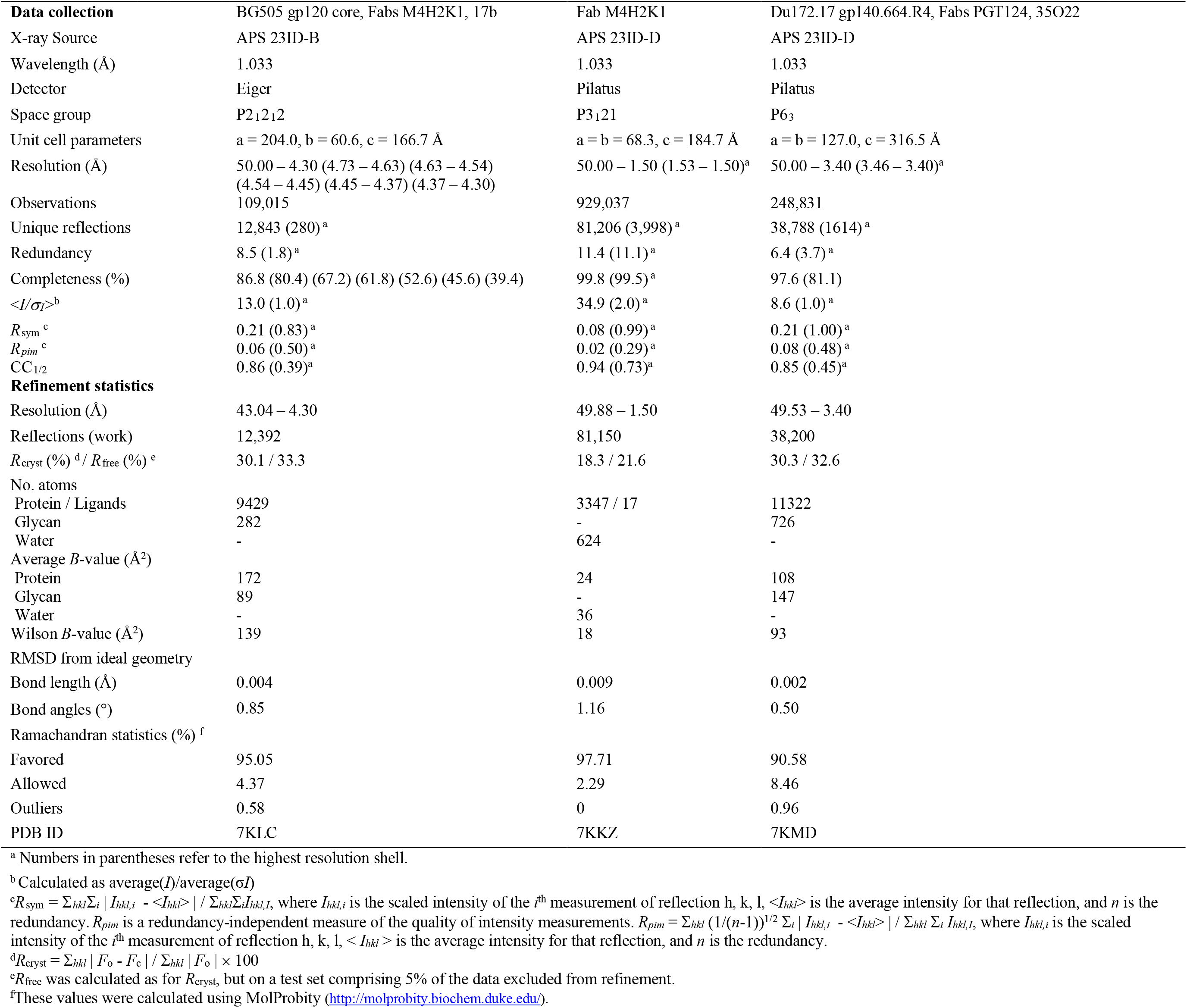
X-ray crystallographic data collection and refinement statistics.

## Acknowledgments

We thank Y. Hua, H. Tien, R. Stanfield, and Drs. X. Dai, and M. Elsliger for excellent technical help; Diffraction data were collected at the Advanced Photon Source (APS) beamline 23-IDD, and Stanford Synchrotron Radiation Lightsource (SSRL) beamline 12-2. Use of the APS was supported by the DOE, Basic Energy Sciences, Office of Science, under contract no. DE-AC02-06CH11357. Use of the SSRL was supported by the US Department of Energy, Basic Energy Sciences, Office of Science, under contract no. DE-AC02-76SF00515.

## Author contributions

Project design by S.K., X.L., I.A.W. and J.Z.; NGS, bioinformatics, antibody selection, and synthesis by X.L., L.H., C.S., and J.Z.; B cell sorting and antibody cloning by C.S., L.Z., and L.H.; antibody expression, purification, and ELISA by X.L., B.S., and T.N.; plasma and antibody neutralization by X.L., B.S., T.N., and L.H.; nsEM analysis by J.C., G.O., and A.B.W.; glycan analysis by J.D.A., and M.C.; Env expression and purification by S.K., X.L., and B.S.; x-ray crystallography by S.K. and I.A.W.; Manuscript written by S.K., L.X., I.A.W., and J.Z. All authors were asked to comment on the manuscript. The TSRI manuscript number is 30047.

## Data and materials availability

All data and code to understand and assess the conclusions of this research are available in the main text, Supplementary Materials, PDB (accession codes 7KLC, 7KKZ and 7KMD) and EMDB (accession code EMD-22999). Additional data related to this paper may be requested from the authors.

