## Supplemental Figures and Tables for "Neutralizing antibodies induced by first-generation gp41-stabilized HIV-1 envelope trimers and nanoparticles"

### Figure S1

A

**Env-specific bulk sorting of mouse splenic B cells from the I3-01 nanoparticle group [a]**

| Mouse sample | Total sorted trimer-specific B cells | %Trimer-specific B cells |
|---|---|---|
| G5-1 | 578 | 0.06% |
| G5-2 | 87 | 0.01% |
| G5-3 | 727 | 0.01% |
| G5-4 | 1064 | 0.03% |

[a] The sorting protocol and the bait are described in the Methods.

B

**Next-generation sequencing (NGS) analysis of Env-specific splenic B cells from mice immunized with a 60-meric I3-01 nanoparticle presenting the HR1-stabilized BG505 trimer [a]**

| Mouse sample | $N_{Raw}$ | $N_{Align}$ | Chain | $N_{Chain}$ | <Length> | $N_{Usable}$ | $Perc_{Usable}$ |
|---|---|---|---|---|---|---|---|
| G5-1 | 1,228,018 | 238,985 | H | 16,832 | 606.2 | 5,145 | 30.6% |
| | | | K | 222,153 | 501.4 | 221,019 | 99.5% |
| G5-2 | 297,492 | 29,218 | H | 23,880 | 605.2 | 21,124 | 88.5% |
| | | | K | 5,338 | 361.5 | 745 | 14.0% |
| G5-3 | 190,567 | 117,225 | H | 17,946 | 828.4 | 17,709 | 98.7% |
| | | | K | 99,279 | 481.2 | 99,031 | 99.8% |
| G5-4 | 315,064 | 202,970 | H | 58,370 | 620.9 | 49,753 | 85.2% |
| | | | K | 144,600 | 495.7 | 144,194 | 99.7% |

[a] Listed items include the mouse sample ID, number of raw reads ($N_{Raw}$), number of sequences after $V_H/V_K$ gene assignment and removing fragments with a V-gene alignment of 250bp or shorter ($N_{Align}$), Chain type (H or K), number of $V_H/V_K$ chains, average read length of specific chain type, number of usable full-length antibody chains after the *Antibodyomics* pipeline processing ($N_{Usable}$), and percentage of usable chains ($Perc_{Usable}=N_{Usable}/N_{Chain}\times100\%$). NGS was performed on Ion S5 using an Ion 530 chip.

C

**Amino acid sequences of consensus antibody chains identified from NGS analysis of Env-specific bulk-sorted splenic B cells [a]**

**Antibody chains from mouse 1 in the I3-01 group**

| | |
|---|---|
| >M1H1[b] | GAAGTTCAGCTATTGGAGACTGGAGGAGGCTTGGTGCAACCTGGGGGGTCACGGGGACTCTCTTGTGAAGGCTCAGGGTTTAGTTTTAGTGGCTTCTG<u>G</u>ATGAACTGGGTTCGACAGACACCTGGGAA |
| | GACCCTGGACTGGATTGGAGACATTAATTCTGATGGCACATCAATAAGCTACGCACCCTCCATAAAGGATCGATTCACTGTCTTCAGATACACTGACAAGGACACCCTGTATCTGCAGATGAACAATG |
| | TGCGATCTGAAGACACAGCCCCGTATTTCTGTATGAGGGGGTTTTACTTACTGGGGCCAAGGCTTACTGGGGCCAAGGGACTCTGGTCACTGTCTCTACA |
| >M1H2 | GAAGTGAACCTTGAGGAGTCTGGAGGAGGCTTGGTGCAACCTGGAGGATCCATGAAACTCTCCTGTGTTGCCTCTGGATTCACTATCAGTAACTACTGGATGAACTGGGTCCGCCAGTCTCCAGAGAA |
| | GGGGCTTGAGTGGGTTGCTGAAATTAGATTGAAAGCTAATAATGATGCAACACATTATGCGGAGTCTGTGAAAGGGAGGTTCACCATCTCAAGAGATGATTCCAAAAATAGTGTCTACCTGCAAATGA |
| | ACAACTTAAGAGCTGAAGACACTGCCAATTATTACTGTACCAGGCCCGGTTACTACGGCTACTATGCTATGGACCAGTGGGGTCAAGGAACCTCAGTCACCGTCTCCTCA |
| >M1H3 | CAGGTTCAGTTGCAGCAGTCTGGCGCTGAGTTGGTGAAACCTGGGGCCTCAGTGAAGATATCCTGCAAGGCTTCTGGCTACACCTTCACTGACCGTGCTATTCACTGGGTGAAACAGAAGCCTGAACA |
| | GGGCCTGGAATGGATTGGATATATAGTTCCCGGAAATAGTGATATTAAGTACAGTGAGAAGTTCAAGGGCAAGGCCACACTGACTGCAGACAAATCCTCCACCACTGCCTACATGCAGGTCAACAGCC |
| | TGACATCTGAGGACTCTGCAGTGTATTTCTGTAATTGCTATGATTACGACGACGGCTATTGGGGTCAAGGAACCTCAGTCACCGTCTCCTCA |
| >M1K1 | GACATTGTGCTGACCCAATCTCCAGCTTCTTTGGCTGTGTCTCTAGGACAGAGGGCCACCATCTCCTGCAGAGCCAGCGAAAGTGTTGATAATTATGGCATTAGTTTTATGAACTGGTTCCAACAGAA |
| | ACCAGGACAGCCACCCAAACTCCTCATCTATGGTGCATCCAACCAAGGATCCGGGGTCCCTGCCAGGTTTAGTGGCAGTGGGTCTGGGACGGATTTCAGCCTCAACATCCATCCTATGGAGGAGGATG |
| | ATACTGCAATGTTTTTCTGTCAGCAAAGTAAGGAGGTTCCGTACACGTTCGGAGGGGGGACCAAGGTGGAAATAAAA |
| >M1K2 | GACATTGTGATGACCCAGTCTCACAAATTCATGTCCACATCAGTAGGAGACAGGGTCAGCATCACCTGCAAGGCCAGTCAGGATGTGAGTACTGCTGTAGCCTGGTATCAACAAAAACCAGGGCAATC |
| | TCCTAAACTACTGTTTTACTGGACATCCGCCCGGCACACTGGAGTCCCTGATCGCTTCACAGGCAGTGGATCTGGGACAGATTATACTCTCACCATCAACAATGTAATGGCTGAAGACCTGGCACTTT |
| | ATTACTGTCAACAACATTATAGCACTCCGTGGACGTTCGGTGGAGGCACCAAGCTGGAGATCAAA |
| >M1K3 | CAAATTGTTCTCACCCAGTCTCCACGAATCATGTCTGCATCTCCAGGGGAGAGGGTCACCGTGACCTGCAGTGCCAGCTCAAGTGTAAGTTACATGTCCTGGTACCAGCAGAAGCCAGGATCCTCCCC |
| | CAGACTCCTGATTTTTGACACATCCAACCTGGCTTCTGGAGTCCCTTTTCGCTTCAGTGGCAGTGGGTCTGGGACCTCTTACTCTCTCACAATCAGCCGAATGGAGGCTGAAGATGCTGCCACTTATT |
| | GCTGCCAGCAGTGGAGTCGTTACCCATTCACGTTCGGCTCGGGGACAAAGTTGGAAATAAAA |

**Antibody chains from mouse #4 in the I3-01 group**

| | |
|---|---|
| >M4H1[b] | CAGGTCCAGTTGCAGCAGTCTGGAGCTGACCTGGTCAGGCCTGGGACTTCATTGAAGAAGTCCTCCAAGGTTTCTGGCTACACCTTCACTAACTACTGGATAGGTTGGGTAAAGCAGAGGCCTGGACA |
| | TGGCCTTGAGTGGATTGGAGATGTTTACCCTGGAGACGGTTTTACTCAGAACAATGAGAAGTTCAAGGACAAGGCCACACTGACTGCAGACAAATCCTCCAGCACATCCTACAGG<u>C</u>A<u>G</u>CTCAGCAGTC |
| | TGACATCTGAGGAATCTGCGGTCCATTA<u>C</u>TGTTCGACACCTACGGTAGTGCCTGACTACTGGGGCCAAGGCACCACTCTCACAGTCTCCTCA |
| >M4H2 | GAAGTGAAGCTTGAGGAGTCCGGAGGAGGCTTGGTGCAACCTGGAGGATCCATGAAACTCTCCTGTGTTGCCTCTGGAATCACTTTCAGTAACTCCTGGATGAGCTGGGTCCGCCAGTCTCCAGAGAA |
| | GGGGCTTGAGTGGGTTGCTGAAATTAGATTGAAAGCTCAAAATTATGCAACACATTATGCGGCGTCTGTGAAAGGGAGGTTCACCATCTCAAGAGATGACTCCAAAAGTAGTGTCTACCTACAAATGA |
| | ACAACTTAAGACCTGAAGACACTGGCATCTATTACTGTACCACCCCACTGGGAGGCTATTTTGATATGGACTACTGGGGTCAAGGAACCTCACTCACCGTCTCCTCA |
| >M4K1 | GACATTGTGCTGACCCAATCTCCAGCTTCTTTGGCTGTGTCTCTAGGGCAGAGGGCCACCATCTCCTGCAGAGCCAGCGAAAGTGTTGATATTTATGGCATTAGTTTTATGAACTGGTTCCAACAGAG |
| | ACCAGGACAGCCACCCAAACTCCTCATCTATGCTGCATCCAACCGAGGATCCGGGGTCCCTGCCAGGTTTAGTGGCAGTGGGTCTGGGACAGACTTCAGCCTCAACATCCATCCTATGGAGGAGGATG |
| | ATACTGCAATGTATTTCTGTCAGCAAAGTAAGGAGGTTCCGTGGACGTTCGGTGGAGGCACCAAGCTGGAAATCAAA |
| >M4K2 | CAAATTGTTCTCACCCAGTCTCCAGCAATCATGTCTGCATCTCCAGGGGAGAAGGTCACCATGACCTGCAGTGCCAGCTCAAGTGTAAGTTACATGTCTTGGTACCAGCAGAAGCCAGGATCCTCCCC |
| | CCGACTCCTGATTTATGACACATCCGACCTGGCTTCTGGAGTCCCTTTTCGCTTCAGTGGCAGTGGGTCTGGGACCTCTTACTCTCTCACAATCAGCCGAATGGAGGCTGAAGATGCTGCCACTTATT |
| | ACTGCCAACAGTGGGATCCTTACCCGCTCACGTTCGGTTCTGGGACCAAGCTGGAGCTGAAA |
| >M4K3 | GACATTGTGATGTCACAGTCTCCATCCTCCCTAGCTGTGTCAGTTGGAGAGAAGGTTACTATGAGCTGCAAGTCCAGTCAGAGCCTTTTATATAGTAGCAATCAAAAGAACTACTTGGCCTGGTACCA |
| | GCAGAAACCAGGGCAGTCTCCTAAACTGCTGATTTACTGGGCATCCACTAGGGAATCTGGGGTCCCTGATCGCTTCACAGGCAGTGGATCTGGGACAGATTTCACTCTCACCATCAGCAGTGTGAAGG |
| | CTGAAGACCTGGCAGTTTATTACTGTCAGCAATATTATAGCTATCCGCTCACGTTCGGTGCTGGGACCAAGCTGGAGCTGAAA |

[a] The antibody chain sequences were identified from the NGS of Env-specific splenic B cells using a clustering algorithm and consensus as described in detail in the Methods.
[b] Nucleotide that was modified to remove a stop codon is colored in red and underscored.

**Amino acid sequences of consensus antibody chains identified from NGS analysis of Env-specific bulk-sorted splenic B cells [a]**

**Antibody chains from mouse 1 in the I3-01 group**

| | |
|---|---|
| >M1H1[b] | EVQLLETGGGLVQPGGSRGLSCEGSGFSFSGFWMNWVRQTPGKTLDWIGDINSDGTSISYAPSIKDRFTVFRYTDKDTLYLQMNNVRSEDTAPYFCMRGFYLLGPRLTGAKGLWSLSL |
| >M1H2 | EVNLEESGGGLVQPGGSMKLSCVASGFTISNYWMNWVRQSPEKGLEWVAEIRLKANNDATHYAESVKGRFTISRDDSKNSVYLQMNNLRAEDTANYYCTRPGYYGYYAMDQWGQGTSVTVSS |
| >M1H3 | QVQLQQSGAELVKPGASVKISCKASGYTFTDRAIHWVKQKPEQGLEWIGYIVPGNSDIKYSEKFKGKATLTADKSSTTAYMQVNSLTSEDSAVYFCNCYDYDDGYWGQGTSVTVSS |
| >M1K1 | DIVLTQSPASLAVSLGQRATISCRASESVDNYGISFMNWFQQKPGQPPKLLIYGASNQGSGVPARFSGSGSGTDFSLNIHPMEEDDTAMFFCQQSKEVPYTFGGGTKVEIK |
| >M1K2 | DIVMTQSHKFMSTSVGDRVSITCKASQDVSTAVAWYQQKPGQSPKLLFYWTSARHTGVPDRFTGSGSGTDYTLTINNVMAEDLALYYCQQHYSTPWTFGGGTKLEIK |
| >M1K3 | QIVLTQSPRIMSASPGERVTVTCSASSSVSYMSWYQQKPGSSPRLLIFDTSNLASGVPFRFSGSGSGTSYSLTISRMEAEDAATYCCQQWSRYPFTFGSGTKLEIK |

**Antibody chains from mouse 4 in the I3-01 group**

| | |
|---|---|
| >M4H1 | QVQLQQSGADLVRPGTSLKKSSKVSGYTFTNYWIGWVKQRPGHGLEWIGDVYPGDGFTQNNEKFKDKATLTADKSSSTSYRQLSSLTSEESAVHYCSTPTVVPDYWGQGTTLTVSS |
| >M4H2 | EVKLEESGGGLVQPGGSMKLSCVASGITFSNSWMSWVRQSPEKGLEWVAEIRLKAQNYATHYAASVKGRFTISRDDSKSSVYLQMNNLRPEDTGIYYCTTPLGGYFDMDYWGQGTSLTVSS |
| >M4K1 | DIVLTQSPASLAVSLGQRATISCRASESVDIYGISFMNWFQQRPGQPPKLLIYAASNRGSGVPARFSGSGSGTDFSLNIHPMEEDDTAMYFCQQSKEVPWTFGGGTKLEIK |
| >M4K2 | QIVLTQSPAIMSASPGEKVTMTCSASSSVSYMSWYQQKPGSSPRLLIYDTSDLASGVPFRFSGSGSGTSYSLTISRMEAEDAATYYCQQWDPYPLTFGSGTKLELK |
| >M4K3 | DIVMSQSPSSLAVSVGEKVTMSCKSSQSLLYSSNQKNYLAWYQQKPGQSPKLLIYWASTRESGVPDRFTGSGSGTDFTLTISSVKAEDLAVYYCQQYYSYPLTFGAGTKLELK |

[a] The amino acid sequences were translated from the nucleotide sequences using TRANSEQ.
[b] HCDR3 and framework 4 (FR4) region missing the “WGXG” motif and “VSS” motifs are highlighted in gray shading.

D

**Nucleotide sequences of two antibodies identified from splenic B cells of mouse 4 (M4) in the I3-01 group by Env-specific single-cell sorting [a]**

```
>M4-Ab3H CAGGTGCAGCTGCAGCAGCCTGGAGGAGGCTCGGTGCAACCTGGAGGATCCATGAAACTCTCCTGTGTTGCCTCTGGATTCACCTTCAGTAATTCCTGGATGAACTGGGTCCGCCAGTCTCCAGAGAA
         GGGGCTTGAGTGGGTTGCTGAAATTCGATTGAAAGTTCATAATTATGCAACACATTATGCGGAGTCTGTGAAAGGGAGGTTCACCATCTCAAGAGATGATTCCAAAAGTAGTGTCTACCTGCAAATGA
         TCAACTTAAGACCAGAAGACACTGGCATTTATTATTGTACTACCCCACTGGGTGGCTACTTTCCTATGGACTACTGGGGTCAAGGAACCACTCTCACAGTCTCCTCA
>M4-Ab3K GACATTGTGCTGACCCAATCTCCAACTTCTTTGGCTGTGTCTCTAGGGCAGAGGGCCACCATCTCCTGCAGAGCCAGCGAAAGTGTTGATAATTATGGCGTTAGTTTTATGAACTGGTTCCAACAGAA
         ACCAGGACGGCCACCCAAACTCCTCATCTATGCTGCATCCAAGCAAGGATCCGGGGTCCCTGCCAGGTTTAGTGGCAGTGGGTCTGGGACAGATTTCAGCCTCAACATCCATCCAATGGAGGAGGATG
         ATATTGCAATGTATTTCTGTCAGCAAAATAAGGAGCTTCCGTGGACGTTCGGTGGAGGCACCAAGCTGGAAATCAAA
>M4-Ab9H AGGGTGCAGCTGCAGCAGTCTTGTGGAGGCTTGGTGCAACCTGGAGGATCCATGAAACTCTCCTGCGTTGCCTCTGGAATCACTTTCAGTAACTCCTGGATGAACTGGGTCCGCCAGTCTCCAGAGAA
         GGGGCTTGAGTGGGTTGCTGAAATTAGATTGAAAGTTAATAATTATGCAACACATTATGCGGAGTCTGTGAAAGGGAGGTTCACCATCTCAAGAGATGATTCCAAAAGGAGTGTCTACCTGCAAATGA
         ACAACTTAAGAGCTGAAGACACTGGCATTTATTACTGTACCACCCCACTGGGTGGCTACTATGCTGTGGACTACTGGGGTCAAGGAGCCACTCTCACAGTCTCCTCA
>M4-Ab9K GACATCCAGATGATTCAGTCTCCAGCTTCTTTGGCTGTGTCTCTAGGGCAGAGGGCCACCATCTCCTGCAGAGCCAGCGAAAGTGTTGATAATTATGGCATTAGTTTTATGAACTGGTTCCAACAGAA
         ACCAGGACAGCCACCCAAACTCCTCATCTATGCTGCATCCAACCAAGGATCCGGGGTCCCTGCCAGGTTTAGTGGCAGTGGGTCTGGGACAGACTTCAGCCTCAACATCCATCCTATGGAGGAGGATG
         ATACTGCAATGTATTTCTGTCAGCAAAGTAAGGAGGTTCCGTGGACGTTCGGTGGAGGCACCAAGCTGGAAATCAAA
```

[a] The antibody chain sequences were identified from Env-specific single-cell sorting, PCR and cloning as described in the Methods.

**Amino acid sequences of two antibodies identified from splenic B cells of mouse 4 (M4) in the I3-01 group by Env-specific single-cell sorting [a]**

```
>M4-Ab3H    QVQLQQPGGGSVQPGGSMKLSCVASGFTFSNSWMNWVRQSPEKGLEWVAEIRLKVHNYATHYAESVKGRFTISRDDSKSSVYLQMINLRPEDTGIYYCTTPLGGYFPMDYWGQGTTLTVSS
>M4-Ab3K    DIVLTQSPTSLAVSLGQRATISCRASESVDNYGVSFMNWFQQKPGRPPKLLIYAASKQGSGVPARFSGSGSGTDFSLNIHPMEEDDIAMYFCQQNKELPWTFGGGTKLEIK
>M4-Ab9H    RVQLQQSCGGLVQPGGSMKLSCVASGITFSNSWMNWVRQSPEKGLEWVAEIRLKVNNYATHYAESVKGRFTISRDDSKRSVYLQMNNLRAEDTGIYYCTTPLGGYYAVDYWGQGATLTVSS
>M4-Ab9K    DIQMIQSPASLAVSLGQRATISCRASESVDNYGISFMNWFQQKPGQPPKLLIYAASNQGSGVPARFSGSGSGTDFSLNIHPMEEDDTAMYFCQQSKEVPWTFGGGTKLEIK
```

[a] The amino acid sequences were translated from the nucleotide sequences using TRANSEQ.

**fig S1. HIV-1 Env-specific sorting and NGS of mouse splenic B cells for antibody isolation.** Mice immunized with BG505 gp140.664.R1-PADRE-I3-01 nanoparticle (see ref. 37) were analyzed in this study. (**A**) Env-specific mouse splenic B cells obtained from bulk sorting using a biotinylated Avi-tagged BG505 gp140.664.R1 trimer probe. (**B**) Antibodyomics pipeline processing of NGS data obtained from sequencing of Env-specific mouse splenic B cells on the Ion S5 platform. **(C)** Nucleotide and amino acid sequences of consensus antibody heavy and κ-light chains (HC and KC) identified from NGS analysis of Env-specific splenic B cells from mice 1 and 4 (M1 and M4) in the I3-01 nanoparticle group. (**D**) Nucleotide and amino acid sequences of two antibodies, Ab3 and Ab9, identified by single-cell sorting and antibody cloning from M4 splenic B cells. Ab3 and Ab9 use the same germline genes as the NGS-derived NAb, M4H2K1.

**Figure S2**

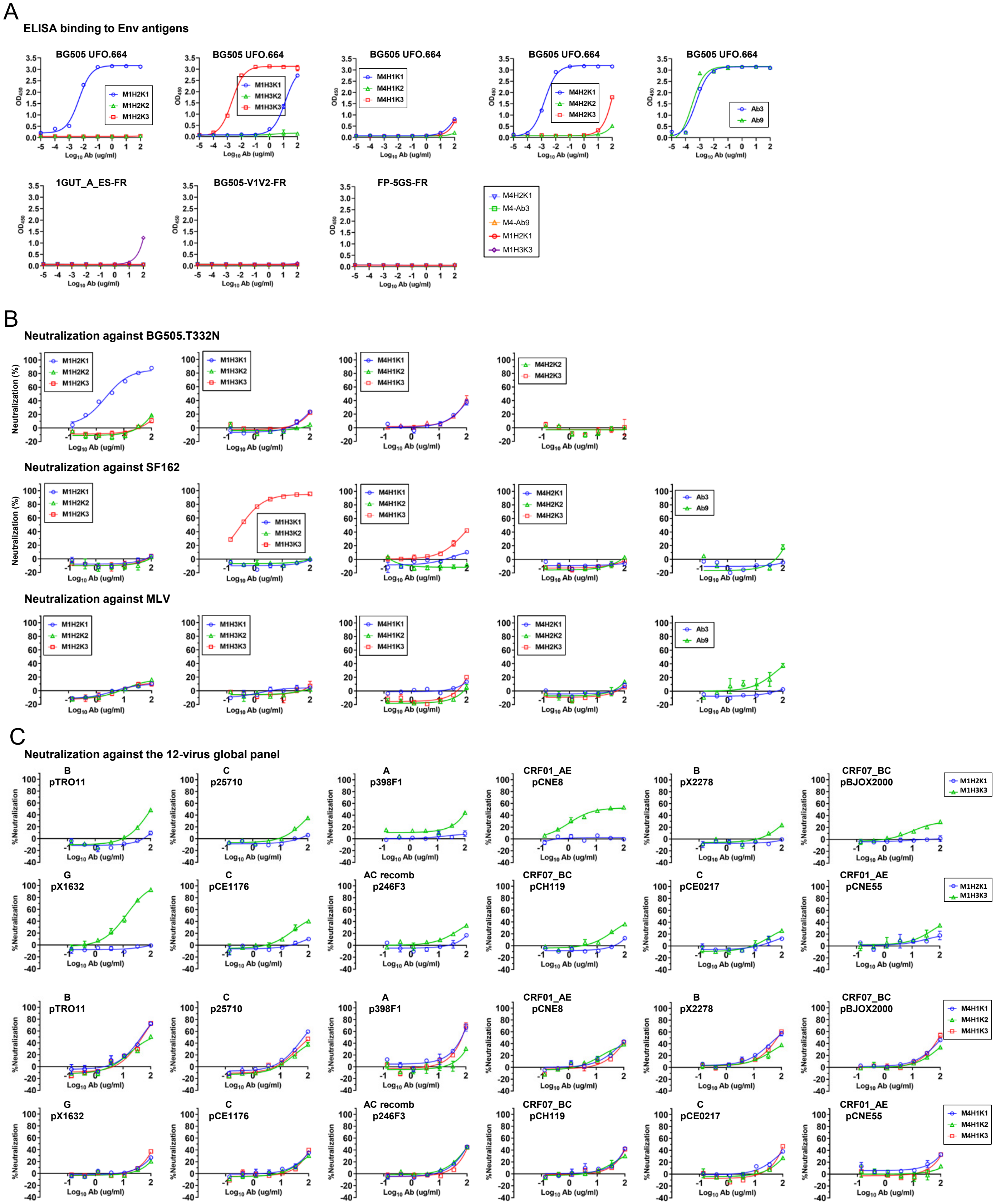
A
ELISA binding to Env antigens
BG505 UFO.664
M1H2K1
M1H2K2
M1H2K3
M1H3K1
M1H3K2
M1H3K3
M4H1K1
M4H1K2
M4H1K3
M4H2K1
M4H2K2
M4H2K3
Ab3
Ab9
1GUT_A_ES-FR
BG505-V1V2-FR
FP-5GS-FR
M4-Ab3
M4-Ab9
OD450
Log10 Ab (ug/ml)
B
Neutralization against BG505.T332N
Neutralization against SF162
Neutralization against MLV
Neutralization (%)
C
Neutralization against the 12-virus global panel
B
pTRO11
C
p25710
A
p398F1
CRF01_AE
pCNE8
pX2278
CRF07_BC
pBJOX2000
G
pX1632
pCE1176
AC recomb
p246F3
pCH119
pCE0217
pCNE55
%Neutralization

### Figure S2

C (continued)

**Neutralization against the 12-virus global panel**

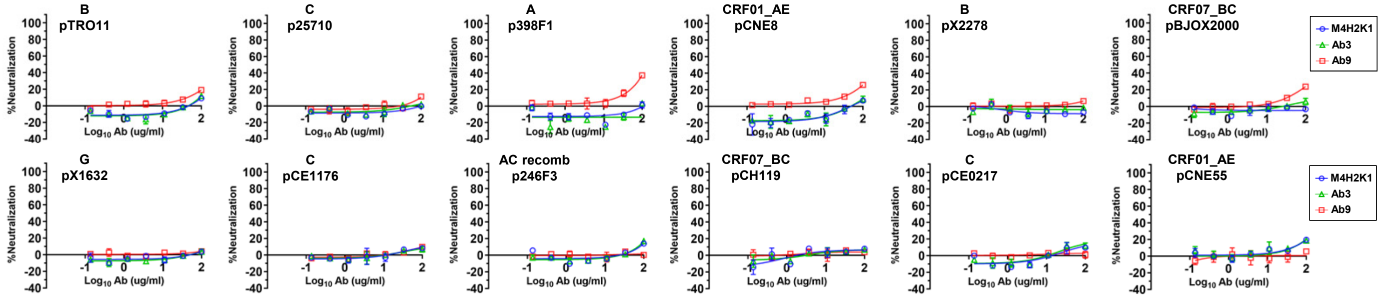


**fig S2. Functional evaluation of NGS and single-cell-derived mouse mAbs.** (**A**) ELISA binding of mouse mAbs to Env antigens including BG505 UFO.664 trimer (top panel) and three individual epitope probes including 1GUT_A_ES-FR (N332 supersite), BG505 V1V2-FR (V1V2 apex), and FP-5GS-FR (fusion peptide), which are all ferritin nanoparticles. **(B)** Neutralization of autologous tier 2 clade A BG505.T332N by mouse mAbs. **(C)** Neutralization of all 12 isolates from a global panel by mouse mAbs.

### Figure S3

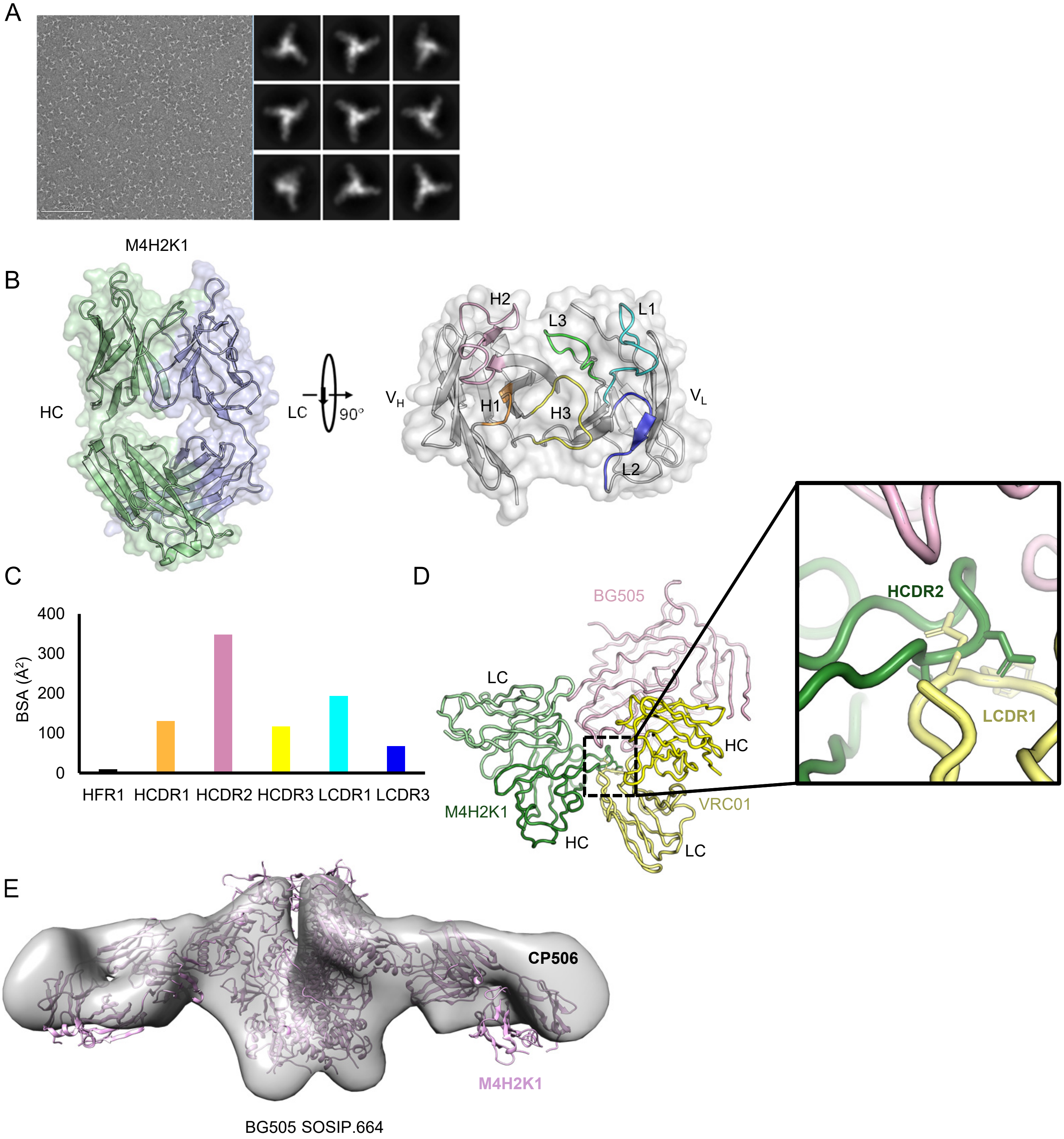


**fig. S3. Structural characterization of the NGS-derived mouse NAb, M4H2K1**. (**A**) Negative-stain EM (nsEM) analysis of mouse NAb M4H2K1 in complex with BG505 UFO.664 Env trimer. Left: EM micrograph; Right: 2D class averages. (**B**) The unbound structure of M4H2K1 in a ribbons model within the molecular surface. Left: side view; Right: top view. The H/LCDR loops are labeled on the structure. (**C**) Buried surface area (Å$^2$) of the CDR loops and FRs of M4H2K1 Fab when bound to BG505 gp120 core. (**D**). Superimposition of VRC01 (yellow) Fab-bound BG505 SOSIP with M4H2K1 (green) Fab-bound BG505 core (pink). The right inset shows a clash of M4H2K1 HCDR2 with VRC01 LCDR1. (**E**) Comparison of the mode of recognition for M4H2K1 and CP506 when bound to the BG505 SOSIP.664 Env trimer.

### Figure S4

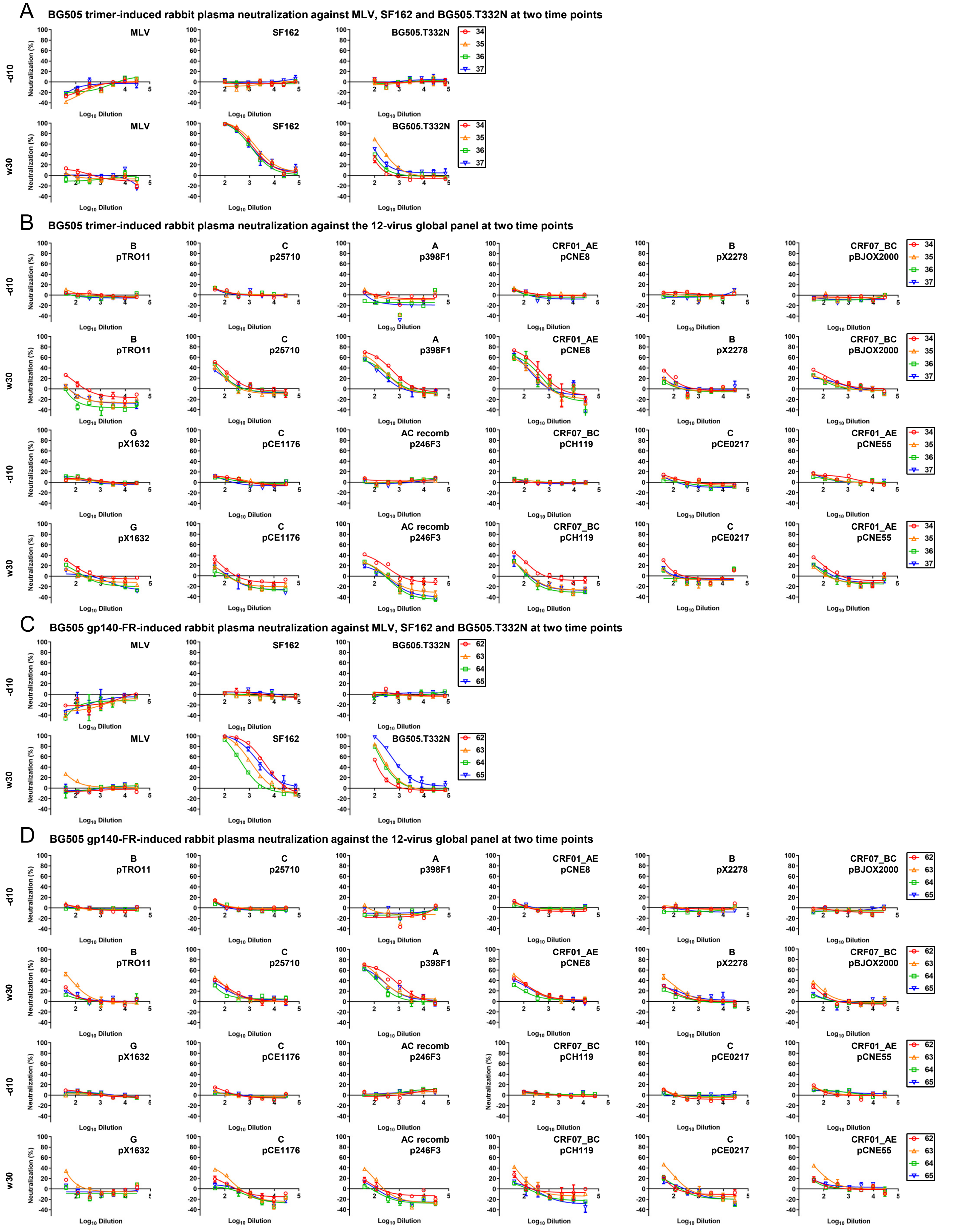
A
BG505 trimer-induced rabbit plasma neutralization against MLV, SF162 and BG505.T332N at two time points
MLV
SF162
BG505.T332N
-d10
w30
34
35
36
37
Neutralization (%)
Log10 Dilution
B
BG505 trimer-induced rabbit plasma neutralization against the 12-virus global panel at two time points
B pTRO11
C p25710
A p398F1
CRF01_AE pCNE8
B pX2278
CRF07_BC pBJOX2000
G pX1632
C pCE1176
AC recomb p246F3
CRF07_BC pCH119
C pCE0217
CRF01_AE pCNE55
C
BG505 gp140-FR-induced rabbit plasma neutralization against MLV, SF162 and BG505.T332N at two time points
62
63
64
65
D
BG505 gp140-FR-induced rabbit plasma neutralization against the 12-virus global panel at two time points

### Figure S4

**fig. S4. Rabbit plasma neutralization from two BG505 Env-immunized rabbit groups.** In the previous study (see ref. 37), two groups of rabbits were immunized with BG505 gp140.664.R1 trimer and its ferritin nanoparticle. (**A**) Neutralization of MLV, tier 1 clade B SF162, and tier 2 clade A BG505.T332N by day -10 (-d10) and week 30 (w30) rabbit plasma from the BG505 trimer group. (**B**) Neutralization of 12 isolates in the global panel by day -10 (-d10) and week 30 (w30) rabbit plasma from the BG505 trimer group. (**C**) Neutralization of MLV, SF162, and BG505.T332N by day -10 (-d10) and week 30 (w30) rabbit plasma from the BG505 ferritin nanoparticle group. (**D**) Neutralization of 12 isolates in the global panel by day -10 (-d10) and week 30 (w30) rabbit plasma from the BG505 ferritin nanoparticle group. The heat-inactivated rabbit plasma was diluted 100-fold for autologous tier 2 BG505.T332N and tier 1 SF162 and subjected to a 3-fold dilution series in the TZM-bl assay. To increase the sensitivity of detection, heat-inactivated plasma was diluted 40-fold for MLV and all 12 isolates from a global panel and followed by a 3-fold dilution series in the TZM-bl assays. $ID_{50}$ titers for plots (A) – (D) are shown in Fig. 3B.

### Figure S5

A **Antibody isolation by single B-cell sorting, cloning and screening**

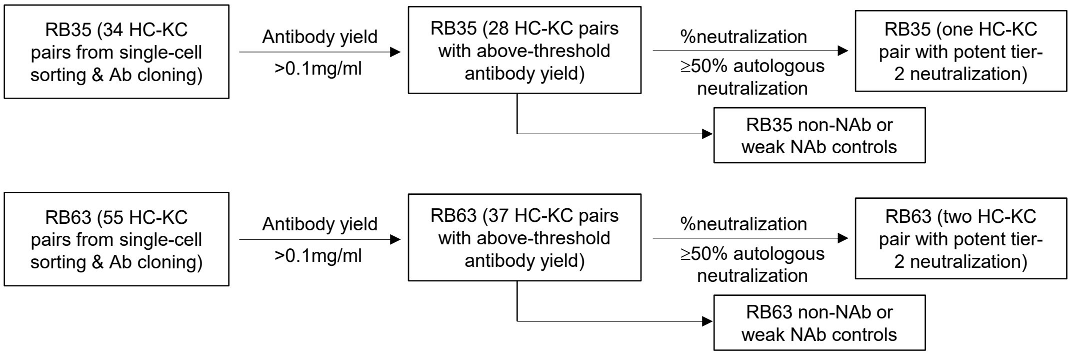


B **Sequences of three rabbit monoclonal antibodies (mAbs)**

**>RB35-1B11 HC (IGHV1S45*01/IGHD6-1*01/IGHJ4*01)**
QLEESGGGLVKPGGTLTLTCKASGFDFYDGVYMCWVRQAPGKGLEWIGCIFTDNSRTYYASWAKGRFTISKTSSTTVTLQMTSLTAADTATYFCTRDYPGDADPYRLWGPGTLVTVSS
**>RB35-1B11 KC (IGKV1S36*01/IGKJ1-2*01)**
DIVMTQTPASVSAAVGGTVTIKCQASESIYSNLAWYQQKPGQAPKVLIYGSSNLESGVPSRFKGSGSGAEYTLTISDLECADAATYYCQCTYDVTTGGYYGNSFGGGTGVLVK

**>RB63-1E7 HC (IGHV1S40*01/IGHD2-1*01/IGHJ4*01)**
HSQLVESGGGLVQPGASLTLTCKASGFSFSDGYYISWVRQAPGKGLEWIGWIYTDSGVTSYASWAHGRFTISKTSSTTVTLQMTSLTAADTATYFCARVDHDRDYRAVRGKLWGPGTLVTVSS
**>RB63-1E7 KC (IGKV1S10*01/IGKJ1-2*01)**
ELVMTQTPASVEAAVGGTVTIKCQASQSISNYLSWYQQKPGQPPKLLIYRASTLESGVPSRFKGSGSGTQFTLTISDLECADAATYYCQCTFGTAVDRGFGDTFGGGTEVVVK

**>RB63-4B5 HC (IGHV1S40*01/undetermined IGHD/IGHJ2*01)**
SQSLEESGGGLVQPGGSLTVTCKASGFSFSSSYWVCWVRQAPGKGLEWIGCIYNDYGHTAYASWVNGRFTISKPSSTTVTLQMTSLTAADTATYFCARGIELDWLNAFDPWGPGTLVTVSS
**>RB63-4B5 KC (IGKV1S15*01/IGKJ1-2*01)**
ELDMTQTPSSTSAAVGGTVTITCQSSESVWRNNWLAWYQQKTGQPPKLLIYLASTLASGVPSRFKGSGSGTQFTLTISGVQCEDAATYYCQGTYSSHAWYVTFGGGTEVVVK

C **ELISA binding to three epitope probes**

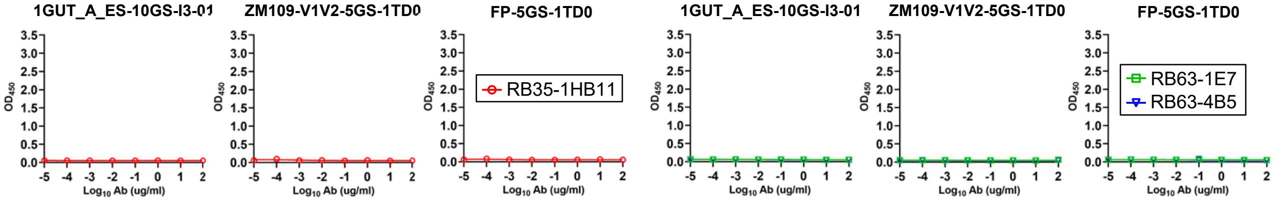


D **Neutralization against a negative control, MLV**

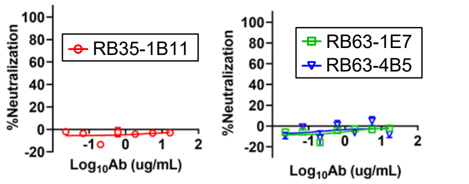

E

**Neutralization against the 12-virus global panel**

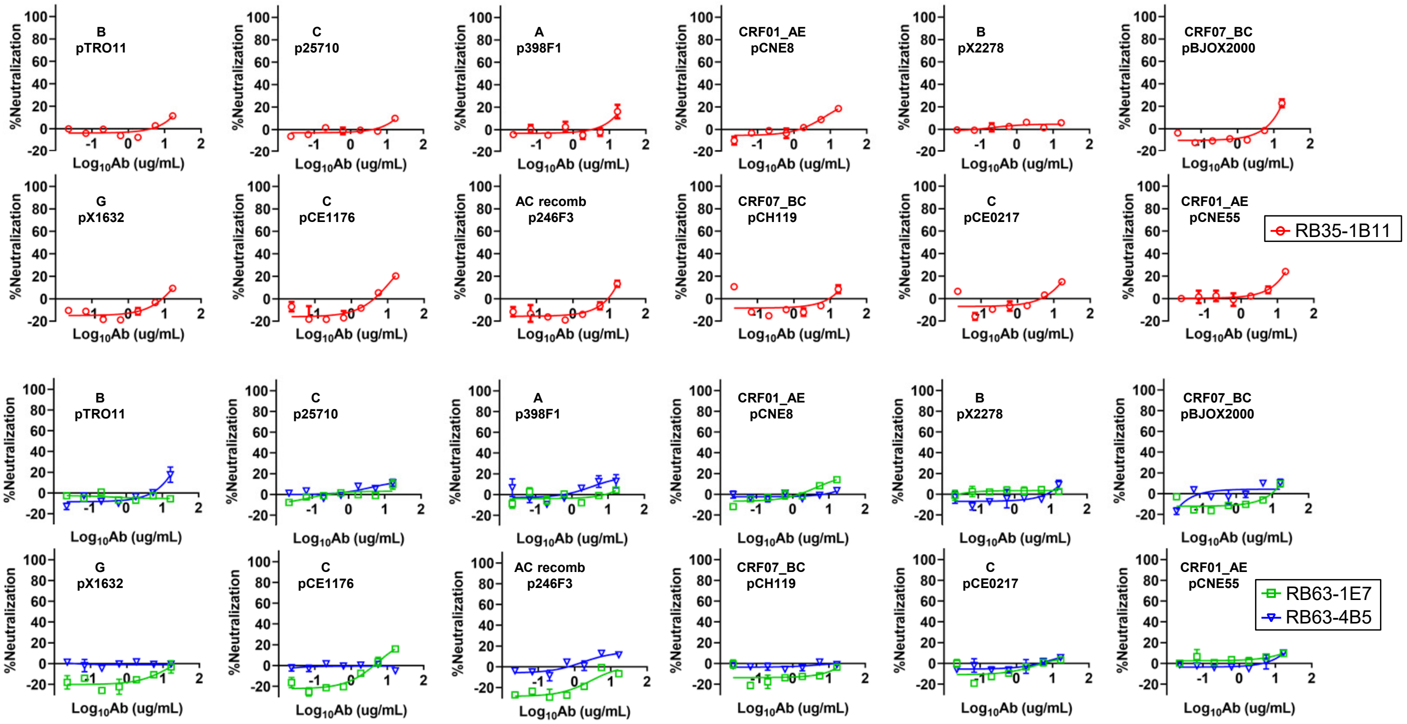


F

**Neutralization against BG505.T332N**

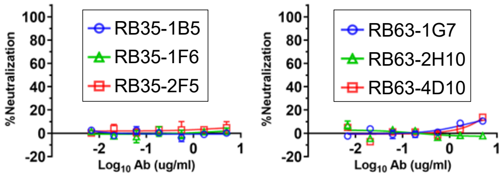


**fig. S5. Functional evaluation of single-cell sorted rabbit mAbs**. (**A**) Schematic representation of the procedure used to select functional mAbs from a rabbit immunized with BG505 gp140.664.R1 trimer (RB35) and a rabbit immunized with BG505 gp140.664.R1-FR nanoparticle (RB63). The two major selection criteria are: (1) yield ≥ 0.1mg/ml after purification and concentration, and (2) %neutralization ≥ 50% at 10ug/ml for BG505.T332N. Weak/non-NAbs matching only the first criterion may be selected for comparison. (**B**) Amino acid sequences of three rabbit NAbs identified from this screening procedure. (**C**) ELISA binding by the RB35/RB63 NAbs to an I3-01 nanoparticle presenting 24 copies of an N332 scaffold (1GUT_A_ES), a trimeric scaffold (1TD0) presenting ZM109 V1V2, and the same trimeric scaffold (1TD0) presenting fusion peptide (FP-5GS-1TD0). Antibodies were diluted to 100ug/ml and subjected to a 10-fold dilution series in the assay. (**D**) Neutralization of MLV by the RB35/RB63 NAbs. **(E)** Neutralization of all 12 isolates from a global panel by the RB35/RB63 NAbs. Antibodies were diluted to 33.3ug/ml and followed by a 3-fold dilution series in the TZM-bl assay. **(F)** ELISA binding of six non-NAbs, two from each rabbit, to BG505 UFO.664 trimer.

### Figure S6

A

**Env-specific bulk sorting of rabbit B cells from PBMCs of RB35 and RB63 [a]**

| Mouse sample | Total sorted trimer-specific B cells | %Trimer-specific B cells |
|---|---|---|
| RB35 | 363 | 0.01% |
| RB63 | 370 | 0.03% |

[a] The sorting protocol and the bait are described in the Methods.

B

**Next-generation sequencing (NGS) analysis of Env-specific B cells from two rabbits immunized with an HR1-stabilized BG505 trimer (RB35) and with a ferritin nanoparticle presenting 8 HR1-stabilized BG505 trimers[a]**

| Rabbit sample | $N_{Raw}$ | $N_{Align}$ | Chain | $N_{Chain}$ | <Length> | $N_{Usable}$ | $Perc_{Usable}$ |
|---|---|---|---|---|---|---|---|
| RB35 | 1,116,425 | 628,914 | H | 243,116 | 601.3 | 179,476 | 73.8% |
| | | | K | 385,712 | 525.6 | 240,635 | 62.4% |
| RB63 | 1,887,035 | 716,716 | H | 262,463 | 497.0 | 118,422 | 45.1% |
| | | | K | 453,975 | 506.6 | 265,203 | 58.4% |

[a] Listed items include the mouse sample ID, number of raw reads ($N_{Raw}$), number of sequences after $V_H/V_K$ gene assignment and removing fragments with a V-gene alignment of 250bp or shorter ($N_{Align}$), Chain type (H or K), number of $V_H/V_K$ chains, average read length of specific chain type, number of usable full-length antibody chains after the *Antibodyomics* pipeline processing ($N_{Usable}$), and percentage of usable chains ($Perc_{Usable}=N_{Usable}/N_{Chain}\times100\%$). NGS was performed on Ion S5 using an Ion 530 chip.

C

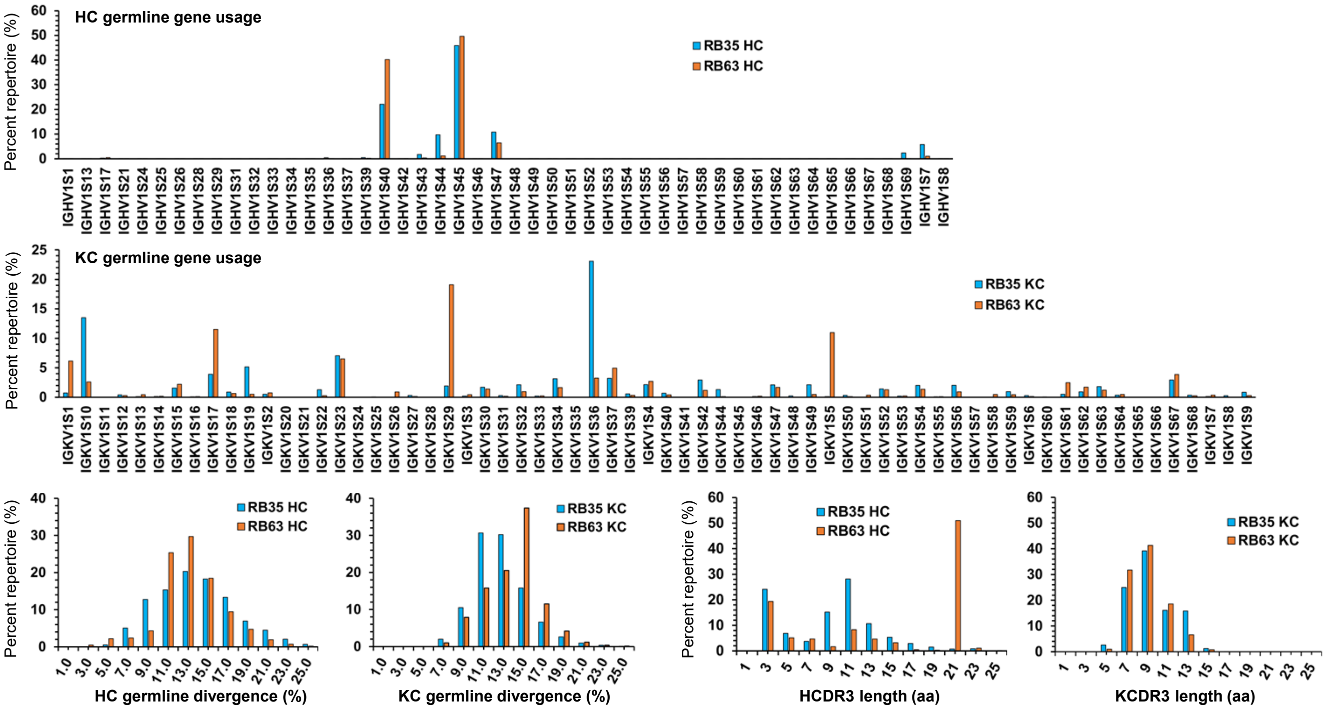


D

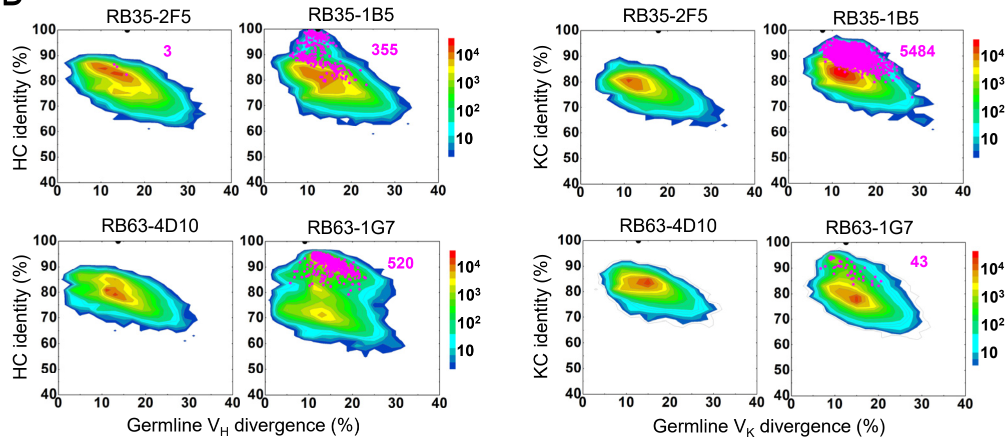

### Figure S6

E

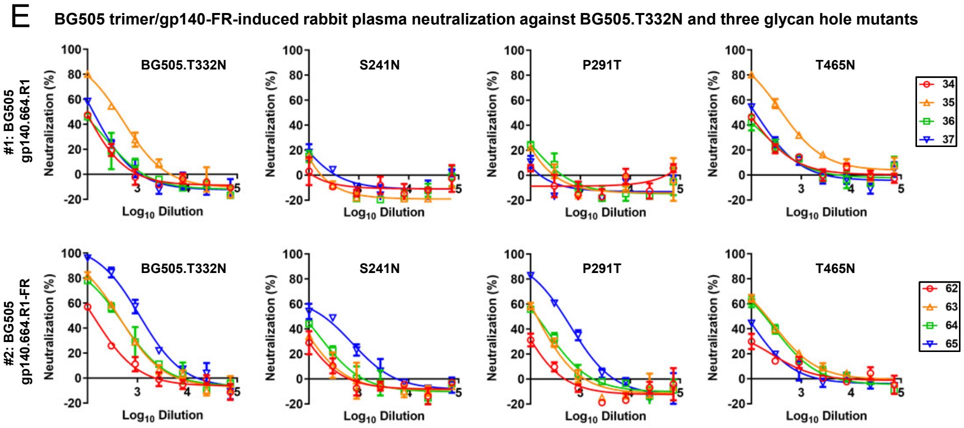


**fig. S6. HIV-1 Env-specific sorting and NGS of rabbit B cells for antibody isolation.** PBMCs from a rabbit immunized with BG505 gp140.664.R1 trimer (RB35) and a rabbit immunized with BG505 gp140.664.R1-FR10 nanoparticle (RB63) were analyzed. (**A**) Env-specific rabbit B cells obtained from bulk sorting using a biotinylated Avi-tagged BG505 gp140.664.R1 trimer probe. (**B**) Antibodyomics pipeline processing of NGS data obtained from sequencing of Env-specific rabbit B cells on the Ion S5 platform. **(C)** Quantitative B cell repertoire profiles derived from the NGS analysis of Env-specific RB35 and RB63 B cells, including HC and KC germline gene usage, somatic hypermutation (SHM), and CDR3 length. **(D)** Divergence-identity analysis of four representative non-NAbs in the context of Env-specific antibody repertories for RB35 and RB63. HC and KC sequences are plotted as a function of sequence identity to the template and sequence divergence from putative germline genes. Color coding indicates sequence density. Templates and sequences identified based on the CDR3 identity of 95% or greater are shown as black and magenta dots on the plots, respectively, with the number of sequences labeled accordingly. **(E)** Rabbit plasma neutralization from two BG505 Env-immunized rabbit groups against three glycan hole mutants with respect to BG505.T332N. The heat-inactivated rabbit plasma was diluted 100-fold as the starting point and subjected to a 3-fold dilution series in the TZM-bl assay. The %neutralization values obtained from the first dilution are reported in Fig. 3F.

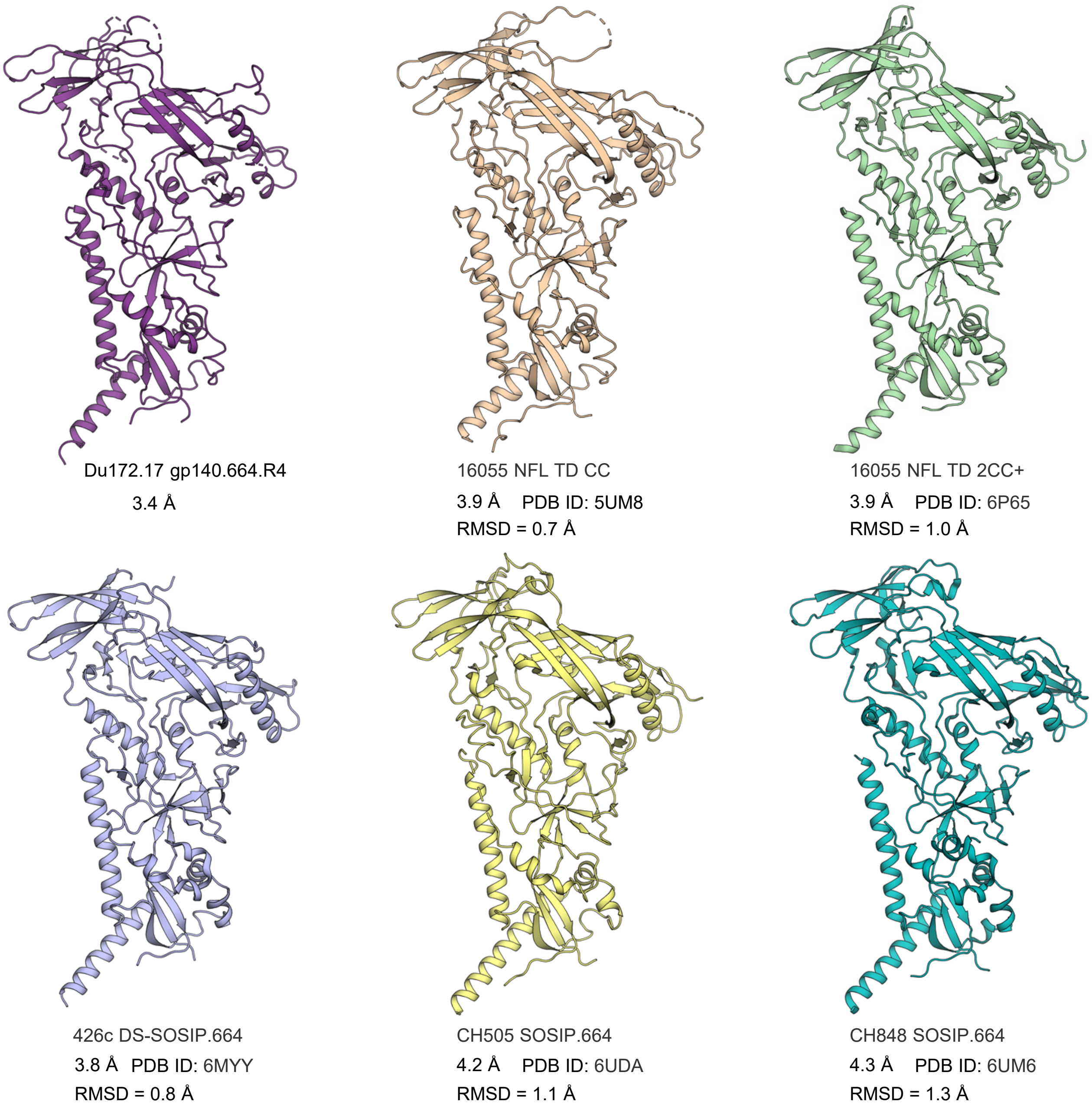


**fig. S7. Structural comparison of HIV-1 Envs across clade C isolates.** Ribbons view of two crystal structures (Du172.17 here and PDB ID: 5UM8) and four cryo-EM models (PDB IDs: 6P65, 6MYY, 6UDA, and 6UM6) obtained for clade C isolates. The C$\alpha$ RMSD after superposition of each structure on Du172.17 gp140.664.R4 is shown.

### Figure S8

| SOSIP.664 | N88 | N130 | N139 | N142 | N148 | N156 | N160 | N186 | N189 | N197 | N230 | N234 | N241 | N262 | N276 | N289 | N295 | N301 | N332 | N339 | N355 | N392 | N406 | N412 | N442 | N448 | N463 | N611 | N618 | N625 | N637 |
|---|---|---|---|---|---|---|---|---|---|---|---|---|---|---|---|---|---|---|---|---|---|---|---|---|---|---|---|---|---|---|---|
| High Mannose | 95 | 99 | 30 | 0 | 100 | n.d. | 99 | n.d. | n.d. | 76 | 100 | n.d. | n.d. | 100 | 99 | n.d. | 100 | n.d. | 100 | 100 | 14 | 100 | 0 | 100 | 100 | 100 | 7 | 0 | n.d. | 39 | 91 |
| M9 | 0 | 41 | 0 | 0 | 0 | | 10 | | | 9 | 80 | | | 82 | 4 | | 100 | | 90 | 31 | 0 | 0 | 0 | 100 | 52 | 76 | 0 | 0 | | 0 | 4 |
| M8 | 16 | 44 | 0 | 0 | 0 | | 48 | | | 28 | 20 | | | 15 | 70 | | 0 | | 9 | 51 | 0 | 100 | 0 | 0 | 42 | 20 | 0 | 0 | | 2 | 27 |
| M7 | 21 | 9 | 0 | 0 | 100 | | 20 | | | 13 | 0 | | | 2 | 14 | | 0 | | 0 | 13 | 1 | 0 | 0 | 0 | 5 | 4 | 0 | 0 | | 3 | 33 |
| M6 | 29 | 4 | 5 | 0 | 0 | | 10 | | | 7 | 0 | | | 1 | 6 | | 0 | | 0 | 4 | 2 | 0 | 0 | 0 | 1 | 1 | 0 | 0 | | 5 | 7 |
| M5 | 20 | 2 | 25 | 0 | 0 | | 9 | | | 16 | 0 | | | 0 | 3 | | 0 | | 0 | 0 | 10 | 0 | 0 | 0 | 0 | 0 | 5 | 0 | | 24 | 19 |
| M4 | 6 | 0 | 0 | 0 | 0 | | 1 | | | 0 | 0 | | | 0 | 1 | | 0 | | 0 | 0 | 0 | 0 | 0 | 0 | 0 | 0 | 0 | 0 | | 0 | 0 |
| M3 | 0 | 0 | 0 | 0 | 0 | | 0 | | | 0 | 0 | | | 0 | 0 | | 0 | | 0 | 0 | 0 | 0 | 0 | 0 | 0 | 0 | 0 | 0 | | 0 | 0 |
| Hybrid | 3 | 0 | 0 | 0 | 0 | | 1 | | | 2 | 0 | | | 0 | 0 | | 0 | | 0 | 0 | 0 | 0 | 0 | 0 | 0 | 0 | 0 | 0 | | 3 | 0 |
| Fhybrid | 0 | 0 | 0 | 0 | 0 | | 0 | | | 2 | 0 | | | 0 | 0 | | 0 | | 0 | 0 | 1 | 0 | 0 | 0 | 0 | 0 | 0 | 0 | | 1 | 0 |
| A1 | 1 | 0 | 0 | 0 | 0 | | 0 | | | 1 | 0 | | | 0 | 0 | | 0 | | 0 | 0 | 1 | 0 | 0 | 0 | 0 | 0 | 1 | 0 | | 4 | 0 |
| FA1 | 0 | 0 | 11 | 0 | 0 | | 0 | | | 4 | 0 | | | 0 | 0 | | 0 | | 0 | 0 | 6 | 0 | 0 | 0 | 0 | 0 | 3 | 0 | | 4 | 4 |
| A2/A1B | 1 | 0 | 0 | 0 | 0 | | 0 | | | 0 | 0 | | | 0 | 0 | | 0 | | 0 | 0 | 0 | 0 | 0 | 0 | 0 | 0 | 6 | 0 | | 9 | 0 |
| FA2/FA1B | 1 | 0 | 12 | 6 | 0 | n.d. | 1 | n.d. | n.d. | 10 | 0 | n.d. | n.d. | 0 | 0 | n.d. | 0 | n.d. | 0 | 0 | 16 | 0 | 47 | 0 | 0 | 0 | 21 | 25 | n.d. | 23 | 4 |
| A3/A2B | 0 | 0 | 0 | 0 | 0 | | 0 | | | 0 | 0 | | | 0 | 0 | | 0 | | 0 | 0 | 0 | 0 | 0 | 0 | 0 | 0 | 1 | 5 | | 1 | 0 |
| FA3/FA2B | 1 | 0 | 47 | 56 | 0 | | 0 | | | 7 | 0 | | | 0 | 0 | | 0 | | 0 | 0 | 50 | 0 | 53 | 0 | 0 | 0 | 46 | 67 | | 20 | 0 |
| A4/A3B | 0 | 0 | 0 | 0 | 0 | | 0 | | | 0 | 0 | | | 0 | 0 | | 0 | | 0 | 0 | 0 | 0 | 0 | 0 | 0 | 0 | 0 | 0 | | 0 | 0 |
| FA4/FA3B | 0 | 0 | 0 | 38 | 0 | | 0 | | | 1 | 0 | | | 0 | 0 | | 0 | | 0 | 0 | 12 | 0 | 0 | 0 | 0 | 0 | 15 | 3 | | 0 | 0 |
| Unoccupied | 0 | 0 | 0 | 0 | 0 | 0 | 0 | 0 | 0 | 0 | 0 | 0 | 0 | 0 | 0 | 0 | 0 | 0 | 0 | 0 | 1 | 0 | 0 | 0 | 0 | 0 | 0 | 0 | 0 | 0 | 1 |

| gp140.664.R4 | N88 | N130 | N139 | N142 | N148 | N156 | N160 | N186 | N189 | N197 | N230 | N234 | N241 | N262 | N276 | N289 | N295 | N301 | N332 | N339 | N355 | N392 | N406 | N412 | N442 | N448 | N463 | N611 | N618 | N625 | N637 |
|---|---|---|---|---|---|---|---|---|---|---|---|---|---|---|---|---|---|---|---|---|---|---|---|---|---|---|---|---|---|---|---|
| High mannose | 99 | 99 | 34 | 0 | 100 | n.d. | 98 | n.d. | n.d. | 96 | 100 | n.d. | n.d. | 100 | 100 | n.d. | 100 | 100 | 100 | 100 | 21 | 100 | 0 | 100 | 100 | 100 | 10 | 67 | 0 | 51 | 99 |
| M9 | 0 | 47 | 0 | 0 | 0 | | 38 | | | 38 | 88 | | | 87 | 15 | | 100 | 86 | 93 | 36 | 0 | 0 | 0 | 100 | 60 | 77 | 0 | 0 | 0 | 0 | 11 |
| M8 | 23 | 45 | 0 | 0 | 0 | | 46 | | | 42 | 12 | | | 13 | 75 | | 0 | 14 | 6 | 49 | 1 | 82 | 0 | 0 | 32 | 20 | 0 | 0 | 0 | 2 | 34 |
| M7 | 24 | 4 | 7 | 0 | 68 | | 7 | | | 8 | 0 | | | 0 | 8 | | 0 | 0 | 0 | 12 | 2 | 18 | 0 | 0 | 7 | 3 | 1 | 5 | 0 | 11 | 31 |
| M6 | 31 | 2 | 5 | 0 | 0 | | 4 | | | 4 | 0 | | | 0 | 2 | | 0 | 0 | 0 | 3 | 3 | 0 | 0 | 0 | 0 | 0 | 0 | 4 | 0 | 8 | 12 |
| M5 | 15 | 1 | 21 | 0 | 32 | | 3 | | | 4 | 0 | | | 0 | 0 | | 0 | 0 | 0 | 0 | 13 | 0 | 0 | 0 | 0 | 0 | 7 | 59 | 0 | 25 | 7 |
| M4 | 4 | 0 | 2 | 0 | 0 | | 1 | | | 0 | 0 | | | 0 | 0 | | 0 | 0 | 0 | 0 | 1 | 0 | 0 | 0 | 0 | 0 | 1 | 0 | 0 | 0 | 0 |
| M3 | 0 | 0 | 0 | 0 | 0 | | 0 | | | 0 | 0 | | | 0 | 0 | | 0 | 0 | 0 | 0 | 0 | 0 | 0 | 0 | 0 | 0 | 1 | 0 | 0 | 0 | 0 |
| Hybrid | 1 | 0 | 0 | 0 | 0 | | 0 | | | 0 | 0 | | | 0 | 0 | | 0 | 0 | 0 | 0 | 0 | 0 | 0 | 0 | 0 | 0 | 0 | 0 | 0 | 4 | 3 |
| Fhybrid | 0 | 0 | 0 | 0 | 0 | | 0 | | | 0 | 0 | | | 0 | 0 | | 0 | 0 | 0 | 0 | 1 | 0 | 0 | 0 | 0 | 0 | 0 | 0 | 0 | 1 | 0 |
| A1 | 1 | 0 | 1 | 0 | 0 | | 1 | | | 0 | 0 | | | 0 | 0 | | 0 | 0 | 0 | 0 | 1 | 0 | 0 | 0 | 0 | 0 | 0 | 2 | 0 | 5 | 0 |
| FA1 | 0 | 0 | 7 | 0 | 0 | | 0 | | | 0 | 0 | | | 0 | 0 | | 0 | 0 | 0 | 0 | 7 | 0 | 0 | 0 | 0 | 0 | 4 | 4 | 0 | 4 | 0 |
| A2/A1B | 0 | 0 | 0 | 0 | 0 | | 0 | | | 0 | 0 | | | 0 | 0 | | 0 | 0 | 0 | 0 | 0 | 0 | 0 | 0 | 0 | 0 | 0 | 0 | 0 | 10 | 0 |
| FA2/FA1B | 0 | 0 | 14 | 33 | 0 | n.d. | 0 | n.d. | n.d. | 1 | 0 | n.d. | n.d. | 0 | 0 | n.d. | 0 | 0 | 0 | 0 | 18 | 0 | 100 | 0 | 0 | 0 | 25 | 9 | 51 | 19 | 1 |
| A3/A2B | 0 | 0 | 0 | 0 | 0 | | 0 | | | 0 | 0 | | | 0 | 0 | | 0 | 0 | 0 | 0 | 0 | 0 | 0 | 0 | 0 | 0 | 0 | 0 | 0 | 1 | 0 |
| FA3/FA2B | 0 | 0 | 45 | 37 | 0 | | 0 | | | 1 | 0 | | | 0 | 0 | | 0 | 0 | 0 | 0 | 42 | 0 | 0 | 0 | 0 | 0 | 52 | 18 | 49 | 10 | 0 |
| A4/A3B | 0 | 0 | 0 | 0 | 0 | | 0 | | | 0 | 0 | | | 0 | 0 | | 0 | 0 | 0 | 0 | 0 | 0 | 0 | 0 | 0 | 0 | 0 | 0 | 0 | 0 | 0 |
| FA4/FA3B | 0 | 0 | 0 | 30 | 0 | | 0 | | | 0 | 0 | | | 0 | 0 | | 0 | 0 | 0 | 0 | 7 | 0 | 0 | 0 | 0 | 0 | 8 | 0 | 0 | 0 | 0 |
| Unoccupied | 0 | 0 | 0 | 0 | 0 | 0 | 0 | 0 | 0 | 1 | 0 | 0 | 0 | 0 | 0 | 0 | 0 | 0 | 0 | 0 | 3 | 0 | 0 | 0 | 0 | 0 | 0 | 0 | 0 | 0 | 0 |

| UFO.664 | N88 | N130 | N139 | N142 | N148 | N156 | N160 | N186 | N189 | N197 | N230 | N234 | N241 | N262 | N276 | N289 | N295 | N301 | N332 | N339 | N355 | N392 | N406 | N412 | N442 | N448 | N463 | N611 | N618 | N625 | N637 |
|---|---|---|---|---|---|---|---|---|---|---|---|---|---|---|---|---|---|---|---|---|---|---|---|---|---|---|---|---|---|---|---|
| High mannose | 88 | 100 | 23 | 0 | 100 | 100 | 100 | n.d | n.d. | 99 | 100 | 100 | 100 | 100 | 100 | n.d. | 100 | 100 | 100 | 100 | 11 | 100 | 0 | 100 | 100 | 100 | 4 | 24 | 0 | 0 | 84 |
| M9 | 0 | 70 | 0 | 0 | 0 | 100 | 60 | | | 58 | 83 | 70 | 59 | 89 | 20 | | 100 | 71 | 94 | 42 | 0 | 28 | 0 | 100 | 78 | 75 | 0 | 0 | 0 | 0 | 0 |
| M8 | 8 | 22 | 0 | 0 | 0 | 0 | 35 | | | 33 | 17 | 30 | 41 | 10 | 73 | | 0 | 29 | 5 | 46 | 0 | 44 | 0 | 0 | 19 | 22 | 0 | 0 | 0 | 0 | 4 |
| M7 | 12 | 4 | 0 | 0 | 25 | 0 | 5 | | | 4 | 0 | 0 | 0 | 1 | 6 | | 0 | 0 | 1 | 10 | 0 | 29 | 0 | 0 | 2 | 3 | 0 | 0 | 0 | 0 | 38 |
| M6 | 21 | 1 | 0 | 0 | 0 | 0 | 0 | | | 2 | 0 | 0 | 0 | 0 | 1 | | 0 | 0 | 0 | 2 | 0 | 0 | 0 | 0 | 0 | 1 | 0 | 0 | 0 | 0 | 17 |
| M5 | 31 | 2 | 23 | 0 | 75 | 0 | 0 | | | 2 | 0 | 0 | 0 | 0 | 0 | | 0 | 0 | 0 | 0 | 8 | 0 | 0 | 0 | 0 | 0 | 3 | 20 | 0 | 0 | 16 |
| M4 | 7 | 0 | 0 | 0 | 0 | 0 | 0 | | | 0 | 0 | 0 | 0 | 0 | 0 | | 0 | 0 | 0 | 0 | 0 | 0 | 0 | 0 | 0 | 0 | 0 | 0 | 0 | 0 | 1 |
| M3 | 1 | 0 | 0 | 0 | 0 | 0 | 0 | | | 0 | 0 | 0 | 0 | 0 | 0 | | 0 | 0 | 0 | 0 | 0 | 0 | 0 | 0 | 0 | 0 | 0 | 0 | 0 | 0 | 0 |
| Hybrid | 7 | 0 | 0 | 0 | 0 | 0 | 0 | | | 0 | 0 | 0 | 0 | 0 | 0 | | 0 | 0 | 0 | 0 | 0 | 0 | 0 | 0 | 0 | 0 | 0 | 2 | 0 | 0 | 4 |
| Fhybrid | 1 | 0 | 0 | 0 | 0 | 0 | 0 | | | 0 | 0 | 0 | 0 | 0 | 0 | | 0 | 0 | 0 | 0 | 1 | 0 | 0 | 0 | 0 | 0 | 0 | 1 | 0 | 0 | 3 |
| A1 | 5 | 0 | 0 | 0 | 0 | 0 | 0 | | | 0 | 0 | 0 | 0 | 0 | 0 | | 0 | 0 | 0 | 0 | 1 | 0 | 0 | 0 | 0 | 0 | 2 | 2 | 0 | 0 | 1 |
| FA1 | 0 | 0 | 12 | 0 | 0 | 0 | 0 | | | 0 | 0 | 0 | 0 | 0 | 0 | | 0 | 0 | 0 | 0 | 6 | 0 | 0 | 0 | 0 | 0 | 5 | 6 | 1 | 0 | 5 |
| A2/A1B | 5 | 0 | 0 | 0 | 0 | 0 | 0 | | | 0 | 0 | 0 | 0 | 0 | 0 | | 0 | 0 | 0 | 0 | 0 | 0 | 0 | 0 | 0 | 0 | 8 | 0 | 0 | 0 | 0 |
| FA2/FA1B | 1 | 0 | 17 | 40 | 0 | 0 | 0 | n.d | n.d. | 0 | 0 | 0 | 0 | 0 | 0 | n .d. | 0 | 0 | 0 | 0 | 19 | 0 | 41 | 0 | 0 | 0 | 26 | 11 | 34 | 9 | 8 |
| A3/A2B | 1 | 0 | 0 | 0 | 0 | 0 | 0 | | | 0 | 0 | 0 | 0 | 0 | 0 | | 0 | 0 | 0 | 0 | 0 | 0 | 0 | 0 | 0 | 0 | 1 | 0 | 0 | 0 | 0 |
| FA3/FA2B | 1 | 0 | 49 | 44 | 0 | 0 | 0 | | | 0 | 0 | 0 | 0 | 0 | 0 | | 0 | 0 | 0 | 0 | 54 | 0 | 58 | 0 | 0 | 0 | 45 | 43 | 64 | 79 | 0 |
| A4/A3B | 0 | 0 | 0 | 0 | 0 | 0 | 0 | | | 0 | 0 | 0 | 0 | 0 | 0 | | 0 | 0 | 0 | 0 | 0 | 0 | 0 | 0 | 0 | 0 | 0 | 0 | 0 | 0 | 0 |
| FA4/FA3B | 0 | 0 | 0 | 16 | 0 | 0 | 0 | | | 0 | 0 | 0 | 0 | 0 | 0 | | 0 | 0 | 0 | 0 | 9 | 0 | 0 | 0 | 0 | 0 | 8 | 4 | 0 | 12 | 0 |
| Unoccupied | 0 | 0 | 0 | 0 | 0 | 0 | 0 | 0 | 0 | 1 | 0 | 0 | 0 | 0 | 0 | 0 | 0 | 0 | 0 | 0 | 0 | 0 | 0 | 0 | 0 | 0 | 0 | 9 | 0 | 0 | 2 |

**fig. S8. Site-specific *N*-linked glycan analysis of Du172.17 SOSIP.664, HR1-redesigned (gp140.664.R4), and UFO.664 trimers produced in HEK293F cells.** Quantification of site-specific glycan occupancy and composition. The table shows the compositions found at each site. Compositions corresponding to oligomannose/hybrid-type glycans are colored in green and fully processed complex type glycans are colored in magenta. The proportion of peptides at each lacking an attached glycan are colored in grey. Oligomannose-type glycans are categorized according to the number of mannose residues, hybrid-type glycans according to the presence/absence of fucose, and complex-type glycans according to the number of processed antenna and the presence/absence of fucose.

### Figure S9

A Group 1 rabbit plasma neutralization against MLV, SF162 and BG505.T332N at –d10 and w30

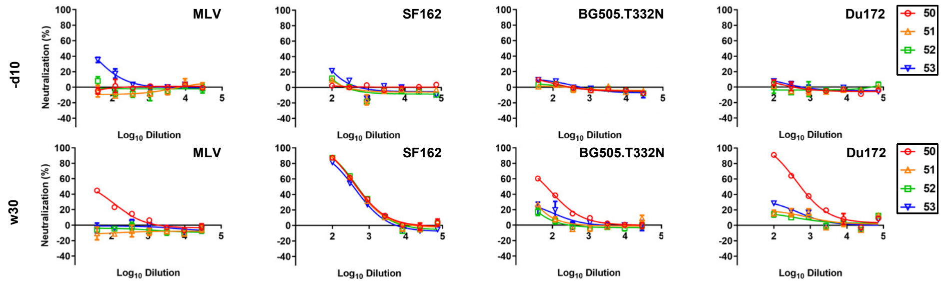


B Group 1 rabbit plasma neutralization against the 12-virus global panel at –d10 and w30

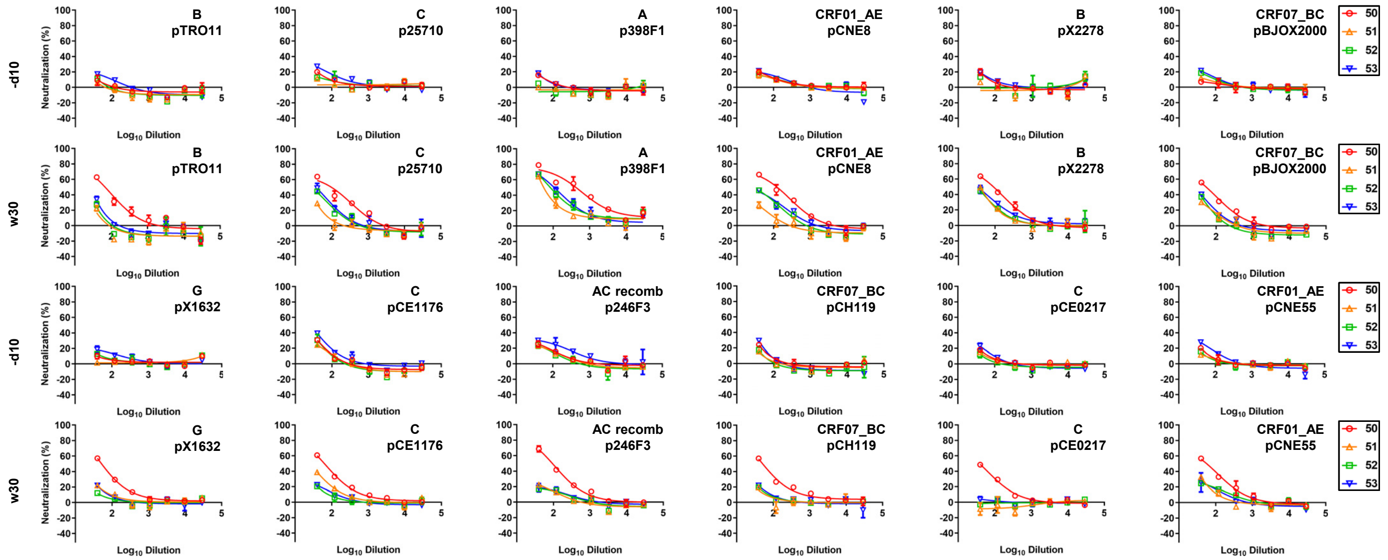


C Group 2 rabbit plasma neutralization against MLV, SF162 and BG505.T332N at –d10 and w30

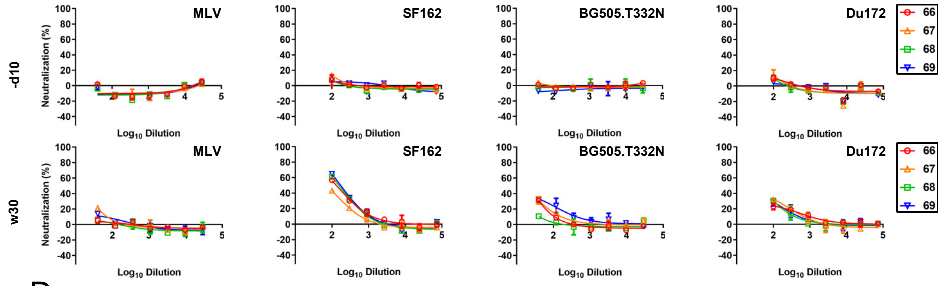


D Group 2 rabbit plasma neutralization against the 12-virus global panel at –d10 and w30

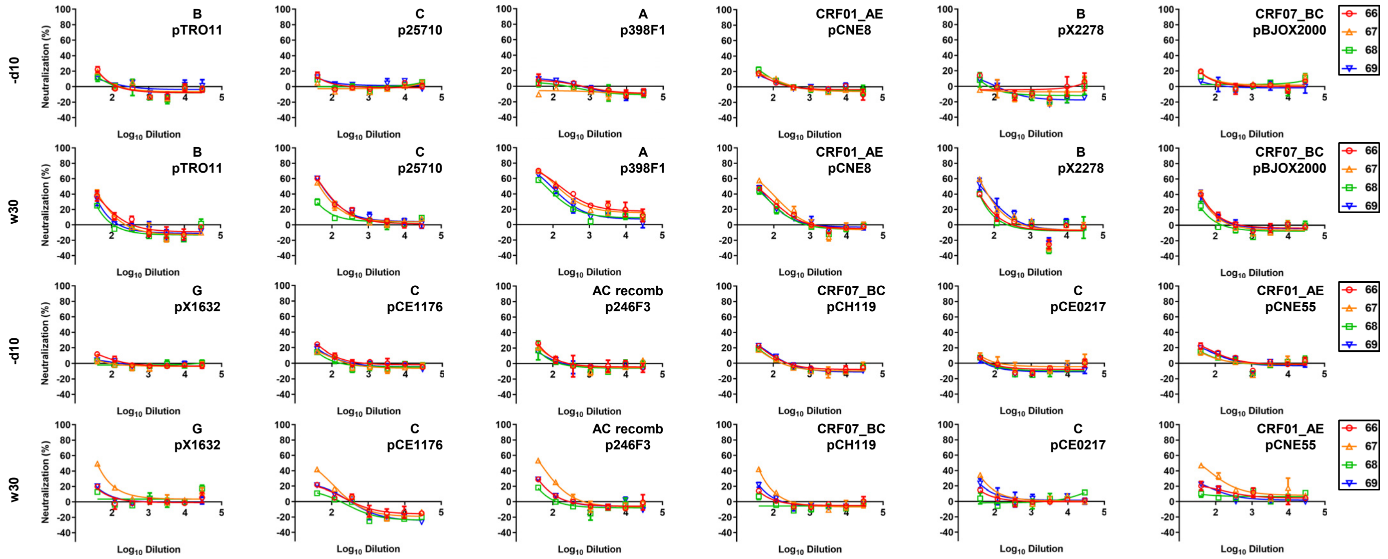

**Figure S9**

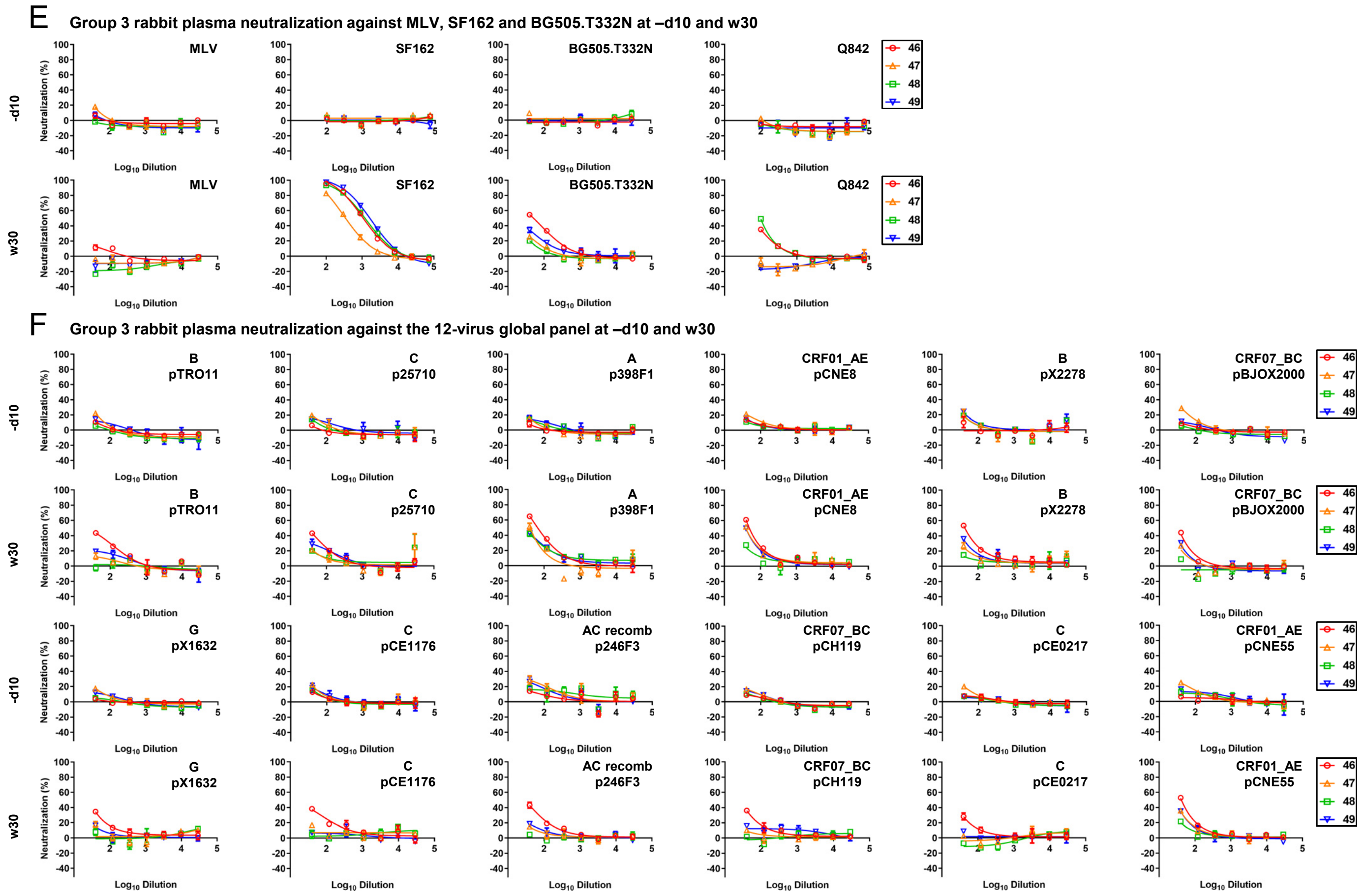


**fig. S9. Rabbit plasma neutralization from three non-BG505 Env-immunized rabbit groups.** Three groups of rabbits were immunized with Du172.17 UFO-BG trimer, gp140.664.R4-FR nanoparticle, and Q842-d12 UFO-BG trimer. (**A**) Neutralization of MLV, tier 1 clade B SF162, tier 2 clade A BG505.T332N, and tier 2 clade C Du172.17 by day -10 (-d10) and week 30 (w30) rabbit plasma from the Du172.17 trimer group. (**B**) Neutralization of all 12 isolates from a global panel by day -10 (-d10) and week 30 (w30) rabbit plasma from the Du172.17 trimer group. (**C**) Neutralization of MLV, SF162, BG505.T332N, and Du172.17 by day -10 (-d10) and week 30 (w30) rabbit plasma from the Du172.17 ferritin nanoparticle group. (**D**) Neutralization of all 12 isolates from a global panel by day -10 (-d10) and week 30 (w30) rabbit plasma from the Du172.17 ferritin nanoparticle group. (**E**) Neutralization of MLV, SF162, BG505.T332N, and tier 2 clade A Q842-d12 by day -10 (-d10) and week 30 (w30) rabbit plasma from the Q842-d12 trimer group. (**F**) Neutralization of all 12 isolates from a global panel by day -10 (-d10) and week 30 (w30) rabbit plasma from the Q842-d12 trimer group. The heat-inactivated plasma was diluted 100-fold for autologous virus and tier 1 SF162 and subjected to a 3-fold dilution series in the TZM-bl assay. To increase the sensitivity of detection, heat-inactivated plasma was diluted 40-fold for MLV and all other heterologous tier 2 isolates and followed by a 3-fold dilution series in the TZM-bl assay. $ID_{50}$ titers for plots (A) – (F) are summarized in Fig. 4D.

**Figure S10**

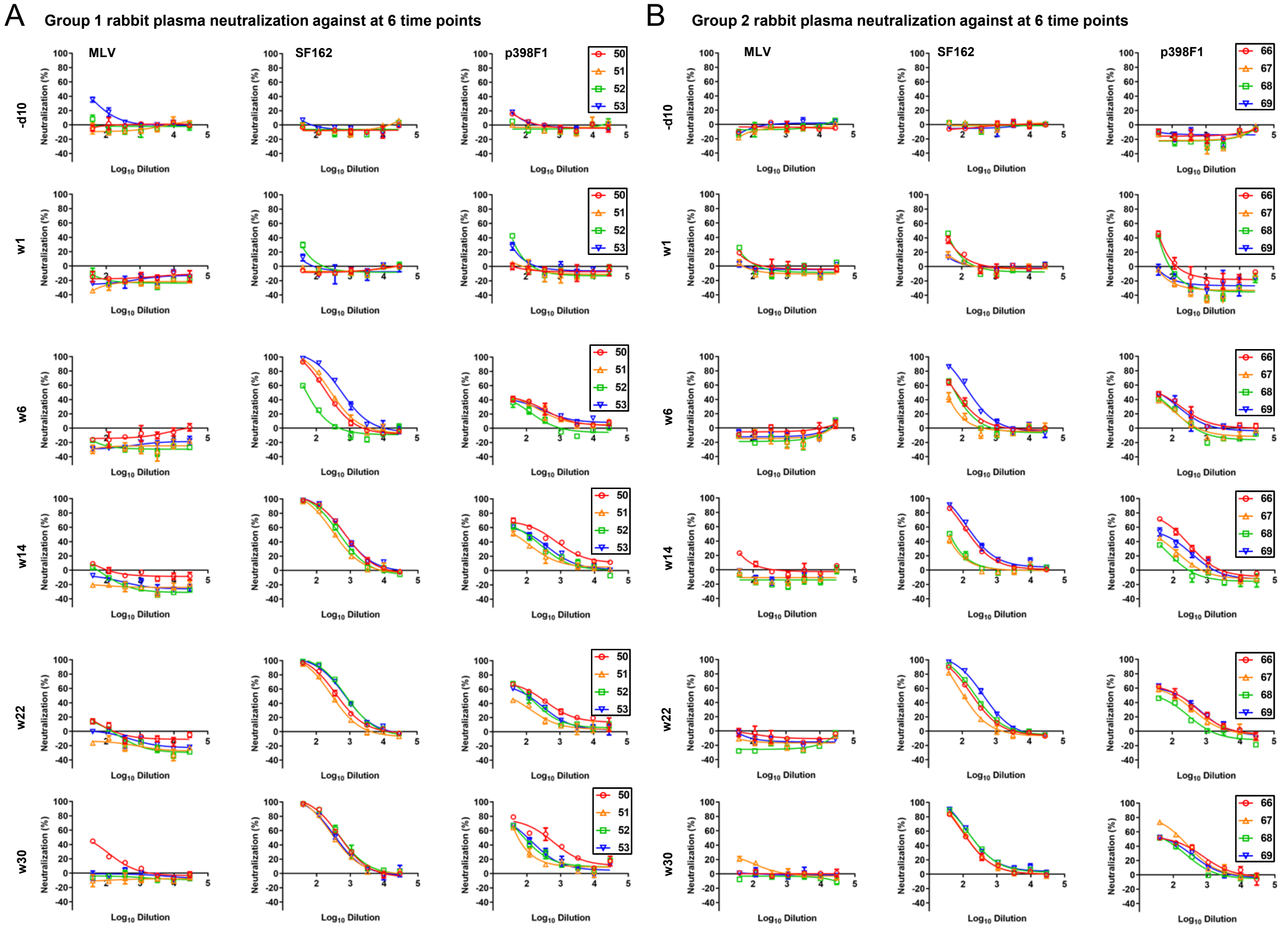


**fig. S10. Longitudinal rabbit plasma neutralization from two clade C Du172.17 Env-immunized rabbit groups.** Two rabbit groups immunized with Du172.17 UFO-BG trimer and gp140.664.R4-FR nanoparticle were analyzed. (**A**) Neutralization of MLV, tier 1 clade B SF162, and tier 2 clade A p398F1 by day -10 (-d10) and weeks 1, 6, 14, 22, and 30 rabbit plasma from the Du172.17 trimer group. **(B)** Neutralization of MLV, tier 1 clade B SF162, and tier 2 clade A p398F1 by day -10 (-d10) and weeks 1, 6, 14, 22, and 30 rabbit plasma from the Du172.17 gp140.664.R4-FR nanoparticle group. In this analysis, the heat-inactivated plasma was diluted 40-fold for both SF162 and p398F1 and then subjected to a 3-fold dilution series in the TZM-bl assay. $ID_{50}$ titers for plots (A) – (B) are shown in Fig. 4E.

**Table S1. X-ray crystallographic data collection and refinement statistics.**

| **Data collection** | BG505 gp120 core, Fabs M4H2K1, 17b | Fab M4H2K1 | Du172.17 gp140.664.R4, Fabs PGT124, 35O22 |
|---|---|---|---|
| X-ray Source | APS 23ID-B | APS 23ID-D | APS 23ID-D |
| Wavelength (Å) | 1.033 | 1.033 | 1.033 |
| Detector | Eiger | Pilatus | Pilatus |
| Space group | $P2_12_12$ | $P3_121$ | $P6_3$ |
| Unit cell parameters | a = 204.0, b = 60.6, c = 166.7 Å | a = b = 68.3, c = 184.7 Å | a = b = 127.0, c = 316.5 Å |
| Resolution (Å) | 50.00 – 4.30 (4.73 – 4.63) (4.63 – 4.54) (4.54 – 4.45) (4.45 – 4.37) (4.37 – 4.30) | 50.00 – 1.50 (1.53 – 1.50)[a] | 50.00 – 3.40 (3.46 – 3.40)[a] |
| Observations | 109,015 | 929,037 | 248,831 |
| Unique reflections | 12,843 (280)[a] | 81,206 (3,998)[a] | 38,788 (1614)[a] |
| Redundancy | 8.5 (1.8)[a] | 11.4 (11.1)[a] | 6.4 (3.7)[a] |
| Completeness (%) | 86.8 (80.4) (67.2) (61.8) (52.6) (45.6) (39.4) | 99.8 (99.5)[a] | 97.6 (81.1) |
| $<I/\sigma_I>$[b] | 13.0 (1.0)[a] | 34.9 (2.0)[a] | 8.6 (1.0)[a] |
| $R_{sym}$[c] | 0.21 (0.83)[a] | 0.08 (0.99)[a] | 0.21 (1.00)[a] |
| $R_{pim}$[c] | 0.06 (0.50)[a] | 0.02 (0.29)[a] | 0.08 (0.48)[a] |
| $CC_{1/2}$ | 0.86 (0.39)[a] | 0.94 (0.73)[a] | 0.85 (0.45)[a] |
| **Refinement statistics** | | | |
| Resolution (Å) | 43.04 – 4.30 | 49.88 – 1.50 | 49.53 – 3.40 |
| Reflections (work) | 12,392 | 81,150 | 38,200 |
| $R_{cryst}$ (%)[d] / $R_{free}$ (%)[e] | 30.1 / 33.3 | 18.3 / 21.6 | 30.3 / 32.6 |
| No. atoms | | | |
| Protein / Ligands | 9429 | 3347 / 17 | 11322 |
| Glycan | 282 | - | 726 |
| Water | - | 624 | - |
| Average *B*-value (Å$^2$) | | | |
| Protein | 172 | 24 | 108 |
| Glycan | 89 | - | 147 |
| Water | - | 36 | - |
| Wilson *B*-value (Å$^2$) | 139 | 18 | 93 |
| RMSD from ideal geometry | | | |
| Bond length (Å) | 0.004 | 0.009 | 0.002 |
| Bond angles (°) | 0.85 | 1.16 | 0.50 |
| Ramachandran statistics (%)[f] | | | |
| Favored | 95.05 | 97.71 | 90.58 |
| Allowed | 4.37 | 2.29 | 8.46 |
| Outliers | 0.58 | 0 | 0.96 |
| PDB ID | 7KLC | 7KKZ | 7KMD |

[a] Numbers in parentheses refer to the highest resolution shell.

[b] Calculated as average($I$)/average($\sigma I$)

[c] $R_{sym} = \Sigma_{hkl}\Sigma_i \mid I_{hkl,i} - \langle I_{hkl}\rangle \mid / \Sigma_{hkl}\Sigma_i I_{hkl,I}$, where $I_{hkl,i}$ is the scaled intensity of the $i^{th}$ measurement of reflection h, k, l, $\langle I_{hkl}\rangle$ is the average intensity for that reflection, and $n$ is the redundancy. $R_{pim}$ is a redundancy-independent measure of the quality of intensity measurements. $R_{pim} = \Sigma_{hkl} (1/(n-1))^{1/2} \Sigma_i \mid I_{hkl,i} - \langle I_{hkl}\rangle \mid / \Sigma_{hkl} \Sigma_i I_{hkl,I}$, where $I_{hkl,i}$ is the scaled intensity of the $i^{th}$ measurement of reflection h, k, l, $\langle I_{hkl} \rangle$ is the average intensity for that reflection, and $n$ is the redundancy.

[d] $R_{cryst} = \Sigma_{hkl} \mid F_o - F_c \mid / \Sigma_{hkl} \mid F_o \mid \times 100$

[e] $R_{free}$ was calculated as for $R_{cryst}$, but on a test set comprising 5% of the data excluded from refinement.

[f] These values were calculated using MolProbity (http://molprobity.biochem.duke.edu/).
